# Albumin-binding RNAi Conjugate for Carrier Free Treatment of Arthritis

**DOI:** 10.1101/2023.05.31.542971

**Authors:** Juan M. Colazo, Ella N. Hoogenboezem, Megan C. Keech, Nora Francini, Veeraj Shah, Fang Yu, Justin H. Lo, Alexander G. Sorets, Joshua T. McCune, Hongsik Cho, Carlisle R. DeJulius, Danielle L. Michell, Tristan Maerz, Kacey C. Vickers, Katherine N. Gibson-Corley, Karen A. Hasty, Leslie Crofford, Rebecca S. Cook, Craig L. Duvall

**Author notes:** **Corresponding author: Craig L. Duvall, Ph.D.** Cornelius Vanderbilt Professor of Biomedical Engineering, Vanderbilt University, **email:**, **mailing address:** 5824 Stevenson Center, Nashville, TN 37232.

## Abstract

Osteoarthritis (OA) and rheumatoid arthritis (RA) are joint diseases that are associated with pain and lost quality of life. No disease modifying OA drugs are currently available. RA treatments are better established but are not always effective and can cause immune suppression. Here, an MMP13-selective siRNA conjugate was developed that, when delivered intravenously, docks onto endogenous albumin and promotes preferential accumulation in articular cartilage and synovia of OA and RA joints. MMP13 expression was diminished upon intravenous delivery of MMP13 siRNA conjugates, consequently decreasing multiple histological and molecular markers of disease severity, while also reducing clinical manifestations such as swelling (RA) and joint pressure sensitivity (RA and OA). Importantly, MMP13 silencing provided more comprehensive OA treatment efficacy than standard of care (steroids) or experimental MMP inhibitors. These data demonstrate the utility of albumin ‘hitchhiking’ for drug delivery to arthritic joints, and establish the therapeutic utility of systemically delivered anti-MMP13 siRNA conjugates in OA and RA.

**Editorial summary:** Lipophilic siRNA conjugates optimized for albumin binding and “hitchhiking” can be leveraged to achieve preferential delivery to and gene silencing activity within arthritic joints. Chemical stabilization of the lipophilic siRNA enables intravenous siRNA delivery without lipid or polymer encapsulation. Using siRNA sequences targeting MMP13, a key driver of arthritis-related inflammation, albumin hitchhiking siRNA diminished MMP13, inflammation, and manifestations of osteoarthritis and rheumatoid arthritis at molecular, histological, and clinical levels, consistently outperforming clinical standards of care and small molecule MMP antagonists.

## Introduction

Osteoarthritis (OA) is a degenerative joint disease that can be idiopathic or secondary to injury, in the case of post-traumatic osteoarthritis (PTOA). OA is characterized by cartilage loss, synovial inflammation, and formation of osteophytes, all of which contribute to pain and lost joint function^1^. Patients often develop broader multi-joint osteoarthritis (MJOA), which can have debilitating impact on quality of life^2^. The autoimmune disease rheumatoid arthritis (RA) has independent etiology and is characterized by severe synovial and systemic inflammation, bone erosion, and cartilage damage^3,4^. Current OA treatment options are palliative and include non-steroidal anti-inflammatory drugs (NSAIDS) or corticosteroids, approaches that do not counteract the root causes of disease nor prevent OA progression; corticosteroids may in fact worsen cartilage thinning^5–7^. Current treatments for RA include synthetic disease-modifying antirheumatic drugs (DMARDs) and biologic DMARDs such as tumor necrosis factor (TNF) inhibitors, interleukin (IL) inhibitors, and B-cell inhibitors. These treatments are efficacious, although they can suppress the immune system, and their benefits often wane over the long-term^8^.

There is extensive overlap between OA and RA in terms of the downstream molecular effectors that drive joint degeneration^9^. Both are associated with inflammation and upregulation of extracellular matrix (ECM) degrading matrix metalloproteinases (MMPs)^10^. MMPs erode cartilage, and the degradation byproducts have inflammatory signaling properties that further induce expression of inflammatory cytokines and MMPs, perpetuating a chronic, catabolic cycle that leads to deep cartilage erosion, chondrocyte loss, and subchondral bone exposure^11–13^.

Therapeutic MMP inhibition is a logical approach for OA and RA treatment. Development of MMP-selective antagonists is critical, as broad spectrum MMP inhibitors cause musculoskeletal toxicities in humans, likely due to universal disruption of normal tissue homeostasis^14–16^. Among MMPs, MMP13 is perhaps the most efficient at cleaving the key ECM components, collagen II^17^ and fibronectin^18^, producing inflammatory fragments. Further, knockout mouse studies pinpoint MMP13 as a key proteolytic driver of cartilage loss in arthritis models^19^. Lastly, MMP13 expression is relatively confined to joint tissues and diseased or injured tissues characterized by fibrosis or inflammation^20^. This narrow expression pattern allows for a potentially wider therapeutic window with minimal toxicities. However, this hypothesis has yet to be tested fully, as clinical development of selective MMP inhibitors has been challenged by the extensive structural similarities between the catalytic sites of the various MMPs^21^.

RNA interference (RNAi) provides an ideal technique for gene-selective inhibition, as specificity is conferred by the genetic sequence. The translatability of siRNA therapeutics is also now well-established based on the FDA approval of lipid nanoparticle and GalNAc-siRNA conjugate drugs, both of which have been successfully applied in humans for gene silencing in the liver^22,23^. Selective MMP13 gene knockdown using siRNA sequences would overcome the hurdle of structural similarity between MMPs, thus providing an opportunity to target MMP13 in the context of OA and RA. This notion is supported by recent PTOA model studies in which intra-articular delivery of MMP13 silencing siRNA nanoparticles (siNPs) protected against knee joint load-induced cartilage loss^24,25^.

To improve upon these relatively-complex technologies for sustained MMP13 siRNA delivery via intra-articular injection of siNPs, we sought an approach in which carrier-free siRNA conjugates delivered intravenously might preferentially accumulate within arthritic joints. A systemically delivered approach avoids the potential joint damage that can occur from repeated intra-articular injections and better enables treatment of MJOA and RA. Relative to siNPs, a molecularly-defined conjugate is also simpler to manufacture and does not come with the added risk of carrier toxicities that often narrow the therapeutic window of cationic lipid or polymer siNP formulations.

Albumin also has natural binding sites for long chain fatty acids^26–31^ that can be co-opted by drugs or fatty acid-modified exogenous molecules, as exemplified by clinically-approved albumin-based/binding formulations (*e.g.*, Abraxane, Semaglutide, Levemir, Optison, and Tirzepatide)^32–34^. Binding to these sites yields the potential for increasing drug pharmacokinetic properties, as albumin accesses natural recycling and kidney re-absorption mechanisms that prevent its degradation and urinary clearance, thus prolonging its half-llfe^30,35–38^. Hitchhiking on albumin also can promote preferential accumulation in arthritic joints because their inflamed state causes leaky vasculature and permits local drug extravasation and sequestration (the “ELVIS” effect^39,40^). Albumin is one of the most prominent proteins in synovial fluid (SF)^41,42^, and it also shows cartilage penetration, being 15-fold more concentrated in the deep layers of arthritic than healthy cartilage and having 2-fold higher penetration into human cartilage than IgG antibodies^43,44^. Recombinant fusion to albumin promotes protein accumulation in inflamed knees^45,46^, further motivating our approach.

Although covalent conjugation of siRNA to albumin has been explored to some extent^47,48^, lipid-siRNA conjugates that enable reversible, in situ association with native circulating albumin has been proof-of-concept tested only in cancer studies^26,49,50^. We show here that systemic delivery of an albumin-hitchhiking siRNA-lipid conjugate targeting MMP13 achieves robust delivery, MMP13 silencing, and therapeutic efficacy in joints afflicted by OA and RA.

## Results and discussion

### Circulating albumin robustly accumulates in PTOA joints

To determine the extent to which circulating albumin accumulates in arthritic joints, we intravenously (i.v.) delivered Cy5-labeled mouse serum albumin (MSA-Cy5) into mice with induced PTOA. In this model, the left knee undergoes repeated loading to induce damage and inflammation, while the right knee remains unchallenged (**Figure 1A**). At 24 hrs after delivery, abundant MSA-Cy5 was observed in the synovium and cartilage of PTOA knees but not in contralateral control knees (**Figure 1B, Supplementary Figure 1**). In contrast, Cy5-conjugated poly(ethylene glycol) (PEG-Cy5) did not accumulate preferentially in PTOA knees (**Figure 1C**). Further, i.v. delivery of the albumin binding dye Evans Blue resulted in preferential accumulation in synovia and cartilage of PTOA knees, but not contralateral healthy knees (**Figure 1D,E, Supplementary Figures 2,3**). Interestingly, mRNAs of genes encoding the albumin delivery / transport factors caveolin-1 and secreted protein acidic and rich in cysteine (SPARC) were elevated in PTOA joint tissues (**Supplementary Figure 3D**), consistent with previous reports of SPARC upregulation in arthritic joints^51–54^. Interestingly, interstitial SPARC expression in tumors correlates with efficacy of albumin-based therapeutics in head and neck cancers^55^. These findings support the idea that hitchhiking on endogenous circulating albumin might be used to skew the biodistribution of siRNA conjugates to arthritic joints following i.v. delivery.

**Figure 1.**
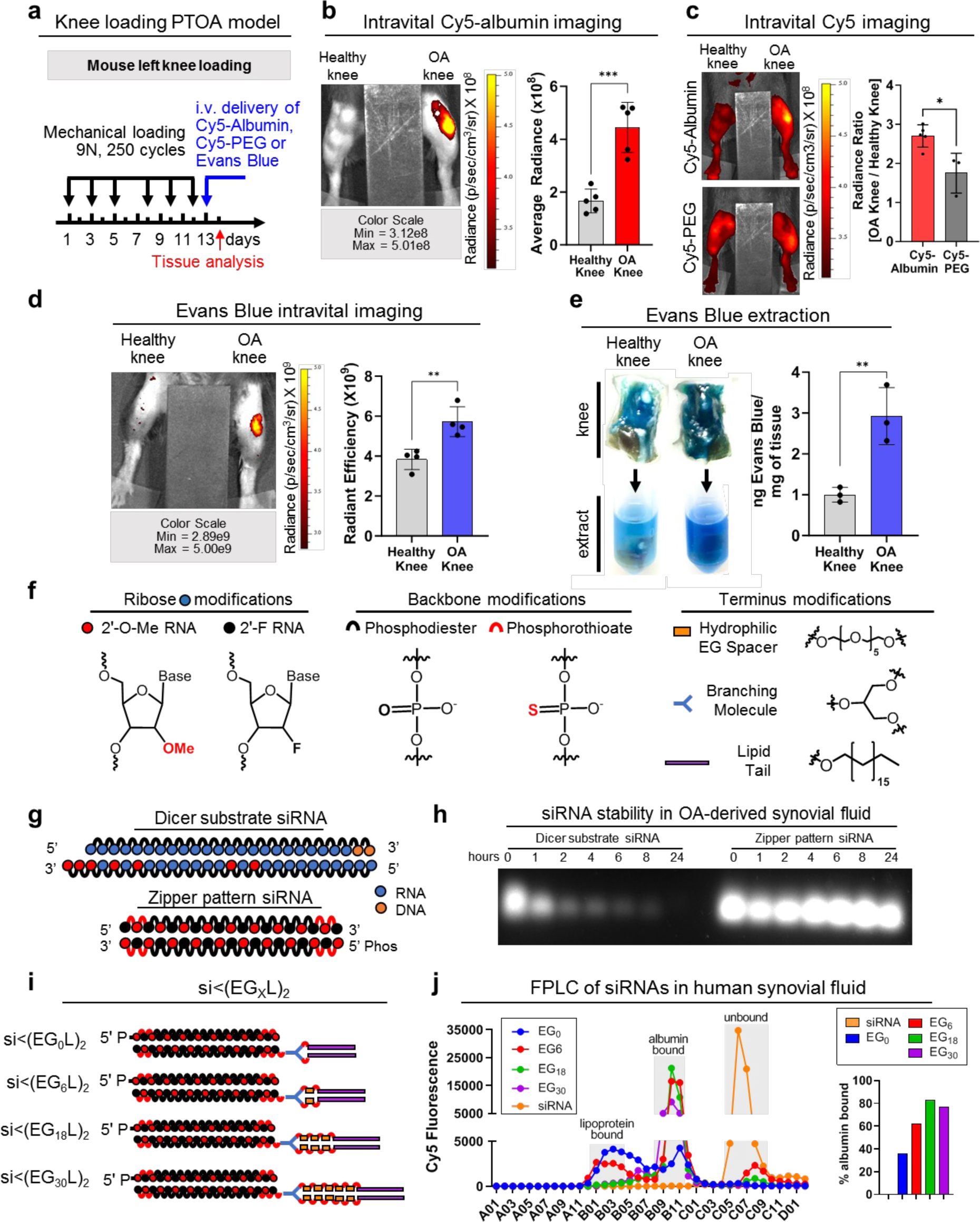
Lipid conjugated siRNA constructs for albumin mediated delivery to PTOA joints. **A-D)** Left knee mechanical loading was used to induce unilateral PTOA in mice, followed by i.v. delivery of MSA-Cy5 (N = 5), PEG-Cy5 (N = 3), and Evans Blue (N = 4). Cy5 and Evans Blue were measured by fluorescence imaging 24 hrs after delivery. Representative images are shown. **A)** Procedural time is shown. **B)** MSA-Cy5 fluorescence in paired PTOA and healthy knees was measured. **C)** Ratio of MSA-Cy5 and PEG-Cy5 fluorescence in PTOA / healthy knees for each treated mouse was measured. **D)** Evans Blue fluorescence in paired contralateral PTOA and healthy knees was measured. **E)** Contralateral PTOA and healthy knees were excised 24 hours after Evans Blue delivery. Evans Blue was extracted and measured by densitometry (N = 3). Representative tissues and resulting extract is shown. **F)** Chemical modifications of siRNA ribose, backbone, and terminus. **G)** Schematic of synthetic dicer substrate siRNA and alternating 2ʹF and 2ʹOMe modified “zipper” siRNA. **H)** Representative gel electrophoresis of dicer substrate and zipper siRNA sequences after incubation in synovial fluid collected from treatment-naïve OA patient. **I)** Schematic representation of the si<(EG_X_L)_2_ series of zipper-modified siRNAs with divalent lipid end-modifications with variations in EG content. **J)** Chromatograph (left panel) of elution of si<(EG_X_L)_2_ variants pre-incubated with healthy human synovial fluid. The percentage of total si<(EG_X_L)_2_ bound to albumin fractions is shown (right panel). *P < 0.05, **P < 0.01, ***P < 0.001.

### Enhanced stability, albumin binding, and PTOA joint accumulation of optimized siRNA-lipid conjugates

RNA therapeutics i.v. administered carrier-free would be highly vulnerable to degradation by endo- and exonucleases. To stabilize the siRNA sequence, we engineered blunt-ended 19-mer RNA strands with alternating 2’-OMe and 2’-F ribose modifications in a “zipper” pattern and replaced phosphodiesters with PS linkages for the last two bases of the 5D- and 3D-termini of the sense and anti-sense strands (**Figure 1F,G**). Stability of the zipper-modified siRNA was assessed in synovial fluid from a treatment-naive OA patient, revealing complete siRNA stability through at least 24 hrs at 37°C, whereas a more lightly modified dicer-substrate siRNA was >50% degraded within 1 hr **(Figure 1H)**. Given the stability of the zipper-modified siRNA, all further siRNA modifications were done using this modification pattern.

The zipper siRNAs were end-modified with bivalent C18 lipids intended to promote binding of the siRNA into the natural fatty acid (FA) binding pockets of albumin (**Figure 1I**)^26^. A library of bivalent lipid structures were synthesized as recently reported^49^. Briefly, a splitter phosphoramidite (<) was added at the 5D end of the sense strand, followed by addition of 0 (EG_0_), 1 (EG_6_), 3 (EG_18_), or 5 (EG_30_) hexa-ethylene glycol (EG_6_) phosphoramidite spacers to each branch. Both branches were terminally appended with a C_18_ lipid, thus generating siRNA<(EG_x_L)_2_, where X is the number of EG repeats linking the splitter to each terminal C18.

Association of the different siRNA conjugates and the parent zipper siRNA structures with synovial fluid albumin was assessed using fast protein liquid chromatography (FPLC). A proportion of each of the siRNA<(EG_X_L)_2_ conjugates was found in the albumin-containing synovial fluid fractions (**Figure 1J**), whereas unconjugated siRNA was not. Notably, >80% of the total siRNA<(EG_18_L)_2_ detected by FPLC was found in the albumin-containing fraction, higher than any other siRNA-lipid conjugate tested. This observation was consistent with the previously-measured high affinity binding of siRNA<(EG_18_L)_2_ to albumin (*K*_D_ = 30 nM)^49^.

Cy5-labeled siRNA<(EG_X_L)_2_ constructs were next delivered i.v. (1 mg/kg) to mice with PTOA induced in the left knee (**Figure 2A**). While each accumulated to a greater extent in PTOA knees over healthy contralateral knees, siRNA<(EG_18_L)_2_ achieved the highest accumulation PTOA joints (**Figure 2B, Supplementary Figure 4**). In contrast to the >4-fold enrichment of siRNA<(EG_18_L)_2_ in PTOA knees over healthy knees, parental siRNA and cholesterol-conjugated siRNA (**siRNA-Chol**) each showed only a 1.5- and 2-fold enrichment to PTOA vs healthy knees (**Figure 2C**). Histological cryosections of PTOA knee synovium and cartilage illustrated holistic joint tissue penetration and cellular uptake of siRNA<(EG_18_L)_2_ (**Supplementary Figure 5**), while siRNA-Chol was observed with greatly reduced intensity in these PTOA joint compartments (**Figure 2D**). Organ biodistribution studies performed by ex vivo Cy5 imaging showed that liver was the main off-target delivery site for both siRNA<(EG_18_L)_2_ and siRNA-Chol, while free siRNA accumulated primarily in kidneys (**Supplementary Figure 6**), a major site of siRNA clearance.

**Figure 2.**
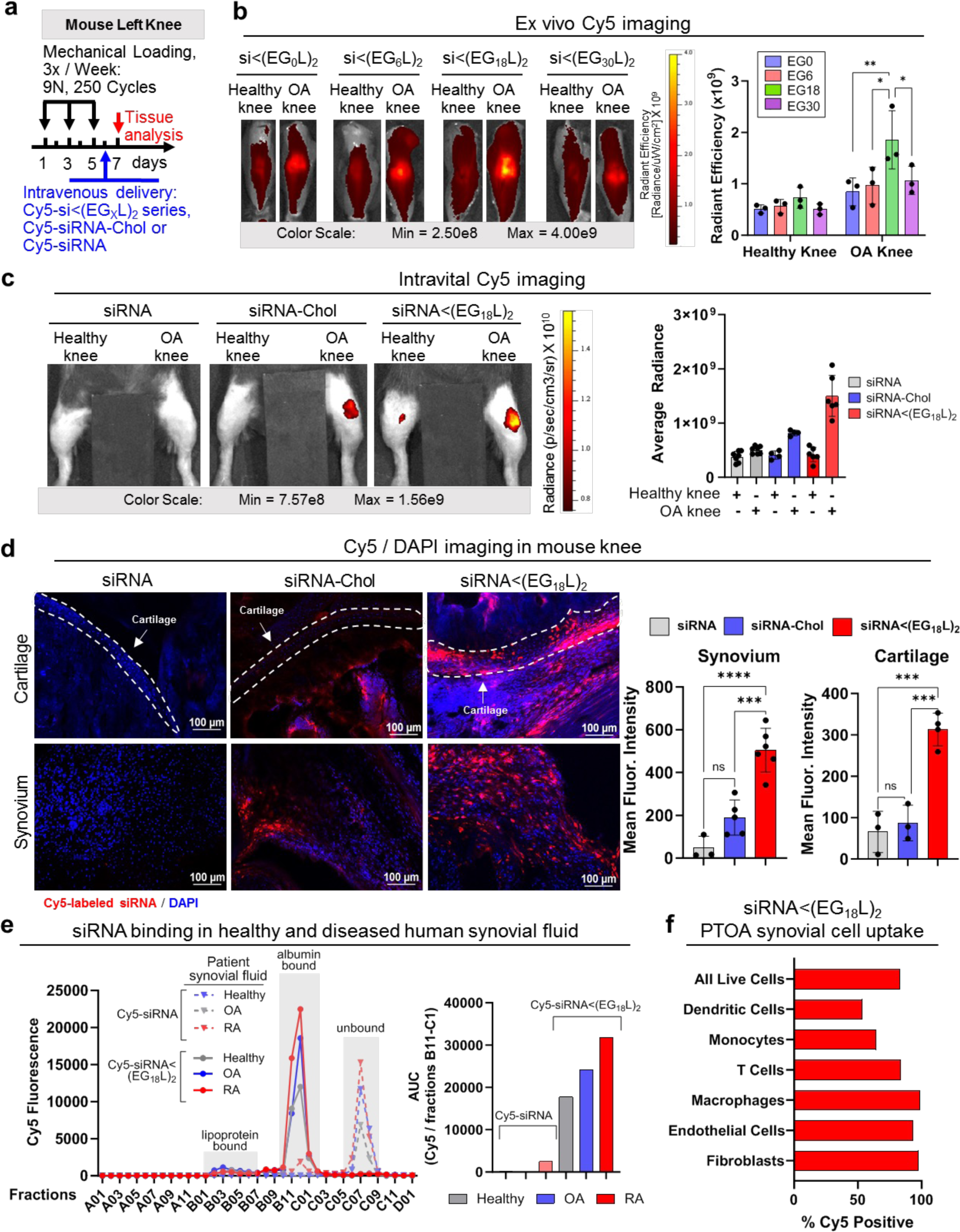
Accumulation of the albumin-binding siRNA-lipid conjugate siRNA<(EG_18_L)_2_ in mouse PTOA knee joints following i.v. delivery. **A)** Unilateral, left knee mechanical loading, treatment, and endpoint protocol used in B-D. **B)** Ex-vivo IVIS images of knee joints and quantification of IVIS signal. **C)** Representative intravital images after 24 hours of Cy5 fluorescence are shown (left panel). Quantification of average Cy5 fluorescence after 24 hours (right panel). Analyzed using mixed effects analysis. **D)** DAPI-counterstained cryosections of knees from treated mice imaged and quantified for Cy5-labeled siRNA fluorescence in cartilage and synovial tissue. **E)** Chromatographic assessment of binding of Cy5-siRNA and Cy5-siRNA<(EG_18_L)_2_ to different components of human synovial fluid taken from a healthy, osteoarthritis, and rheumatoid arthritis patients. Cy5 fluorescence within the albumin-containing fractions was quantitated as area under the curve (AUC). **F)** Flow cytometry assessment cell type specific uptake of Cy5-siRNA<(EG_18_L)_2_ in synovium. Mice were treated i.v. with 10 mg/kg following three bilateral mechanical loading sessions over a week. Cells isolated from synovia of five mice were pooled for analysis. Experimental and flow cytometry gating details are in Supplementary Figure 8. *P < 0.05, **P < 0.01, ***P < 0.001, ****P < 0.0001. Dashed lines outline articular cartilage.

Interactions between albumin and siRNA<(EG_18_L)_2_ were further assessed by FPLC of synovial fluid from OA, RA, or healthy joints. Like what was observed using healthy patient-derived synovial fluid, Cy5-siRNA<(EG_18_L)_2_ eluted primarily in the albumin-containing fractions of OA patient and RA patient-derived synovial fluid samples, but with substantially greater intensity in arthritic samples (**Figure 2E**). In contrast, the parent Cy5-siRNA remained predominantly unbound. Western blotting confirmed the increased albumin content in OA- and RA-derived synovial fractions B12 and C1, the same fractions that siRNA<(EG_18_L)_2_ was predominantly associated with (**Supplementary Figure 7A**). Complementary native gel electrophoresis analyses of synovial fluid fractions B12-C1 revealed colocalization of Cy5-siRNA<(EG_18_L)_2_ with a Coomassie-stained protein of 66 kDa, the known molecular weight of albumin (**Supplementary Figure 7B**). Further, siRNA<(EG_18_L)_2_ showed stability in synovial fluid from OA joints for at least 96 hours (**Supplementary Figure 7C**). Taken together, these findings allowed us to assign siRNA<(EG_18_L)_2_ as our lead siRNA-lipid conjugate for albumin hitchhiking to arthritic joints, motivating a focus on this construct for subsequent therapeutic studies. We also found that siRNA<(EG_18_L)_2_ interacted with albumin across multiple species, including human, mouse, and guinea pig, supporting its utility for all the OA and RA models used herein (**Supplementary Figure 7D**).

To characterize cell type specific uptake of the selected siRNA<(EG_18_L)_2_ conjugate, Cy5-siRNA<(EG_18_L)_2_ (10 mg/kg) or vehicle was delivered i.v. to mice with mechanical load induced PTOA in both knees and cell type specific uptake after 24 hours was characterized using flow cytometry (**Figure 2F, Supplementary Figure 8**). We identified synovial fibroblasts, endothelial cells, macrophages, monocytes, dendritic cells, and T-cells. There was a high total percentage of Cy5 positive cells in the synovium (83.1% of all synovial cells). Fibroblasts, macrophages, and endothelial cells exhibited the highest proportion of Cy5-positive cells.

### siRNA<(EG_18_L)_2_ accumulation in PTOA joints following subcutaneous and intra-articular delivery

Subcutaneous injection of siRNA<(EG_18_L)_2_ was tested as an alternative to the i.v. delivery route, given the utility of subcutaneous injections for patient self-administration of biologic drugs. Subcutaneous delivery of Cy5-siRNA<(EG_18_L)_2_ (2mg/kg) enabled preferential siRNA accumulation in PTOA knees over contralateral unchallenged knees, with similar organ biodistribution profile as i.v. delivery (**Extended Data Fig 1A-C**). However, subcutaneous delivery of siRNA<(EG_18_L)_2_ at 2 mg/kg did not achieve the absolute level of siRNA accumulation in PTOA knees seen with i.v. delivery at 1 mg/kg (**Extended Data Fig 1D**). Notably, a large amount of the Cy5-siRNA<(EG_18_L)_2_ dose was retained at the injection site (data not shown), perhaps due to its rapid interaction with subcutaneous fat or cells at the injection site^56^. Intra-articular (i.a.) delivery, a common clinical route for steroid delivery, was also tested (0.25 mg/kg), revealing greater retention in the PTOA over healthy knee and greater retention than Cy5-siRNA at 48 hrs post-injection, achieving penetration into cartilage and synovia (**Extended Data Fig 1E-G**). While i.a.-delivered Cy5-siRNA<(EG_18_L)_2_ was retained primarily in the knee, Cy5-siRNA redistributed to kidneys, reflecting its relatively rapid removal from the joint by synovial drainage via vasculature and lymphatics, underscoring the importance of albumin-binding moieties for retention of siRNA<(EG_18_L)_2_ within the joint space.

### Potent MMP13 knockdown in PTOA-afflicted joints by siRNA<(EG_18_L)_2_

siRNA sequences against mouse and guinea pig MMP13 (siMMP13) were screened in mouse ATDC5 chondrogenic cells and primary guinea pig chondrocytes, respectively, for MMP13 knockdown potency (**Supplementary Figure 9**). Lead candidate sequences were then synthesized with zipper modifications, enhancing siMMP13 stability in serum (**Supplementary Figure 10**). Transfection of zipper-modified siRNAs achieved potent MMP13 knockdown activity in TNFα-stimulated ATDC5 cells (**Supplementary Figure 10**). Zipper-pattern siMMP13 was further modified with diacyl lipids as described above to enable albumin binding, thus generating siMMP13<(EG_18_L)_2_. Importantly, siMMP13<(EG_18_L)_2_, but not free siMMP13, provided robust carrier-free (no lipofection reagent), *Mmp13* knockdown and efficient cellular uptake in ATDC5 cells (**Supplementary Figure 10, 11**). Cell uptake of siMMP13<(EG_18_L)_2_ was partially diminished in the presence of excess albumin but increased upon treatment with pro-inflammatory TNFα stimulation, even in the presence of excess albumin.

MMP13 silencing by siMMP13<(EG_18_L)_2_ in PTOA-affected mouse knees was first assessed 5 days after i.v. treatment, finding >50% *Mmp13* knockdown with a 5 mg/kg dose, and >60% with a 10 mg/kg dose (**Extended Data Fig 2A**). Neither free siMMP13 nor siMMP13-Chol (each at 10mg/kg, i.v.) significantly affected *Mmp13* levels in PTOA knees. A single intra-articular (i.a.) injection of siMMP13<(EG_18_L)_2_ at 1 mg/kg also decreased *Mmp13* in PTOA knees (**Extended Data Fig 2B**). While subcutaneous delivery of siMMP13<(EG_18_L)_2_ at a relatively high dose (50 mg/kg) diminished *Mmp13* in PTOA knees (**Extended Data Fig 2C**), a lower dose (20 mg/kg) reduced *Mmp13* in PTOA knees only when co-delivered with mouse albumin (1:5 molar ratio). These results show promise for use of siMMP13<(EG_18_L)_2_ with multiple delivery routes. However, subcutaneous delivery requires more development to promote injection site absorption.

Immunohistochemical (**IHC**) analysis of PTOA knees revealed abundant MMP13 in articular cartilage, synovium, and meniscus compared to tissues in healthy knees. PTOA-induced MMP13 was markedly reduced in each PTOA knee compartment at 5 days after treatment with siMMP13<(EG_18_L)_2_ delivered at either 5 or 10 mg/kg. However, neither siMMP13-Chol nor siMMP13 impacted MMP13 levels in PTOA knees (**Extended Data Fig 2D**). MMP13 protein was undetectable in liver or kidneys by IHC (**Extended Data Fig 2E**). RNA expression analyses similarly did not detect *Mmp13* in liver, although low levels were found in kidney, albeit 6000-fold lower than that detected in PTOA knees (**Extended Data Fig 2E**). Regardless, kidney *Mmp13* gene expression was unaffected in siMMP13<(EG_18_L)_2_-treated mice (**Extended Data Fig 2F**), supporting the idea that siMMP13<(EG_18_L)_2_ might have a wide therapeutic window, with minimal risk of molecularly on-target side effects in two of the primary organs commonly associated with siRNA clearance, liver and kidney.

To further characterize cellular sources of MMP13 in PTOA synovium, we analyzed single-cell RNA-sequencing from a murine ACL rupture-induced PTOA model^57^. Synovial fibroblasts, specifically *Thy1+ Pdgfra+ Prg4-* sublining fibroblasts, were the primary *Mmp13-*expressing cell type in synovium, with some myeloid cells exhibiting a much lower degree of *Mmp13* expression. While healthy synovium was essentially devoid of *Mmp13-*expressing cells, PTOA synovium exhibited a large increase in the number of *Mmp13-*positive synovial fibroblasts (**Supplementary Figure 12).** Taken together with Cy5-siRNA<(EG_18_L)_2_ uptake results demonstrating a high degree of uptake by synovial fibroblasts (**Figure 2F, Supplementary Figure 8),** these results demonstrate that our siRNA delivery system efficiently targets the primary *Mmp13*-expressing cells in synovium.

### Sustained stability and bioavailability of siMMP13<(EG_18_L)_2_ enables potent and durable MMP13 knockdown in PTOA joints following systemic administration

Mice were subjected to mechanical left knee loading (3 times / week) for 37 days. They were i.v. treated with a single dose of Cy5-siRNA<(EG_18_L)_2_ at 10mg/kg after the first week of loading, and longitudinal measurements of Cy5 retention in PTOA knees were performed at time points throughout the next 30 days (**Extended Data Fig 3A**). At day 30, remarkable Cy5 signal was retained in cartilage/meniscus and synovial tissues of PTOA knees treated systemically with Cy5-siRNA<(EG_18_L)_2_ (**Extended Data Fig 3B)**. Again, preferential siRNA<(EG_18_L)_2_ accumulation in PTOA knees over contralateral control knees was observed (**Extended Data Fig 3C,D**). Importantly, a single siMMP13<(EG_18_L)_2_ treatment maintained knockdown of MMP13 transcript and protein in PTOA joints through day 30 (**Extended Data Fig 3E,F**). Intra-articular dosing of siRNA<(EG_18_L)_2_ (1mg/kg) was similarly assessed, again revealing preferential retention of Cy5-siRNA<(EG_18_L)_2_ in PTOA knees over contralateral healthy knees through 30 days post-treatment (**Extended Data Fig 4**).

In contrast to the sustained siRNA<(EG_18_L)_2_ retention in PTOA joints, minimal Cy5 fluorescence was seen in other organs at 30 days post-treatment **(Supplementary Figure 13**). In a side-by-side comparison, an i.v. dose of 10 mg/kg and intra-articular dose of 1 mg/kg siRNA<(EG_18_L)_2_ had remarkably similar area under the curve (AUC) profiles, suggesting that an approximately 10-fold higher dose should be used for the i.v. relative to the i.a. delivery route for siRNA<(EG_18_L)_2_ arthritic knee joint treatments. These trends were confirmed by a peptide nucleic acid (PNA) hybridization assay that enables absolute siRNA quantification in tissue. This measurement showed preferential accumulation of siRNA<(EG_18_L)_2,_ but not free siRNA, in PTOA knees over contralateral healthy knees, and similar siRNA levels (measured as ng siRNA / mg tissue) when delivered at 10 mg/kg i.v. and at 1 mg/kg i.a **(Supplementary Figure 13D**).

### siMMP13<(EG_18_L)_2_ diminishes molecular, histological, and clinical manifestations of PTOA

Therapeutic efficacy of siMMP13<(EG_18_L)_2_ was tested in mouse PTOA model after 5 weeks of mechanical knee loading (3 times / week). Mice were treated with siMMP13<(EG_18_L)_2_ or siControl<(EG_18_L)_2_ (10 mg/kg) on day 7 and again on day 21 (**Figure 3A**). The pressure-pain threshold in PTOA knees was measured by algometer, revealing a 45% reduction in threshold pressure tolerance in siControl<(EG_18_L)-treated mice, while siMMP13<(EG_18_L)_2_ lost only 25% (**Figure 3B**). The siMMP13<(EG_18_L)_2_ cohort showed relief of joint sensitivity to a level similar to mice receiving intra-articular delivery of the FDA-approved sustained release corticosteroid Zilretta (8 mg/kg) or 3X week intraperitoneal treatment with the experimental MMP13-selective small molecule inhibitor CL-82198^19^ (10 mg/kg) (**Figure 3A,B**). However, treatment with CL-82198 on the same schedule as siMMP13<(EG_18_L)_2_ (once every two weeks) produced a less favorable pain tolerance in PTAO knees, as did treatment with the pan-MMP inhibitor Marimastat (10 mg/kg, i.p., 3 times weekly).

**Figure 3.**
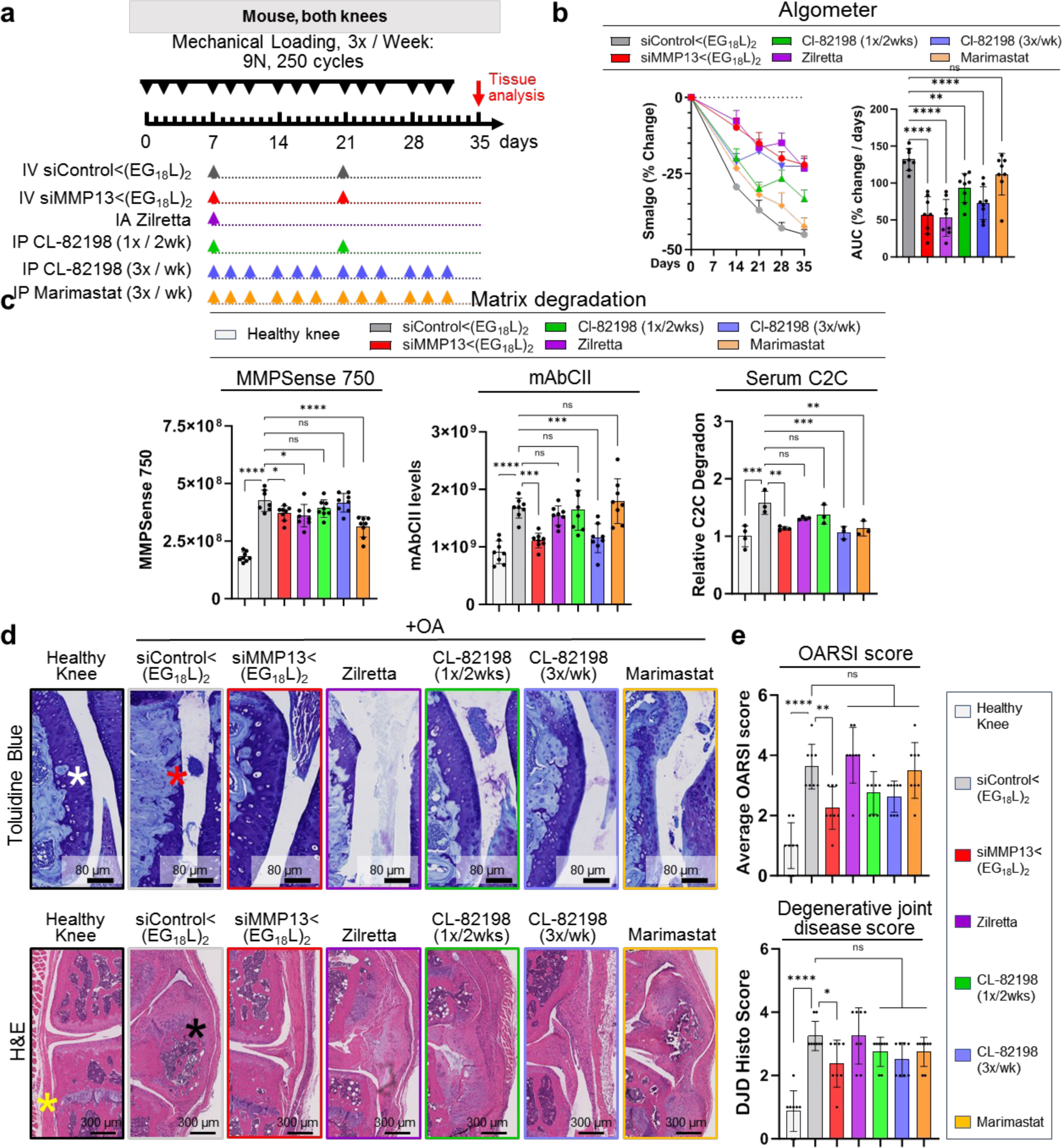
Systemic delivery of siMMP13<(EG_18_L)_2_ decreases PTOA-induced hyperalgesia and joint histopathology. **A)** Schematic timeline for knee loading and treatment. Bilateral knee loading [3X/wk, 5 wks] was used to induce severe PTOA for comparing therapeutic impact of siMMP13<(EG_18_L)_2_, Zilretta, CL-82198, and Marimastat. **B)** Mechanical hyperalgesia was measured via algometer through day 35 (left panel). Data over time displayed as mean + SEM. AUC of average Smalgo readings over time was assessed. N = 8. **C)** Total MMP activity in mouse knees was measured using MMPsense 750 Fast (N = 8; left panel). mAbCII binding in knee joints was measured by fluorescence imaging (N = 8; middle panel). C2C fragments in mouse serum was measured by ELISA (N = 3; right panel). **D)** Knee joints were stained with Toluidine Blue (upper panels, femoral condyles shown) and H&E (lower panels, synovium lining and meniscus shown). Asterisks indicate the following: healthy articular surface (white).; loss of articular surface (red); healthy synovial lining / meniscus (yellow); synovial thickening / meniscal calcification (black). **E)** Joint cartilage damage was quantitated with the OARSI osteoarthritis cartilage histopathology assessment system (upper panel) and Degenerative Joint Disease (DJD) score (lower panel). *P < 0.05, **P < 0.01, ***P < 0.001, ****P < 0.0001.

Pan-MMP activity in PTOA joints was assessed *in vivo* on treatment day 30 using MMPSense, an MMP substrate that fluoresces upon proteolytic cleavage. Intravital imaging revealed significantly elevated MMP-driven fluorescence in PTOA knees of mice treated with siControl<(EG_18_L)_2_ as compared to healthy knees (**Figure 3C**). Decreased pan-MMP activity was seen in PTOA knees of mice treated with siMMP13<(EG_18_L)_2_, Zilretta, and Marimastat, although the MMP13-selective small molecule inhibitor CL-82198 did not significantly diminish pan-MMP activity in PTOA knees. A fluorescently labeled monoclonal antibody against aberrantly exposed CII collagen (mAbCII) was used to assess relative cartilage damage^58–60^, finding significantly elevated mAbCII localization to PTOA knees (**Figure 3C**), thus confirming PTOA-induced cartilage damage in this model. However, mAbCII intensity in PTOA knees was diminished significantly in siMMP13<(EG_18_L)_2_-treated samples, approximating the mAbCII signal in healthy knees. Similar trends were observed for serum-based measurements of collagen C2C degradation fragments, a biomarker of cartilage degradation^61^. PTOA-induced mAbCII signal and serum C2C levels were also diminished by CL-82198 when dosed with higher frequency (3 times per week), underscoring the causative role ofMMP13 in PTOA-induced collagen fragmentation and cartilage damage and confirming that MMP13 targeting by siMMP13<(EG_18_L)_2_ reduces structural OA manifestations. The long-acting steroid Zilretta had little impact on mAbCII binding in PTOA knees or on serum C2C levels, consistent with observations that steroids temporarily control arthritis pain but do not target underlying joint destruction and may even accelerate cartilage thinning^62^.

Histological analyses of PTOA joints collected on treatment day 30 revealed profound disruption of cartilage architecture, proteoglycan loss, synovial thickening, osteophyte formation, and meniscal mineralization (**Figure 3D**). In PTOA knees treated with siMMP13<(EG_18_L)_2_, cartilage morphology, proteoglycan content, synovial structure, and meniscal anatomy were each relatively maintained, as quantitated using the Osteoarthritis Research Society International (OARSI) scoring criteria^63,64^ for OA progression (**Figure 3E, Supplementary Table 1**). Degenerative joint disease scoring (**Supplementary Table 1**) confirmed significantly less PTOA joint structural changes in mice treated with siMMP13<(EG_18_L)_2_ versus those treated with non-targeting siControl<(EG_18_L)_2_. Unlike what was seen with siMMP13<(EG_18_L)_2_, modest reductions in OA progression and joint degeneration were achieved in PTOA joints treated with CL-82198, although these trends did not reach statistical significance. OA progression and joint degeneration scores were not impacted by treatment with Zilretta or Marimastat.

Using micro-computed tomography (**microCT**) to quantify osteophytes and ectopic mineralization in PTOA mouse knees, the significant protection conferred by siMMP13<(EG_18_L)_2_ against meniscal mineralization and osteophyte formation was confirmed **(Extended Data Fig 5A-C).** Importantly, pathologically elevated MMP13 expression in PTOA cartilage, meniscus, and synovial compartments was reduced by treatment with siMMP13<(EG_18_L)_2_, correlating with decreased presence of degradation fragments (**Extended Data Fig 5D-G**). These data show that MMP13 silencing in arthritic joints provides both molecular and clinically tangible benefits, reducing inflammation, clinical pain, articular cartilage loss, and arthritis-associated secondary manifestations.

### MMP13 knockdown in PTOA knees reprograms pro-inflammatory gene expression patterns

To understand the gene expression patterns that underlie the therapeutic outcomes associated with MMP13 silencing, we first measured the impact of PTOA on a small panel of pro-inflammatory genes with previously described roles in OA, including *Ngf*, *Il1b*, *Tnf*, and *Cdkn2a*. These genes were each elevated in mouse PTOA knees compared to healthy knees (**Extended Data Fig 6A**). We found that PTOA-induced levels of *Ngf*, *Il1b*, *Tnf*, and *Cdkn2a* were significantly dampened as early as 5 days following i.v. delivery of siMMP13<(EG_18_L)_2_ at 10 mg/kg, gene expression changes that were sustained through day 30. These findings motivated a broader and unbiased analysis PTOA-induced gene expression changes in knee tissue, and the impact of MMP13 knockdown therein, using the nanoString nCounter Inflammation 254 gene panel. Unsupervised analysis of gene expression illustrated extensive PTOA-induced gene expression changes, including genes associated with the p38 mitogen associated protein kinase (MAPK) stress signaling pathway (**Extended Data Fig 6B, Supplementary File S1**). The serine-threonine kinase p38 MAPK phosphorylates a network of substrates that respond to cell stress^65^. Relevant to OA, p38 MAPK signaling through its substrate MAPK-associated protein kinase 2 (MK2) regulates chondrocyte differentiation, cellular senescence, MMP upregulation and pro-inflammatory cytokine production^66^. MK2 signaling within human arthritic chondrocytes drives expression of prostaglandin E2, MMP3, and MMP13, leading to arthritic pain, inflammation, and cartilage degradation^67^. Interestingly, we found that the PTOA-induced p38 MAPK gene cluster was the gene enrichment set most potently suppressed in siMMP13<(EG_18_L)_2_ treated samples (**Extended Data Fig 6C**).

Treatment with siMMP13<(EG_18_L)_2_ also significantly suppressed IL-1, chemokine, and TNFα / NFκB pathway-associated gene clusters. Notably, these clusters include several genes associated with OA pathogenesis, including those associated with chondrocyte proliferation (*Mef2d*), synovial fibroblast proliferation (*Myc*, *Jun*, *Tgfb2*, *Pdgfa*)^68,69^, angiogenesis (*Pdgfba*, *Flt1*, *Hif1a*), ECM degradation (*Mmp3, Masp1*), inflammatory signaling through Nuclear Factor-κB, (*Nf*κ*b1*, *Rela*, *Relb*), cytokines (*Il1b*, *Stat1*), chemokines (*Cxcl1*, *Cxcl10*, *Ccl3*, *Cxcr4*), and toll-like receptors (*TLR1,-2, -4, -5, -6, and -8*).

Given that many inflammatory and stress response factors induce MMP13 expression and that proteolytic matrix degradation fragments further potentiate inflammation and expression of stress response factors,^70^ these data suggest that siMMP13<(EG_18_L)_2_ uncouples a feed-forward inflammation-degradation cycle that drives arthritic joint pathology. These data support this hypothesis, given that selective MMP13 inhibition in arthritic joints suppresses expression of both pro-inflammatory factors and transcription factors that promote MMP13 expression.

### K/BxN serum transfer arthritis (STA) multi-joint rheumatoid/inflammatory arthritis mouse model

A role for MMP13 has been reported in rheumatoid arthritis (RA)^4,71–73^, including studies in which daily dosing with an experimental MMP13 small molecule inhibitor decreased RA pathogenesis in 2 models^71^. We used the transgenic K/BxN serum transfer RA model to test therapeutic efficacy of siMMP13<(EG_18_L)_2_. In this model, KRN, a T-cell receptor that binds to an endogenous glucose-6-phosphate isomerase (GPI) peptide presented by the IAg7 major histocompatibility complex class II (MHC-II) allele, drives production of anti-GPI autoantibodies and GPI-directed auto-immunity that localizes to articular cartilage/joint tissues. K/BxN mouse serum transiently confers RA to wild-type mouse serum recipients. MMP13 expression is increased in inflamed joints of K/BxN serum recipients, and MMP13-deficient mice are partially protected from K/BxN serum-induced RA^73^. Carrier-free siRNA delivery in RA has not yet been explored to our knowledge, but siMMP13<(EG_18_L)_2_ systemic administration for delivery to multiple-afflicted joints is potentially advantageous. Interestingly, albumin-based delivery approaches have been shown to facilitate therapeutic outcomes in RA preclinical-studies^45,46,74–82^. Importantly, i.v. delivery of albumin-binding Evans Blue to mice resulted in concentrated Evans Blue accumulation in arthritic forepaws and hindpaws of K/BxN serum recipients as compared to unchallenged mice (**Extended Data Fig 7A-7B**). Likewise, i.v. Cy5-labeled MSA delivered i.v. accumulated in arthritic forepaws and hindpaws of K/BxN serum recipients, but not in unchallenged wild-type mice (**Extended Data Fig 7C**). Expression of Caveolin-1 and SPARC was elevated in hindpaws of K/BxN recipient mice (**Extended Data Fig 7D**). These data confirm increased albumin retention in RA-affected joints, supporting therapeutic testing of albumin-binding siRNA<(EG_18_L)_2_ in RA models.

Mice with induced RA showed accumulation of i.v. injected Cy5-siRNA<(EG_18_L)_2_ within multiple inflamed joints (forepaw, wrist, hindpaw, ankle, knee) (**Figure 4A; Supplementary Figure 14**) at substantially higher levels than what was seen in healthy mice (**Extended Data Fig 7E-7G**) and exceeding that achieved by treatment with Cy5-siRNA or Cy5-siRNA-Chol (**Figure 4A**). Fluorescence microscopy confirmed increased siRNA<(EG_18_L)_2_ in fore- and hindpaws of K/BxN recipients, with specific accumulation in soft tissue and cartilage (**Figure 4B**). Longevity of siRNA retention was assessed in limbs of K/BXN serum recipients following a single dose Cy5 **-**siRNA<(EG_18_L)_2_. Intravital fluorescence was detected in limbs throughout the 30 days that mice were monitored (**Supplementary Figure 13**), supporting the notion of albumin-mediated delivery and retention of siRNA<(EG_18_L)_2_ in multiple afflicted joints in RA models.

**Figure 4.**
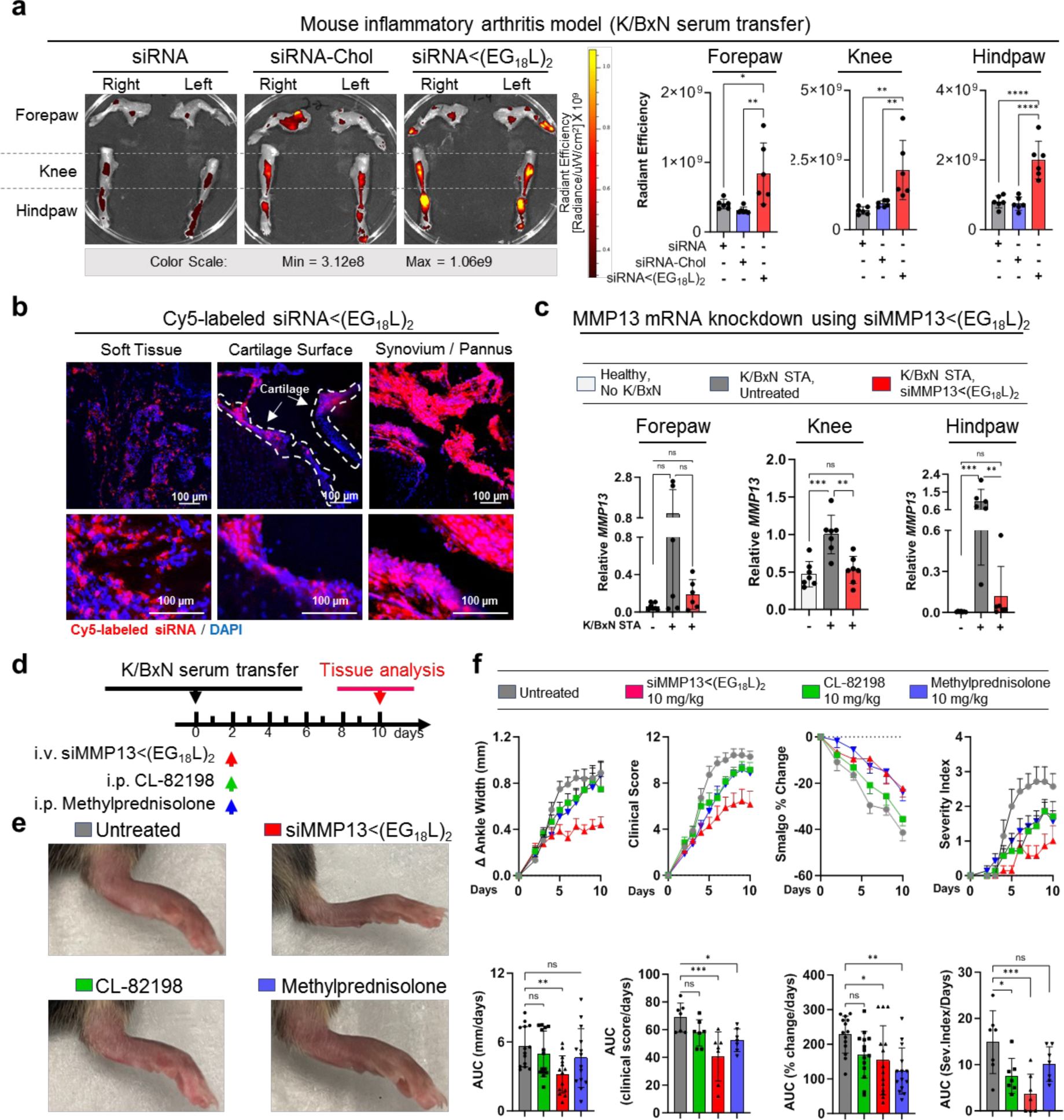
Systemic delivery enables multi-joint accumulation of siMMP13<(EG_18_L)_2_, MMP13 knockdown and diminished arthritis progression in a mouse RA model. **A-B)** Four days after K/BxN serum transfer, mice were treated with Cy5-siRNA<(EG_18_L)_2_, Cy5-siRNA-Chol, and Cy5-siRNA (1 mg/kg). Joints of forepaw, hindpaw, and knee were collected 24 hrs later and assessed by IVIS imaging (**A**) or fluorescence microscopy of hindpaw ankle joints (**B**). Representative images are shown. N = 6. **C)** Relative *Mmp13* mRNA in forepaws, knees, and hindpaws of healthy, untreated wild-type mice, or in K/BxN serum recipients treated with si-Control<(EG_18_L)_2_ or siMMP13 <(EG_18_L)_2_ was measured by RT-qPCR. N = 6. **D-F)** K/BxN serum recipients were treated with si-MMP13<(EG_18_L)_2,_ methylprednisolone, or CL-82198. Schematic timeline for K/BxN serum transfer and treatment is shown (**D**). Photos of hindpaws were collected on treatment day 10, and representative images are shown in (**E**). Ankle width change, clinical score, algometer ankle joint hyperalgesia, and severity index were measured for 10 days after treatment. Data are shown as mean + SEM. (**F**). *P < 0.05, **P < 0.01, ***P < 0.001, ****P < 0.0001. Dashed lines outline articular cartilage.

### siMMP13<(EG_18_L)_2_ therapy reduces manifestations and symptoms of RA

Strong *Mmp13* induction was seen in forepaws, knees, and hindpaws of K/BxN serum recipients as compared to healthy mice (**Figure 4C**). At treatment day 8 following a single 10 mg/kg dose of siMMP13<(EG_18_L)_2_, mRNA and protein was decreased in forepaws, knees, and hindpaws **(Figure 4C and Extended Data Fig 8A)**. Like with PTOA, MMP13 silencing in this RA model reduced expression of pro-inflammatory genes and cytokines encoding cyclooxygenase 2 (COX2), IL-6, TNF-α, and IL-1B **(Supplementary Figure 15)**. Clinical parameters of RA were also reduced upon MMP13 knockdown, including ankle swelling, pressure pain threshold, arthritis clinical score, disease severity index (**Figure 4D-4F**), and the number of afflicted joints per mouse (**Supplementary Figure 16**). In parallel studies, K/BxN serum recipients were treated with a single dose of Cl-82198 (10 mg/kg, i.p.) or methylprednisolone (10 mg/kg, i.p.), neither of which improved ankle swelling (**Figure 4D-F**). Although methylprednisolone, but not CL-82198, improved clinical score and pressure tolerance, the effect size was not as pronounced as what was seen with MMP13<(EG_18_L)_2_. Similarly, the disease severity score was somewhat improved upon treatment with CL-82198, but to a lesser extent than what was seen with siMMP13<(EG_18_L)_2_. Further, MMP13 silencing resulted in decreased mAbCII binding, indicative of cartilage preservation (**Extended Data Fig 8B**) and decreased total MMP activity (**Extended Data Fig 8C**).

Histologic examination of toluidine blue-stained cartilage harvested on treatment day 8 in K/BxN recipient mice revealed substantial cartilage destruction in hindpaws, knees, and forepaws, a phenotype that was diminished by siMMP13<(EG_18_L)_2_ treatment (**Figure 5A-5B** and **Extended Data Fig 9A-9B**; lower magnification images in **Supplementary Figures 17-20**). Inflammation scoring illustrated similar trends **(Figure 5C and Extended Data Fig 9C-D)**. MicroCT studies of bone structure in untreated K/BxN serum recipients revealed erosive bone disease characterized by increased bone surface to volume ratios, decreased bone mineral density (BMD), and decreased bone volume / total volume (BV/TV) ratio (**Figure 5D-5G**; and **Supplementary Figure 18**). Remarkably, treatment with siMMP13<(EG_18_L)_2_ protected K/BxN serum recipients from inflammation-induced changes in bone surface to volume, BMD, and BV/TV measurements. Further, bone erosions characteristic of inflammatory arthritis were evident in forepaws, knees, and hindpaws of untreated K/BxN serum, but were minimal or absent in mice treated with siMMP13<(EG_18_L)_2_ (**Extended Data Fig 9E**). In contrast, Cl-82198 and methylprednisolone did not diminish features of erosive bone disease detected by microCT. Serum C2C degradation fragments were significantly elevated in untreated, CL-82198 treated, and methylprednisolone-treated K/BxN serum recipients (**Extended Data Fig 10A**), while treatment with siMMP13<(EG_18_L)_2_ significantly reduced serum C2C fragments. These findings were confirmed using IHC for C1,2C in K/BxN serum recipients hindpaw/ankle and knee joint tissues **(Extended Data Fig 10B)**.

**Figure 5.**
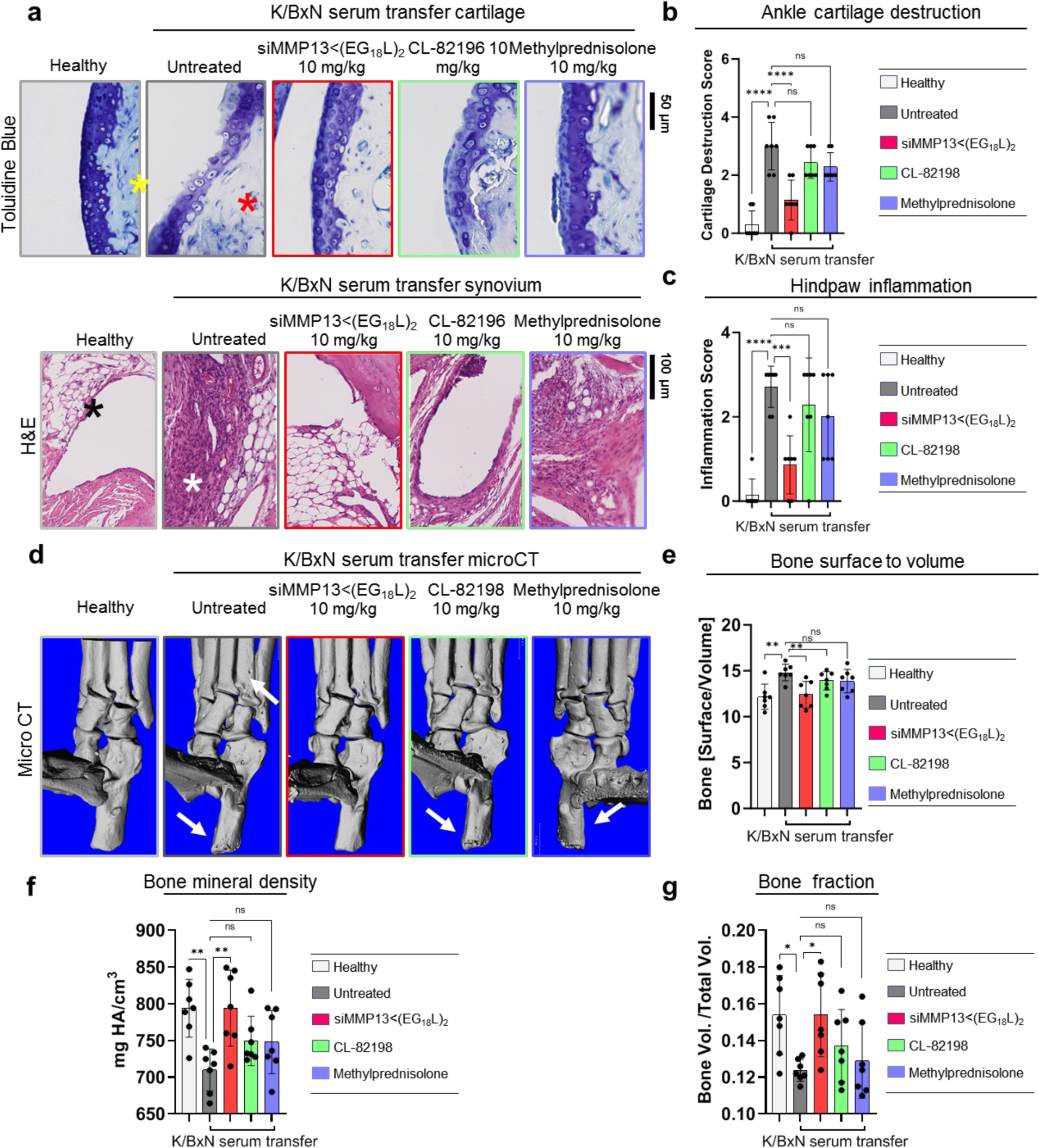
Systemic siMMP13<(EG_18_L)_2_ treatment protects against cartilage / bone destruction and synovial inflammation in a mouse RA model. K/BxN serum recipient mice were treated with siMMP13<(EG_18_L)_2,_ methylprednisolone, or CL-82198. **A-C)** Histological analysis of toluidine blue-stained ankle cartilage sections (**A**, upper panels) and H&E-stained hindpaw sections (**A**, lower panels) were used for cartilage destruction scoring (**B**) and inflammation scoring (**C**), respectively. Asterisks indicate the following: yellow: articular surface; red: articular surface damage. Black: healthy synovium. White: synovial inflammation. **D-G)** MicroCT-based **3D** renderings of hindpaws scans (D, representative images) were used to measure bone surface:volume ratio (**E**), bone mineral density **(F**) and bone fraction (**G**). White arrows in D show bone erosions *P < 0.05, **P < 0.01, ***P < 0.001,

Hindpaw ankle articular cartilage and synovial RNA was harvested from K/BxN serum transfer recipients on treatment day 8 and assessed using the nanoString nCounter mouse inflammation expression panel. These studies identified several gene clusters that were elevated in untreated K/BxN serum transfer recipients as compared to healthy mice (**Supplementary File S1**), many of which were downregulated in samples collected from K/BxN serum transfer recipients treated with siMMP13<(EG_18_L)_2_ (**Extended Data Fig 10C**). Treated RA animals had gene expression patterns that mirrored the changes seen in PTOA knees upon MMP13 knockdown (*e.g.,* IL-1 signaling, NF-kB signaling, and chemokine pathways were suppressed by treatment, **Extended Data Fig 10D**). Additionally, the pro-inflammatory COX-2 encoding gene *Ptgs2*, the chemokine genes *Cxcl5, Ccl7*, *Ccl19*, and the RA-associated macrophage genes *Tnfsf14* and *Tr2* were significantly reduced in K/BxN hindpaws treated with siMMP13<(EG_18_L)_2_. Thus, the molecular markers of inflammation that characterize RA are dampened by MMP13 knockdown achieved by i.v. delivery of siMMP13<(EG_18_L)_2_.

Importantly, serum levels of toxicity markers, including blood urea nitrogen (BUN), alanine transaminase (ALT), aspartate amino transferase (AST), creatine phosphokinase (CPK), lactate dehydrogenase (LDH), and glucose, were detected within the normal ranges, lacking significant differences between healthy, PTOA, or K/BxN serum recipient mice in any of the treatment groups (**Supplementary Figure 21)**. Liver, kidney, lung, heart, and spleen of mice within all groups also showed a normal gross and histological structure (**Supplementary Figure 22**).

### MMP13 silencing in a Dunkin-Hartley guinea pig ACL transection (ACLT) large animal model

ACL transection (ACLT) was done on the left knee of Dunkin Hartley guinea pigs (**DHGP**s) to establish proof of concept in PTOA in a larger species (**Figure 6A**). Like humans, DHGPs develop OA with aging, with the medial knee compartment showing loss of proteoglycans, fibrillation, cloning of chondrocytes, osteophyte formation, and subchondral bone sclerosis^83^. ACLT is used to mimic PTOA and accelerate the spontaneous OA phenotype^84^. MMP13 is strongly expressed and highly localized with OA cartilage lesions in DHGPs^83^. Cy5-siRNA<(EG_18_L)_2_ was delivered i.v. on post-surgical day 13 (1 mg/kg, i.v.), and knees were imaged *ex vivo* to measure Cy5 fluorescence on day 14, revealing significantly increased Cy5-siRNA<(EG_18_L)_2_ accumulation in ACLT knees over unaltered knees (**Figure 6B**), while free Cy5-siRNA did not exhibit preferential accumulation in ACLT damaged knees. Ex vivo Cy5 imaging of organs showed abundant free siRNA in kidneys, while the albumin binding siRNA<(EG_18_L)_2_ was found at highest levels in ACLT-damaged knees and liver, but not in healthy knees nor in kidneys (**Figure 6C**). Fluorescence microcopy confirmed delivery of Cy5-siRNA<(EG_18_L)_2_ but not free siRNA, to synovia of ACLT-damaged knees, with strong penetration into cartilage (**Figure 6D**). Intravenous siMMP13<(EG_18_L)_2_ delivered on post-surgical day 7 at 10 mg/kg decreased *Mmp13* expression in ACLT-damaged knees by >80% (**Figure 6E**), and MMP13 protein expression by >70% on day 14 (**Figure 6F-G)**.

**Figure 6.**
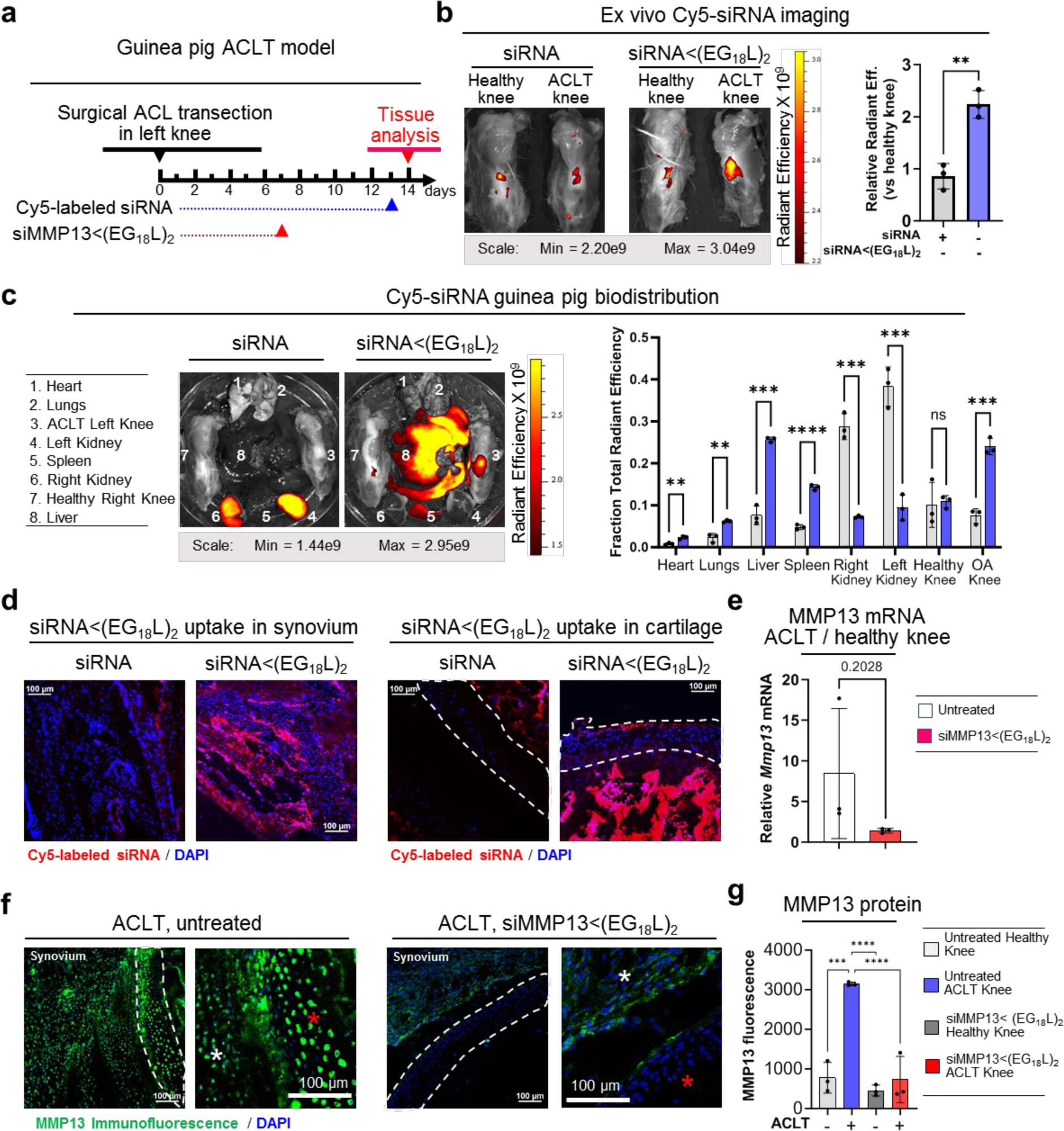
MMP13 silencing upon systemic siMMP13<(EG_18_L)_2_ delivery in guinea pig ACL transection model of arthritis. **A)** Timeline for ACL transection and siRNA delivery in Dunkin-Hartley guinea pigs is shown. **B-D)** Cy5 fluorescence was measured ex vivo in healthy and ACLT knees (**B**) and organs compared using multiple unpaired t-tests with no correction for multiple comparisons (**C**) of guinea pigs by IVIS, and by fluorescence microscopy of sagittal ACLT knee cartilage / synovial cryosections **(D)** at 24 hrs after i.v. delivery of Cy5-siRNA or Cy5-siRNA<(EG_18_L)_2_ at 1 mg/kg. Representative images are shown. In D, Blue = DAPI, Red = Cy5 siRNA. **E)** Guinea Pig *Mmp13* mRNA expression (ACLT/Healthy Knee) as determined by qRT-PCR. **F-G)** MMP13 immunofluorescence in cartilage / synovial cryosections of ACLT knee joints that were untreated (left) or treated with siMMP13<(EG_18_L)_2_ (10 mg/kg, i.v.). White asterisk: synovium; red asterisk; femoral condyle cartilage. Representative images are shown (**F**). MMP13 immunofluorescence intensity was quantitated using morphometric software (**G**). *P < 0.05, **P < 0.01, ***P < 0.001, ****P < 0.0001. Dashed lines outline articular cartilage.

## Conclusion

A unique albumin-binding RNAi conjugate (siRNA<(EG_18_L)_2_) was engineered for MMP13-selective gene targeting in the context of arthritis. This optimized siRNA-lipid conjugate was capable of preferential accumulation and long-term retention within arthritic joints, robustly decreasing arthritis-induced MMP13 and interrupting the forward-feeding cycle of inflammatory joint damage to limit arthritis progression. Overall, this albumin-binding siRNA-L_2_ system shows promise as a systemic and/or local anti-MMP13-specific therapy. This siRNA delivery platform is readily amenable to modular engineering for targeting other arthritis disease-driving genes and/or gene combinations. Beyond the scope of arthritis, we believe this platform may be utilized to deliver to other diseased or injured tissues characterized by inflammation, including tumors, tissues undergoing ischemic injury, and bone fractures.

## Supplementary files

**File S1.** Excel file of gene names and fold expression changes in knee RNA samples harvested from healthy mice, untreated PTOA mice, and PTOA mice treated with siMMP13<(EG_18_L)_2_ as well as hindpaw RNA samples harvested from healthy mice, untreated K/BxN serum recipient mice, and K/BxN serum recipients treated with siMMP13<(EG_18_L)_2_.

## Author contributions

J.M.C. and C.L.D. conceptualized the study and experiments. J.M.C., E.N.H., J.H.L, A.S, and N.F, synthesized and characterized the siRNA conjugates and assisted with PNA hybridization. J.M.C. and M.C.K. performed *in vitro* siRNA studies. M.C.K, J.T.M, and C.R.D. performed in-vivo flow experiments. J.M.C., F.Y. performed *in vivo* pharmacokinetic studies. J.M.C. performed *in vivo* confocal microscopy imaging and analyses. D.L.M performed SEC synovial fluid analysis. J.M.C., F.Y., V.S. performed *in vivo* therapeutic studies including MMPsense, qRT-PCR, microCT, and histological tissue preparation. K.N.G-C performed pathologist-blinded OARSI, DJD, and K/BxN STA scoring of histological sections. L.C. provided K/BxN serum and experimental insight for inflammatory arthritis studies. H.C. and K.A.H. provided mAbCII680 reagent and experimental insight for *in vivo* cartilage damage studies. T.M provided insight for in vivo flow cytometry experimental design and performed and contributed the single cell RNA-Seq analysis. R.C. and C.L.D. provided study mentorship and oversight. J.M.C., R.C, C.L.D. co-wrote the manuscript with input from all authors.

## Acknowledgements and funding

The authors acknowledge the assistance of the Vanderbilt Translational Pathology Shared Resource (TPSR), which is supported by NCI/NIH Cancer Center Support Grant 2P30 CA068485-14 and the Shared Instrumentation Grant S10 OD023475 for the Leica Bond Rx. Micro CT analyses was supported in part by the NIH (S10RR027631-01). The histological technical assistance of Cindy Lowe and Karen Thompson and the microCT technical assistance by Sasidhar Uppuganti is acknowledged. Further, we are appreciative for the help of Alexander Knights in planning and interpreting flow cytometry experiments. Flow Cytometry experiments were performed in the VMC Flow Cytometry Shared Resource. The VMC Flow Cytometry Shared Resource is supported by the Vanderbilt Ingram Cancer Center (P30 CA68485) and the Vanderbilt Digestive Disease Research Center(DK058404). We are grateful for funding from the DOD (DOD CDMRP OR130302), NIH (NIH R01 CA224241, NIH R01 EB019409, NIH R21 AR078636, NIH T32GM007347), Canadian Natural Sciences and Engineering Research Council (NSERC), the Rheumatology Research Foundation (RRF), and the VA Merit Awards BX004151 and BX004472. The content in this report is solely the responsibility of the authors and does not necessarily represent the official views of the funding agencies.

---

**Extended Data Figure 1.**
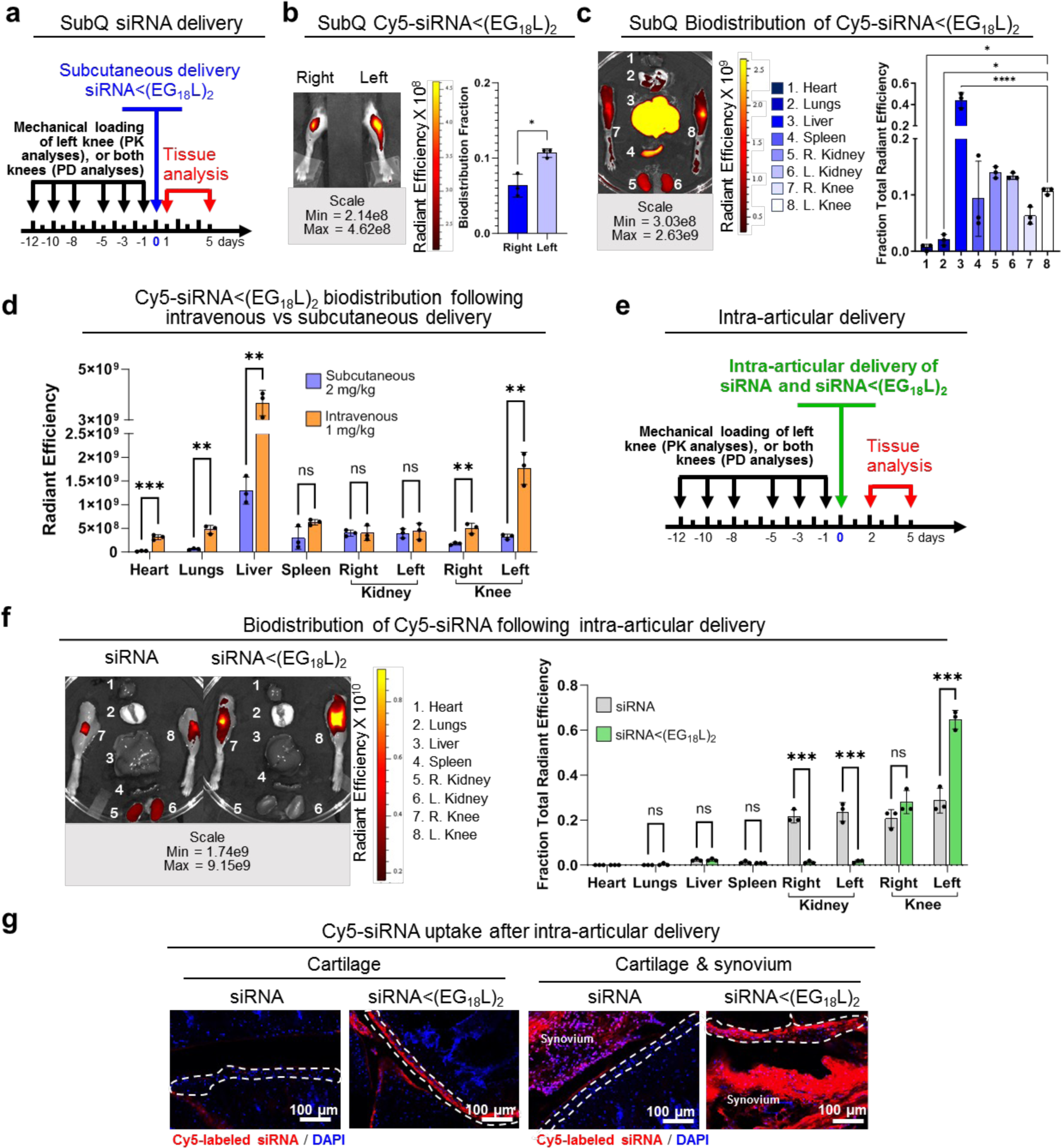
Exploring different routes of delivery in PTOA mouse model. **A-C)** Subcutaneous (SubQ) delivery of Cy5-siRNA<(EG_18_L)_2._ (2 mg/kg) was studied in a PTOA mouse model. Timeline for mechanical knee loading and treatment is shown (A). Intravital (B) and ex vivo (C) Cy5 fluorescence imaging was used to measure Cy5-siRNA<(EG_18_L)_2._ biodistribution to paired healthy and loaded knees and organs 24 hrs after delivery. N = 3. **D)** Quantification of SubQ vs. intravenous (i.v.) delivery route organ biodistribution 24 hours after injection. Analyzed using multiple unpaired t-tests with no correction for multiple comparisons. **E-G)** Intra-articular delivery of Cy5-siRNA<(EG_18_L)_2._ (1 mg/kg) was studied in a PTOA mouse model. Timeline for mechanical knee loading and treatment is shown (**E**). Ex vivo Cy5 fluorescence imaging was used to measure Cy5-siRNA<(EG_18_L)_2._ biodistribution to paired healthy and loaded knees and organs 48 hrs after delivery (**F**). N = 3. Analyzed using multiple unpaired t-tests with no correction for multiple comparisons. (**G**). Cryohistology of the loaded knee joints 48 hours after intra-articular injection with a focus on cartilage and synovial tissues. N = 3. *P < 0.05, **P < 0.01. Dashed lines outline articular cartilage.

**Extended Data Figure 2.**
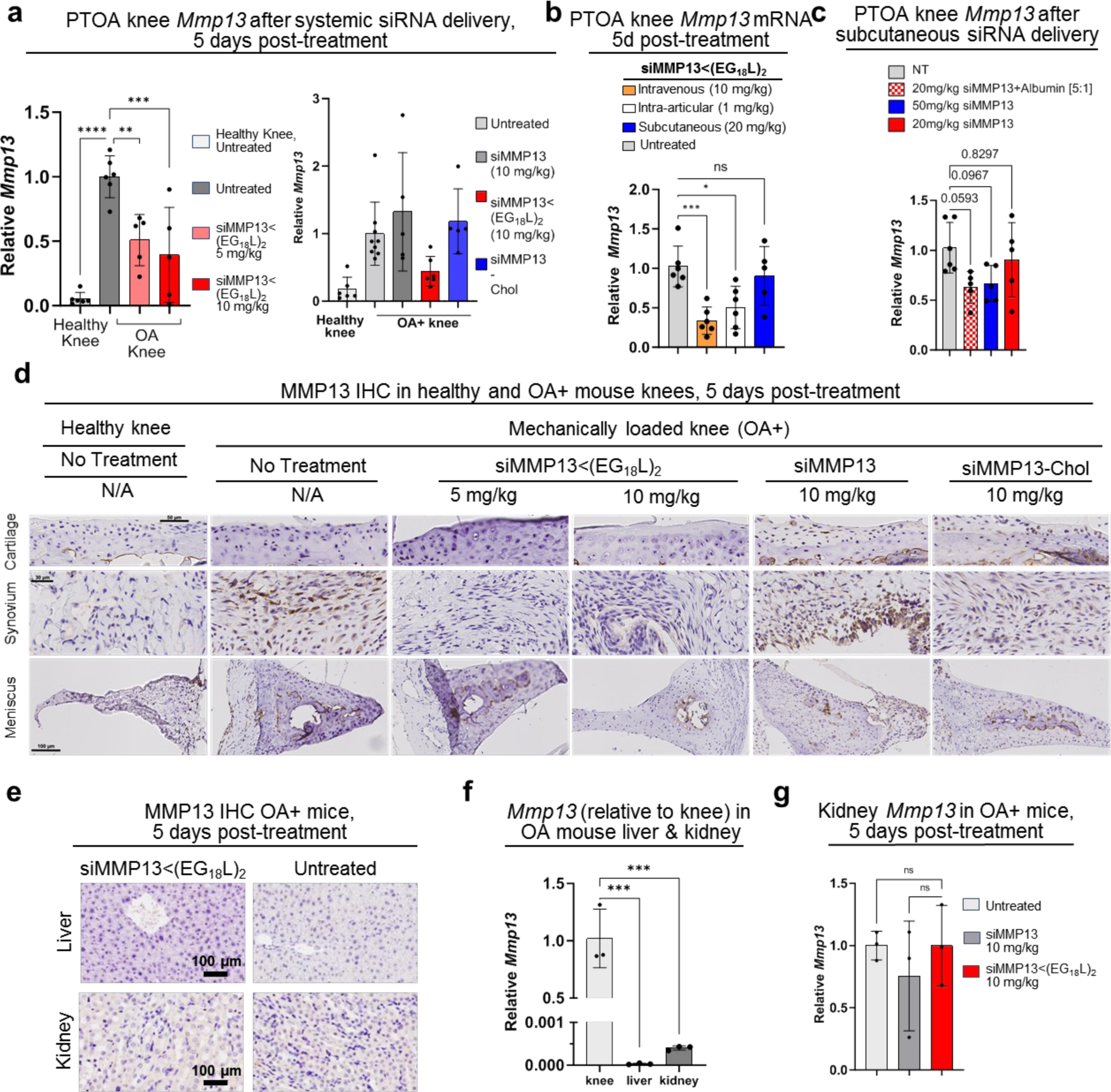
MMP13 knockdown in PTOA joints by systemic delivery of siMMP13<(EG_18_L)_2_. **A)** *Mmp13* mRNA was measured by RT-qPCR in healthy and PTOA knees of mice 5 days after i.v. delivery of siMMP13<(EG_18_L)_2_, siMMP13-Chol, or free siMMP (5 and 10 mg/kg, where indicated). **B)** *Mmp13* mRNA was measured by RT-qPCR in PTOA mouse knees of mice 5 days after i.v., i.a., or subcutaneous siMMP13<(EG_18_L)_2_ delivery at the doses indicated. **C)** *Mmp13* mRNA was measured by RT-qPCR in PTOA mouse knees of mice 5 days subcutaneous siMMP13<(EG_18_L)_2_ delivery at the doses indicated. **D-E)** MMP13 IHC in PTOA knees (D), liver and kidneys (E), 5 days after i.v. delivery (10mg/kg) siMMP13<(EG_18_L), siMMP13-Chol, free siMMP13, or no treatment. **F)** Relative *Mmp13* mRNA expression in liver and kidneys of mice (relative to knee *Mmp13* levels). **G)** Relative *Mmp13* mRNA in kidneys, 5 days post-injection with 10 mg/kg siMMP13<(EG_18_L)_2_, free siMMP13, or PBS. *P < 0.05, **P < 0.01, ***P < 0.001

**Extended Data Figure 3.**
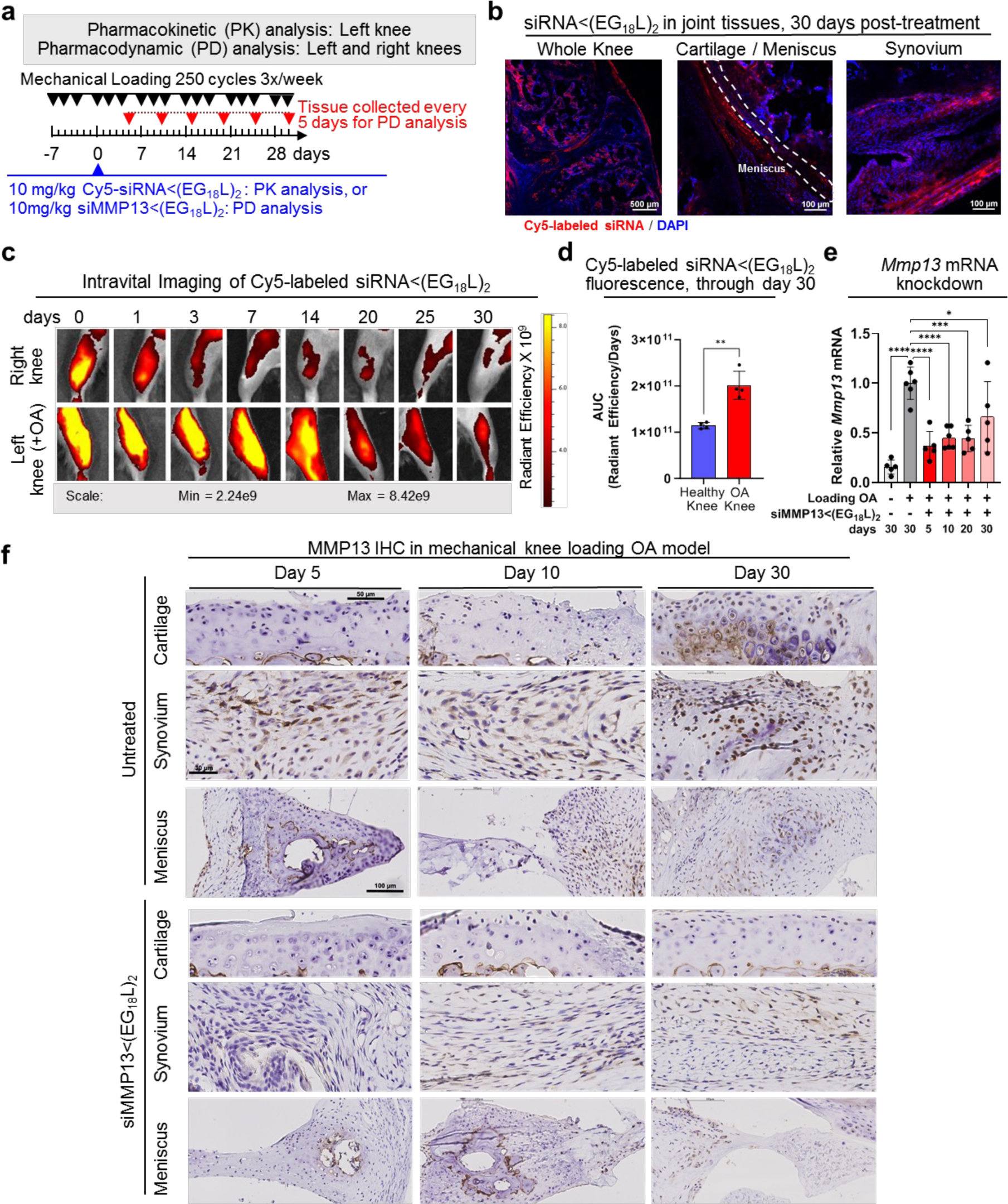
siMMP13<(EG_18_L)_2_ demonstrates long-term retention and MMP13 silencing in PTOA knee joints. **A)** Timeline for mouse left knee mechanical loading (3 times / week) and single treatment (day 7) with Cy5-siRNA<(EG_18_L)_2_ or siMMP13<(EG_18_L)_2._ **B-D)** Cy5-siRNA was measured on day 30 post-treatment by confocal microscopy of knee joint cryosections (**B**, representative images), and at time points from 0-30 days post-treatment by longitudinal intravital Cy5 imaging (**C**). Semiquantitative analysis of the AUC (Cy5 fluorescence / days) was calculated (**D**) (n = 4). **E-F).** Knee joint *Mmp13* was measured at the indicated time points by RT-qPCR (**E**) and IHC (**F**) in healthy or PTOA knee tissues from mice treated with or without siMMP13<(EG_18_L)_2_ collected at the indicated time points. N = 5. *P < 0.05, **P < 0.01, ***P < 0.001, ****P < 0.0001. Dashed lines outline articular cartilage.

**Extended Data Fig 4.**
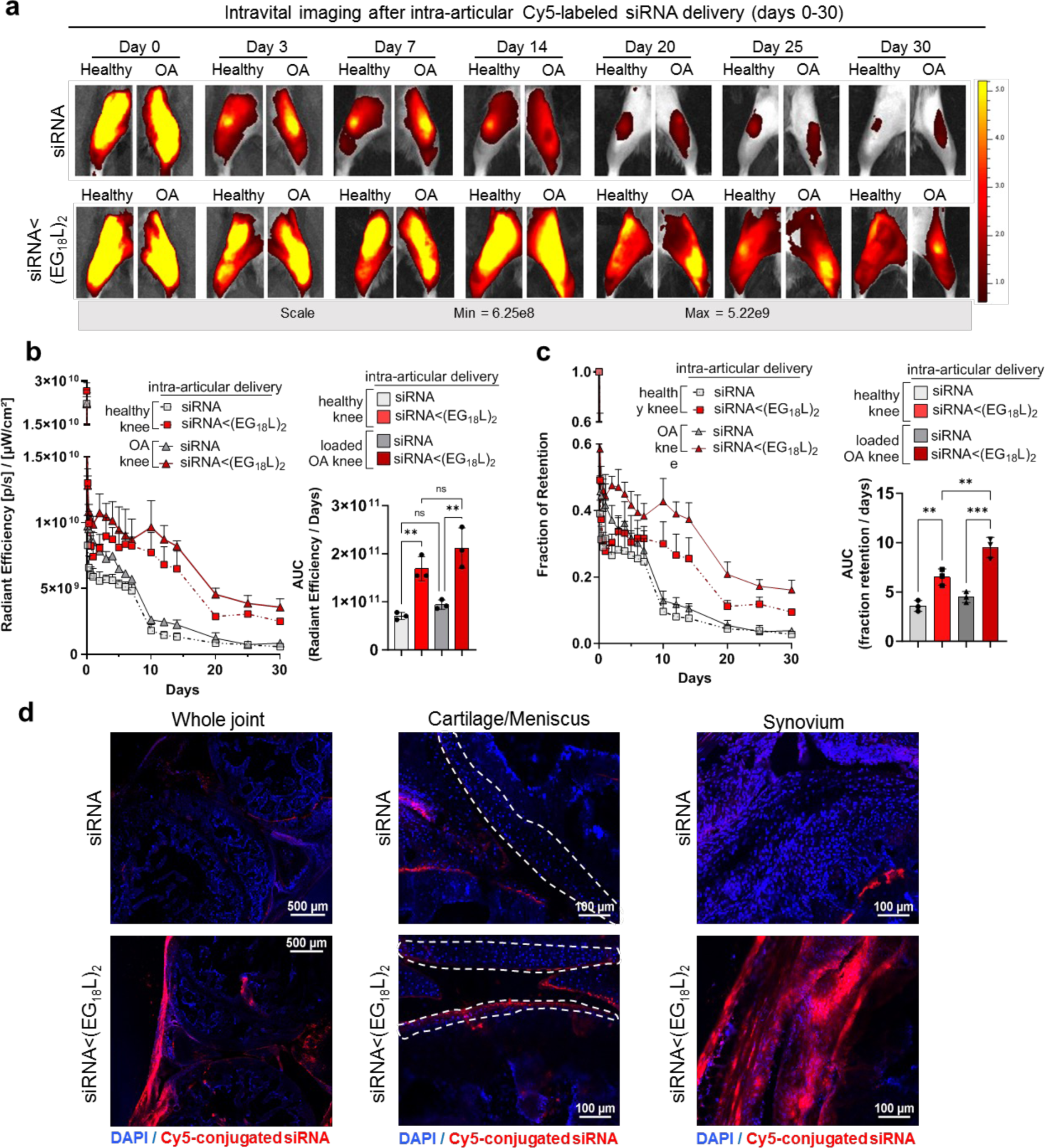
Healthy and PTOA knee joint delivery and retention after intra-articular delivery over a 30-day time course. **A)** Representative IVIS images of healthy and PTOA mouse knee joints over 30 days after a single 1 mg/kg intra-articular (IA) injection of Cy5-siRNA<(EG_18_L)_2_ or Cy5-siRNA. **B)** Semiquantitative analysis of time-course fluorescent radiant efficiency within healthy and OA mouse knee joints over 30 days (N = 3) and AUC based on the fluorescence intensity profiles. Radiant efficiency over time plotted as mean + SEM. **C)** Semiquantitative analysis of time-course fraction of retention within healthy and OA mouse knee joints over 30 days and AUC based on the fraction of retention profiles. **D)** Representative confocal microscopy of knee joint cryosections at the 30-day endpoint showing IA Cy5-siRNA<(EG_18_L)2, and IA Cy5-siRNA in loaded, PTOA, knees with a specific focus on synovial and cartilage/meniscus tissues. *P < 0.05, **P < 0.01, ***P < 0.001, ****P < 0.0001. Dashed lines outline articular cartilage.

**Extended Data Fig 5.**
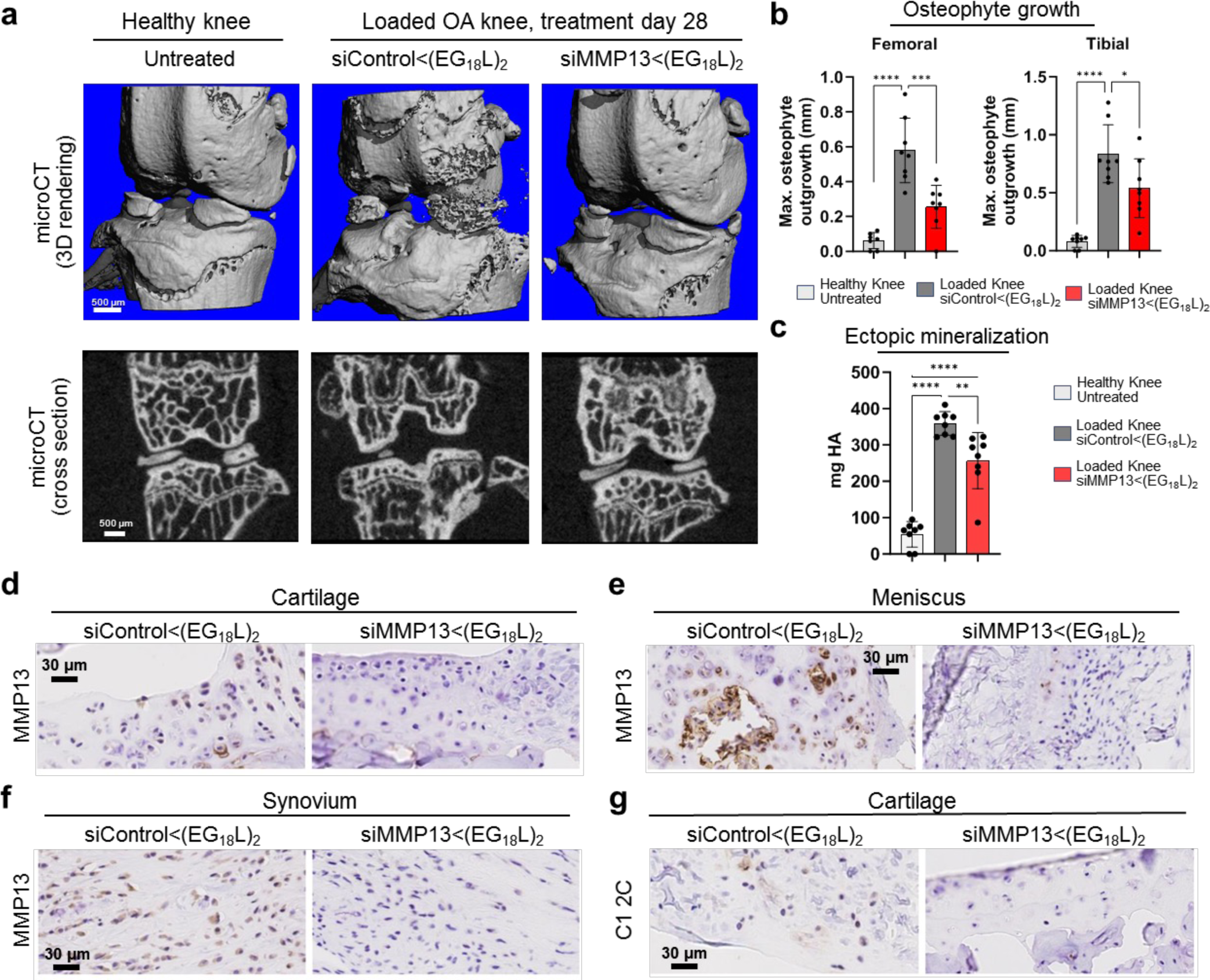
Systemic albumin-hitchhiking siMMP13<(EG_18_L)_2_ treatment provides whole knee joint protection by reducing osteophyte formation and pathologic mineralization meanwhile by MMP13 protein and collagen degradation fragments. **A)** MicroCT-based 3-dimensional (3D) renderings of meniscal/ectopic mineralization and osteophyte growth in healthy and PTOA mice treated with siMMP13<(EG_18_L)_2_ or siControl<(EG_18_L)_2_. **B)** Measurements of femoral and tibial osteophyte size at largest outgrowth from normal cortical bone structure. **C)** Ectopic mineralization [hydroxyapatite (HA)] in menisci and in the form of osteophytes was measured. **D-F)** MMP13 IHC analyses of cartilage (D), meniscus (E), and synovium (F) and showing reduced MMP13 protein levels when treated with siMMP13<(EG_18_L)_2_. **G)** C1,2C collagen 2 degradation fragment IHC. Samples here are from the PTOA therapeutic study in Main Figure 3. *P < 0.05, **P < 0.01, ***P < 0.001, ****P < 0.0001.

**Extended Data Fig 6.**
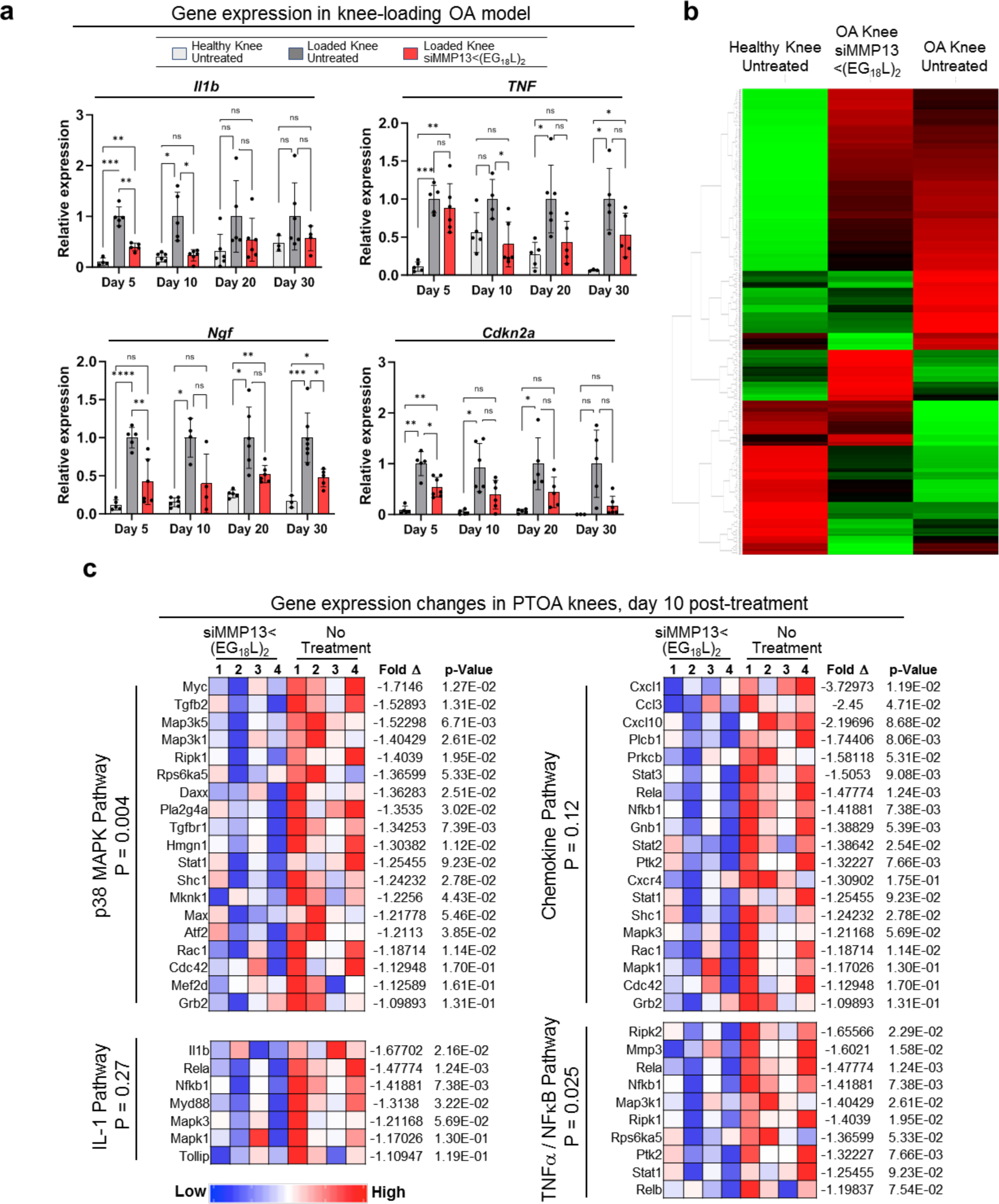
Systemic siMMP13<(EG_18_L)_2_ delivery alters inflammatory gene expression profiles in mechanically-loaded PTOA joints. **A)** qRT-PCR of IL-1B, TNFalpha, NGF, and P16INK4a (Cdkn2a) at days 5, 10, 20, and 30 after a single i.v. injection of 10 mg/kg siMMP13<(EG_18_L)_2._ Analyzed using a mixed effects analysis. **B)** Unsupervised sorting of treatment groups as quantified by nanoString at day 10 after treatment with 10 mg/kg siMMP13<(EG_18_L)_2_. Gene expression is shown as high-(green) or low-expression (red) sorted vertically by differences between treatment groups. **C)** Gene cluster expression changes at day 10 post-treatment. *P < 0.05, **P < 0.01, ***P < 0.001, ****P < 0.0001.

**Extended Data Fig 7.**
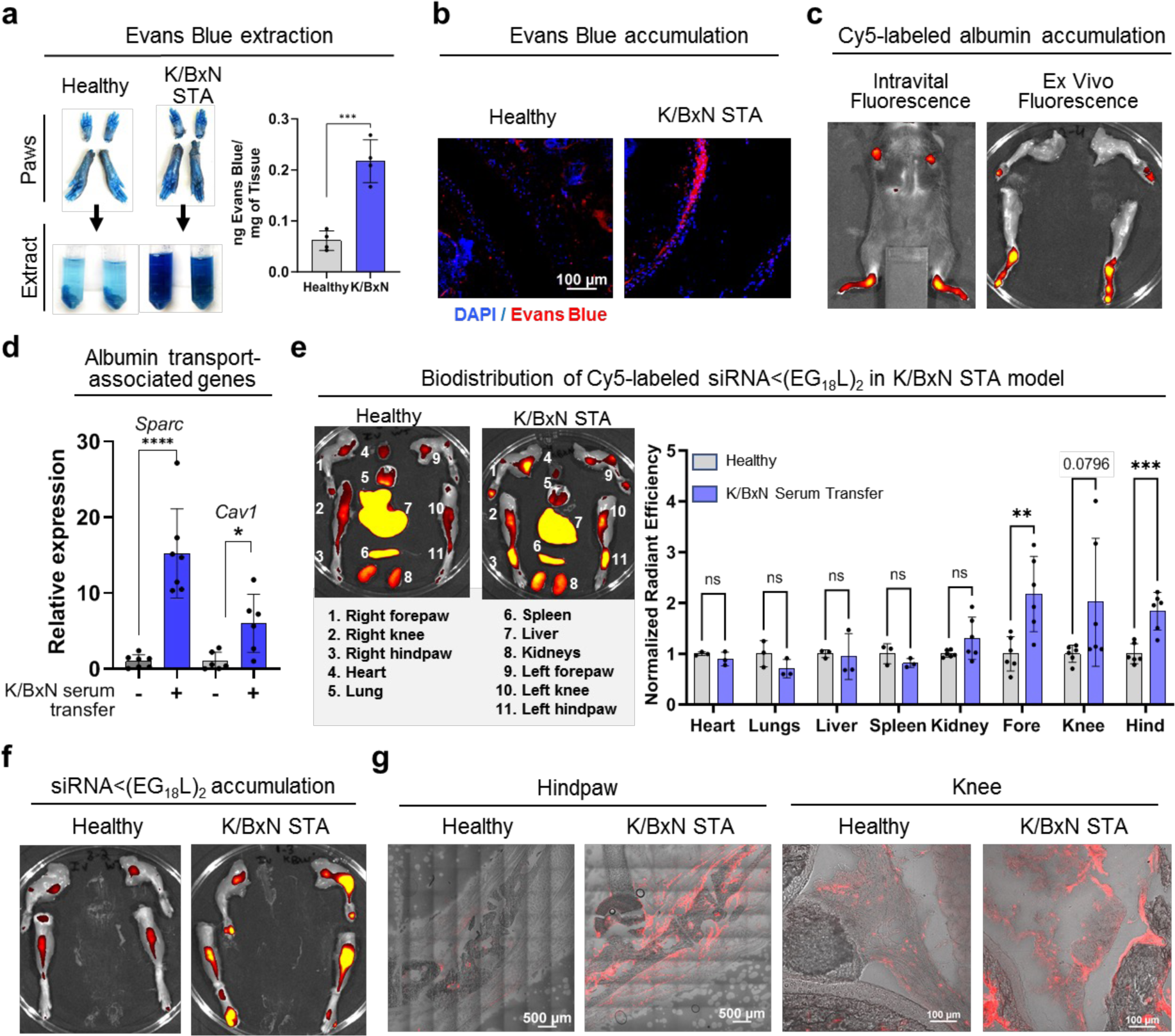
Characterizing albumin-based drug delivery in the K/BxN STA model. **A-C)** Evans Blue or Cy5-MSA was delivered i.v. to wild type (healthy) and K/BxN serum recipient mice 4 days after serum transfer. Joints of forepaws and hindpaws were used for Evans Blue extraction (**A**, representative images), measuring Evans Blue absorbance in extracts, or for visualization of Evans Blue by fluorescence microscopy in joint cryosections (**B**). MSA-Cy5 delivery to joints was assessed by intravital Cy5 fluorescence imaging (**C**). **D)** qRT-PCR was used to measure relative gene expression of SPARC and Caveolin-1 in healthy and K/BxN arthritis hind paws. **E-G)** Cy5-siRNA<(EG_18_L)_2_ (1 mg/kg, i.v.) was delivered to healthy and K/BxN serum recipient mice. Cy5 fluorescence in organs (**E**) and limb joints analyzed using multiple unpaired t-tests with no correction for multiple comparisons (**F**) were measured ex vivo at 24 hrs, and in knee and hindpaw cryosections by fluorescence microscopy (**G**) *P < 0.05, **P < 0.01, ***P < 0.001, ****P < 0.0001.

**Extended Data Fig 8.**
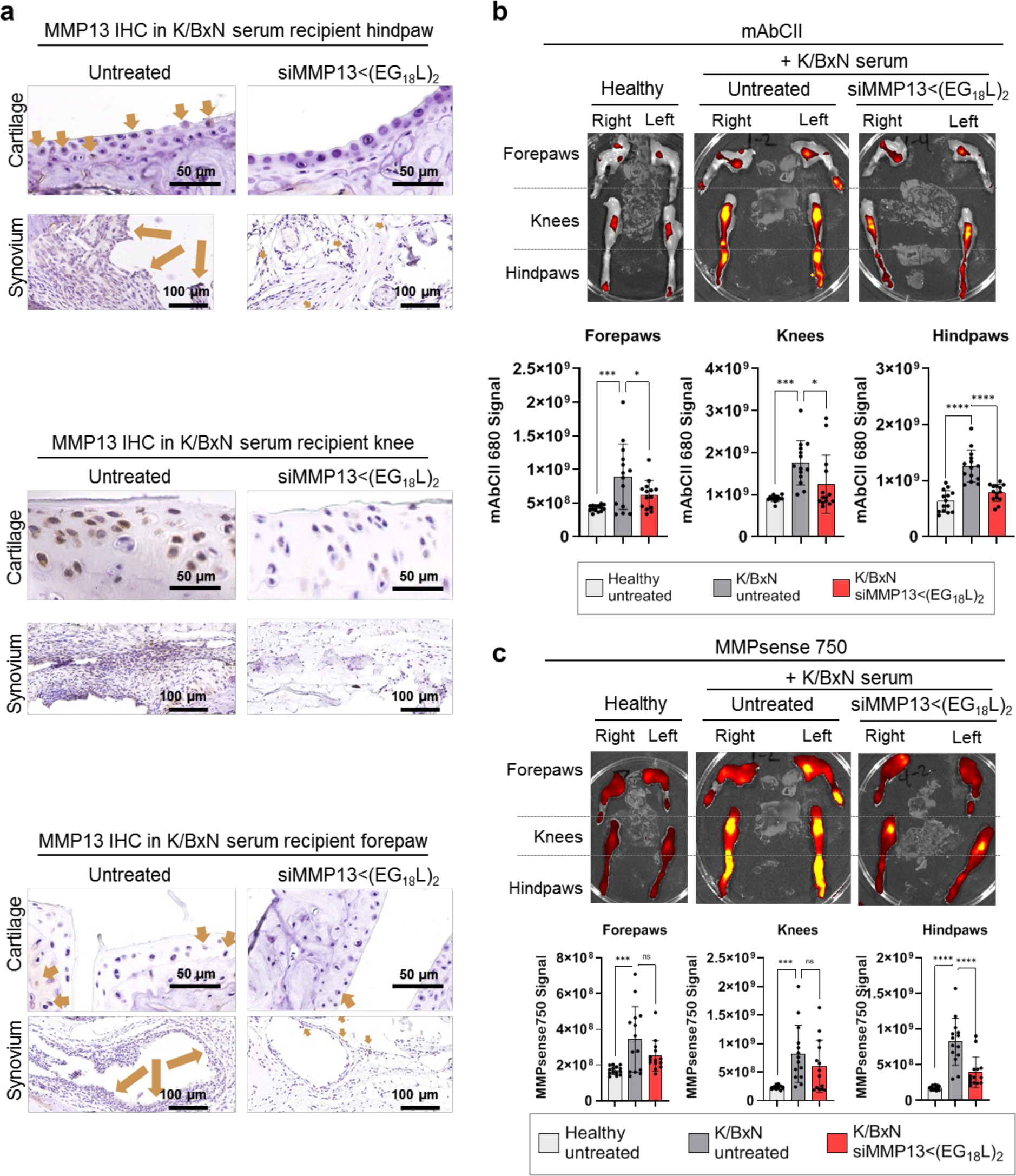
Systemic siMMP13<(EG_18_L)_2_ delivery silences MMP13 and protects cartilage in the multi-joint K/BxN STA model. K/BxN STA mice were treated with or without siMMP13<(EG_18_L)_2_ (10 mg/kg, i.v.), then assessed at day 10. **A)** MMP13 IHC analysis of cartilage and synovium in multiple joints was used to assess MMP13 expression. **B)** Fluorescent mAbCII was delivered to mice, then detected by ex vivo fluorescence imaging (IVIS) to measure relative cartilage damage in healthy and K/BxN STA mice (representative images shown). **C)** MMPsense 750 Fast was delivered to mice, then detected by ex vivo fluorescence imaging (IVIS) used to measure pan-MMP activity. *P < 0.05, **P < 0.01, ***P < 0.001, ****P < 0.0001.

**Extended Data Fig 9.**
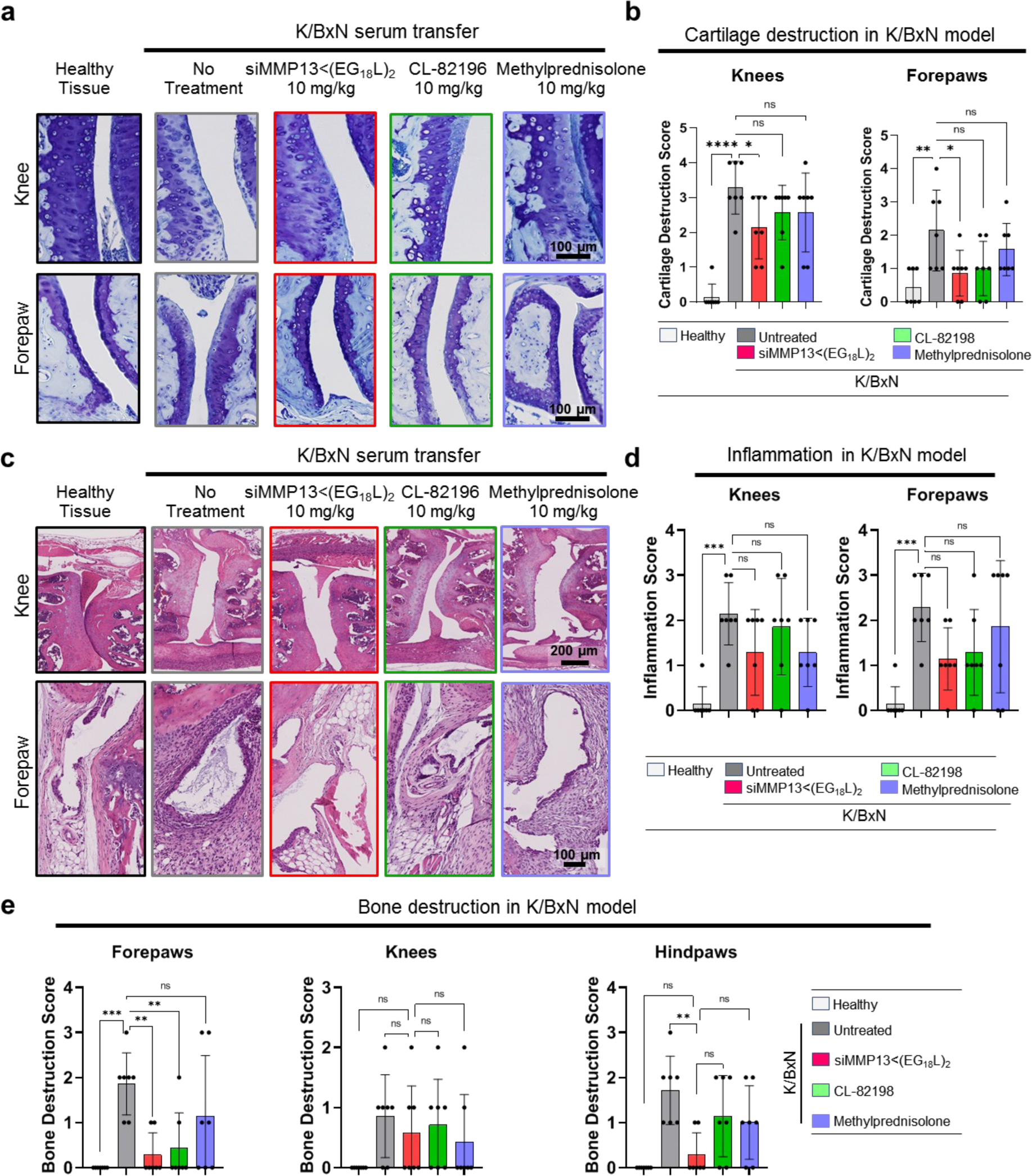
Systemic delivery of siMMP13<(EG_18_L)_2_ protects cartilage and bone while reducing inflammation in multiple joints of the K/BxN STA model. **A-B)** Toluidine Blue stained histological knee and forepaw sections (**A**) were scored using the Cartilage Destruction Score (**B**). Hindpaw data is shown in Figure 5. **C-D)** H&E-stained histological knee and forepaw sections (**C**) were scored using the Inflammation Score (**D**). Hindpaw data is shown in Figure 5. **E)** Histological bone destruction score quantification of hindpaw, knee, and forepaw joint tissues. *P < 0.05, **P < 0.01, ***P < 0.001, ****P < 0.0001.

**Extended Data Fig 10.**
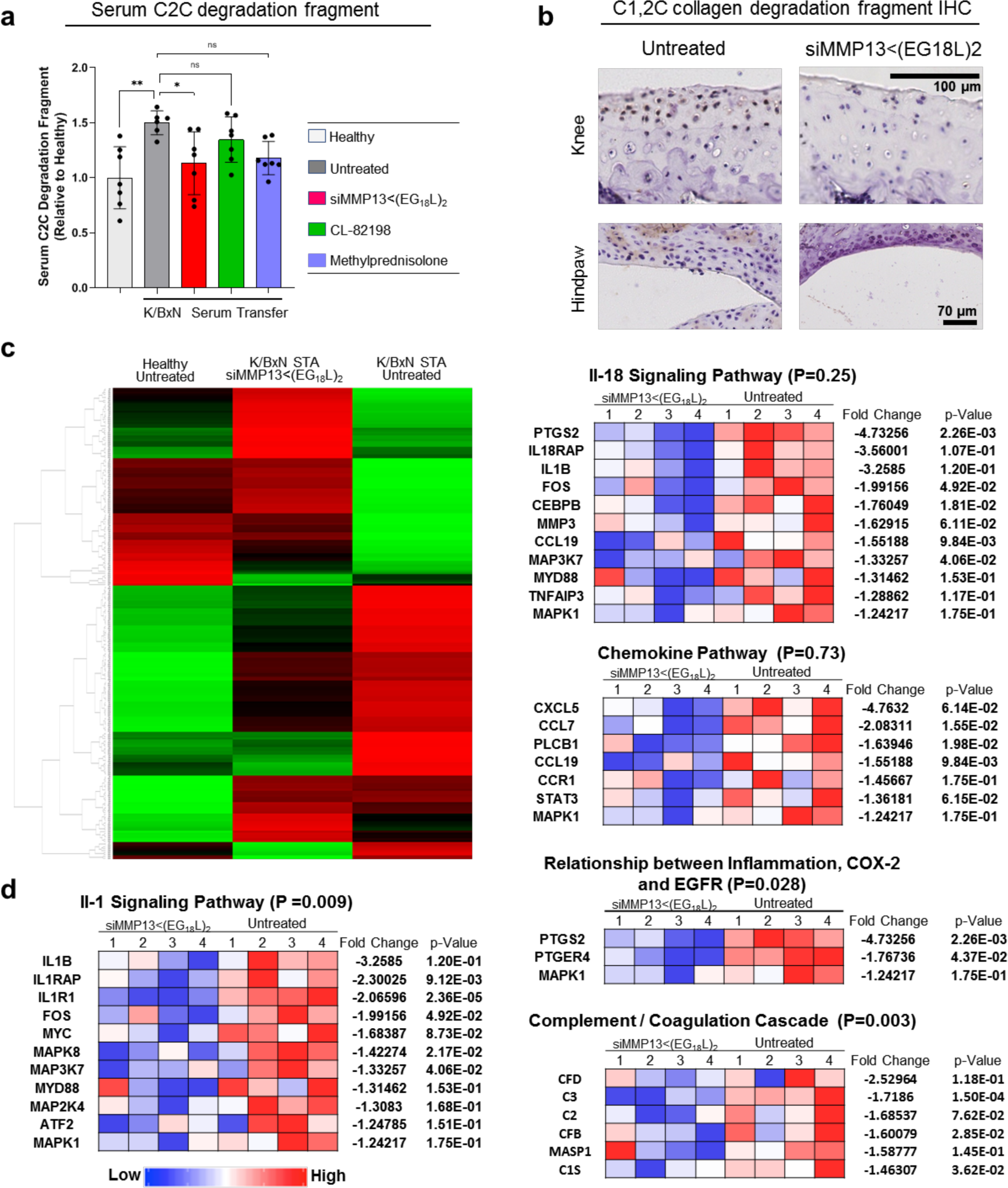
Systemic siMMP13<(EG_18_L)_2_ delivery reduces cartilage degradation fragments and diminishes the inflammatory gene expression profile in the K/BxN STA model. **A)** Relative serum C2C collagen 2 degradation fragment was measured on day 10 by ELISA. **B)** C1,2C collagen 2 degradation fragment was assessed by IHC on day 10. **C-D)** RNA harvested from K/BxN recipient mouse joints was assessed by nanoString nCounter analysis for the Mouse Inflammatory 294 Gene Expression panel. Clustering of gene expression changes [high-(green) or low-(red)] sorted vertically by differences between treatment groups (**C**). Treatment group-associated changes in gene enrichment clusters ion treatment day 10 (**D**). *P < 0.05, **P < 0.01, ***P < 0.001, ****P < 0.0001.

**Supplementary Figure 1.**
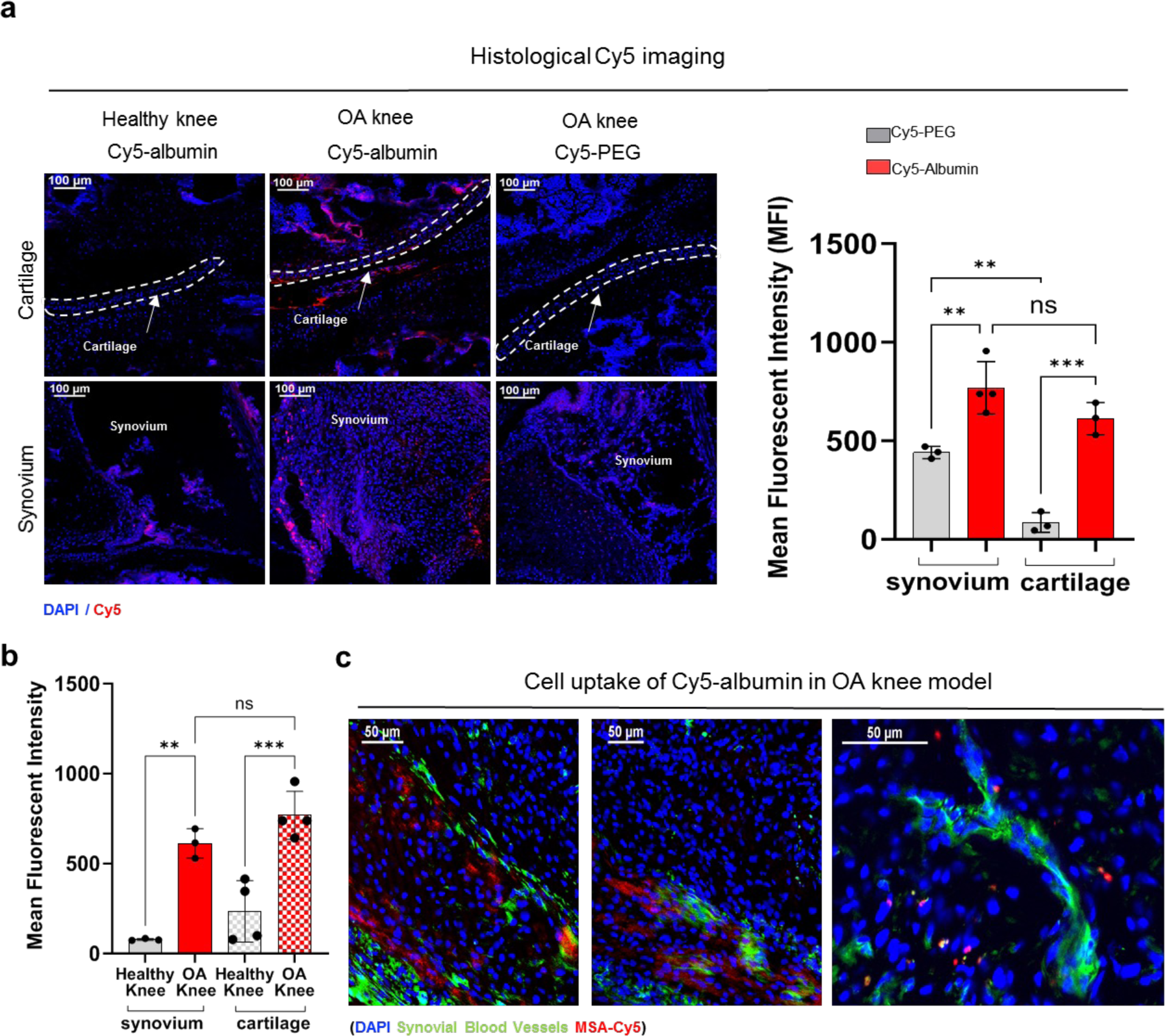
Albumin accumulates in cartilage and synovial tissues of mechanically loaded OA knees to a greater extent than Poly(ethylene glycol). **A)** Representative sagittal cryosections (20X) of knee joints used for assessing MSA-Cy5 (N=5) and PEG-Cy5 (N=3). Fluorescent signal within cartilage and synovial compartments of PTOA mouse knee joints. **B)** Fluorescent signal within cartilage and synovial compartments of PTOA and healthy mouse knee joints treated with MSA-Cy5. **C)** Representative sagittal cryosections (20X) of knee joints assessing MSA-Cy5 and synovial blood vessels. Representative Statistics markers: *P < 0.05, **P < 0.01, ***P < 0.001, ****P < 0.0001. Dashed lines outline articular cartilage.

**Supplementary Figure 2.**
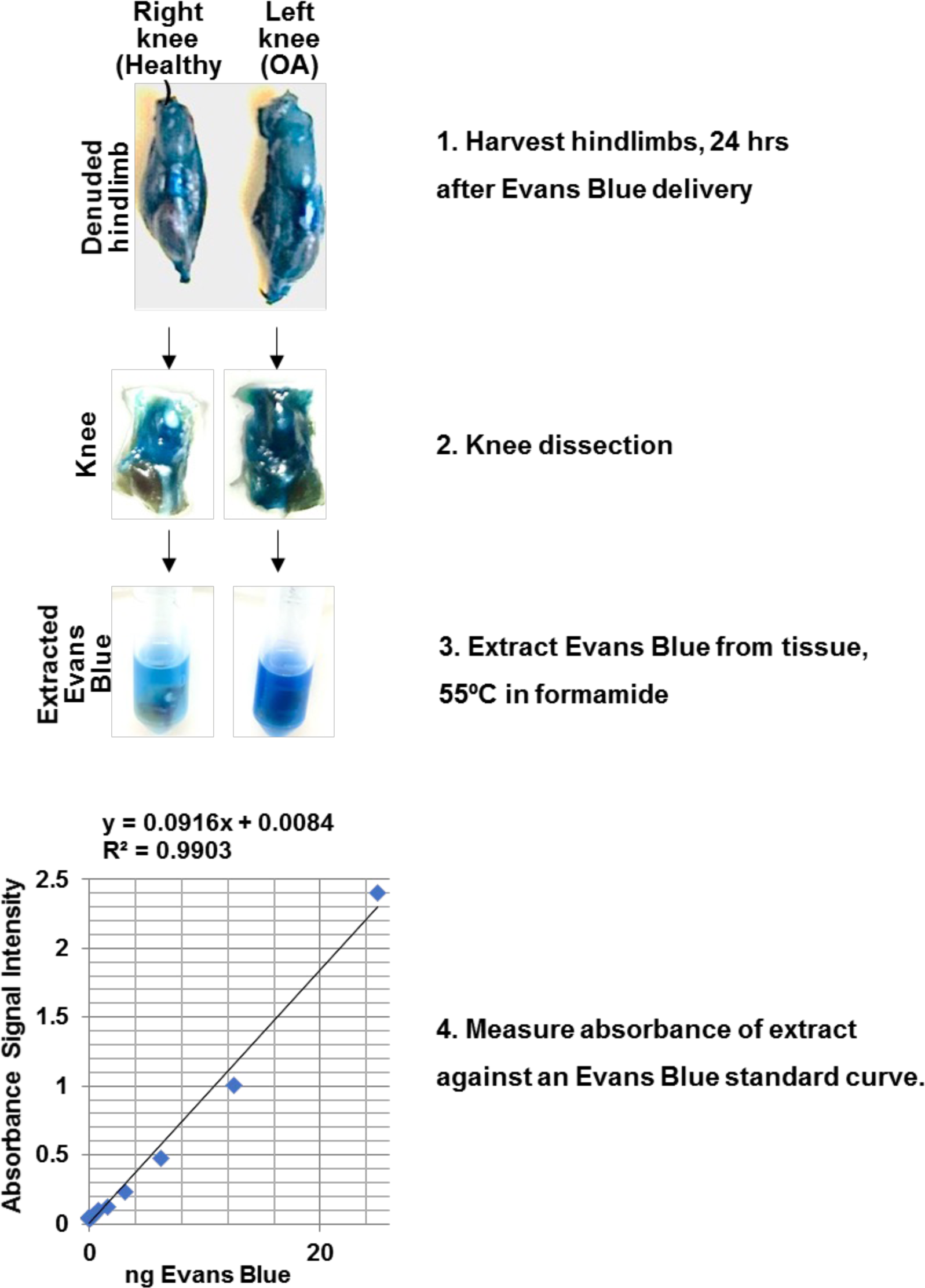
Representative samples and processing protocol for determining Evans Blue ng/mg in explanted tissues.

**Supplementary Figure 3.**
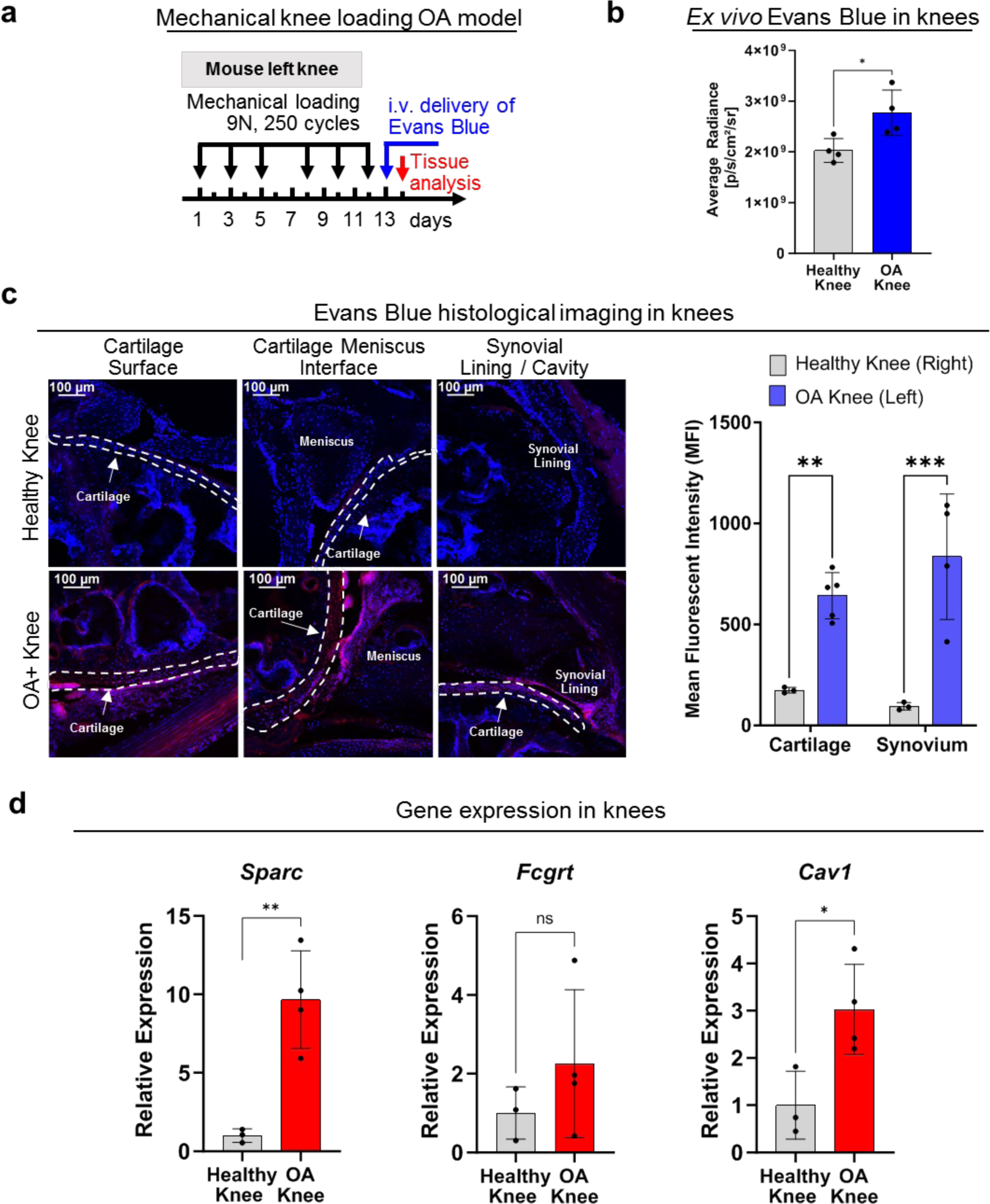
Characterization of albumin delivery/transport in the mechanical loading PTOA model. **A)** Unilateral (left knee loaded, right knee unloaded) mechanical loading protocol/timeline used and injection protocol for Evans Blue dye (N=4). **B)** Intravital (IVIS) quantification of Evans Blue dye. **C)** Representative cryohistology images (20X) of sagittal-sectioned knee joints 24 hours after intravenous Evans Blue injection (Blue = DAPI, Red = Evans Blue). On right is quantification of Evans Blue cryohistology images with a specific focus on cartilage and synovial tissue (N=3-6). **D)** qRT-PCR of cartilage/synovial tissue mix for SPARC, FcRn, Caveolin-1 (N=3-4). Statistics markers: *P < 0.05, **P < 0.01, ***P < 0.001, ****P < 0.0001. Dashed lines outline articular cartilage.

**Supplementary Figure 4.**
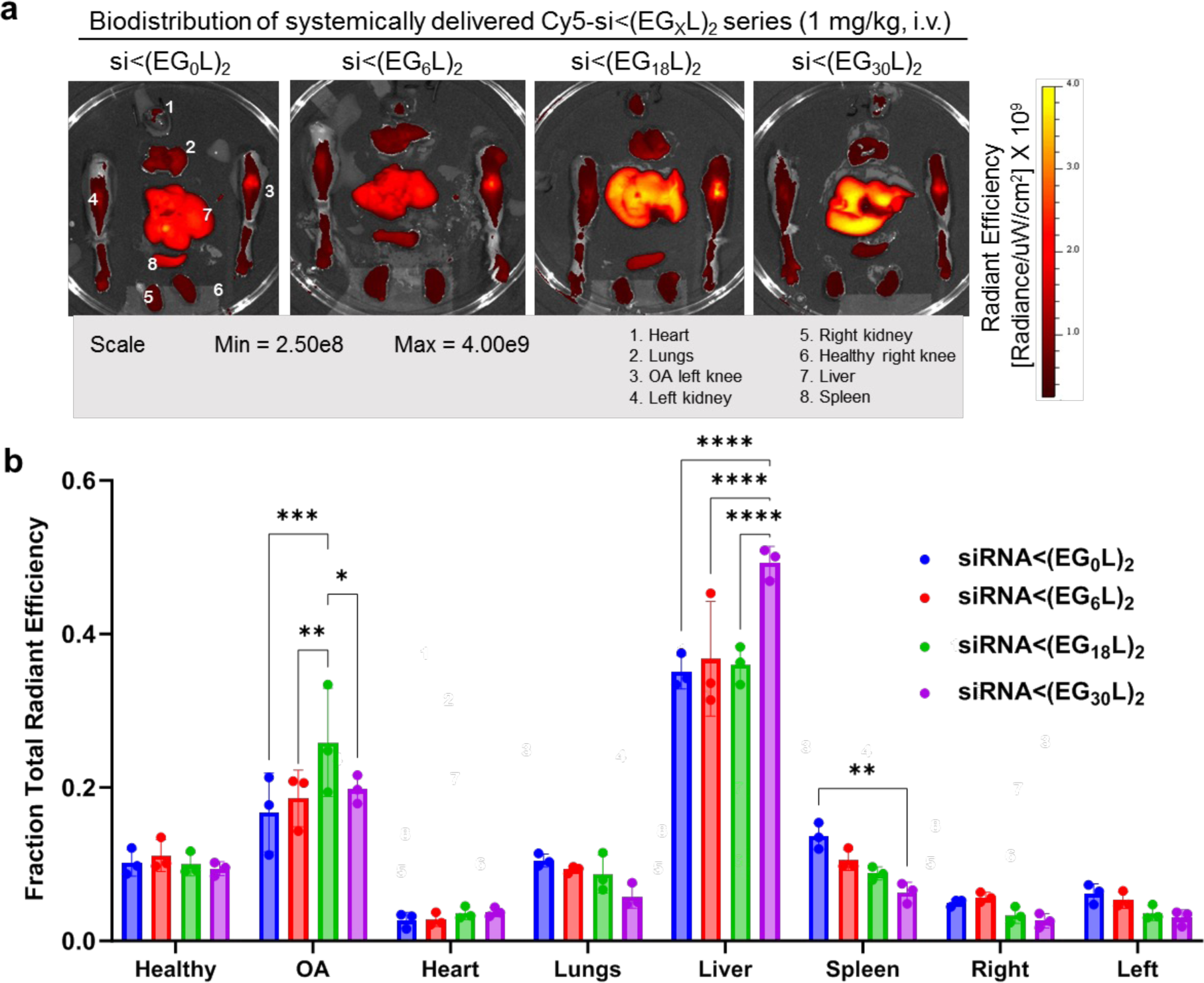
Biodistribution of systemically delivered lipophilic Cy5-si<(EG_X_L)_2_ series in mice with mechanically loaded left knee joints. **A)** Representative ex-vivo IVIS images of Cy5-conjugated siRNAs in organs and limbs. **B)** Organ biodistribution quantification (N=3). Statistics markers: *P < 0.05, **P < 0.01, ***P < 0.001, ****P < 0.0001.

**Supplementary Figure 5.**
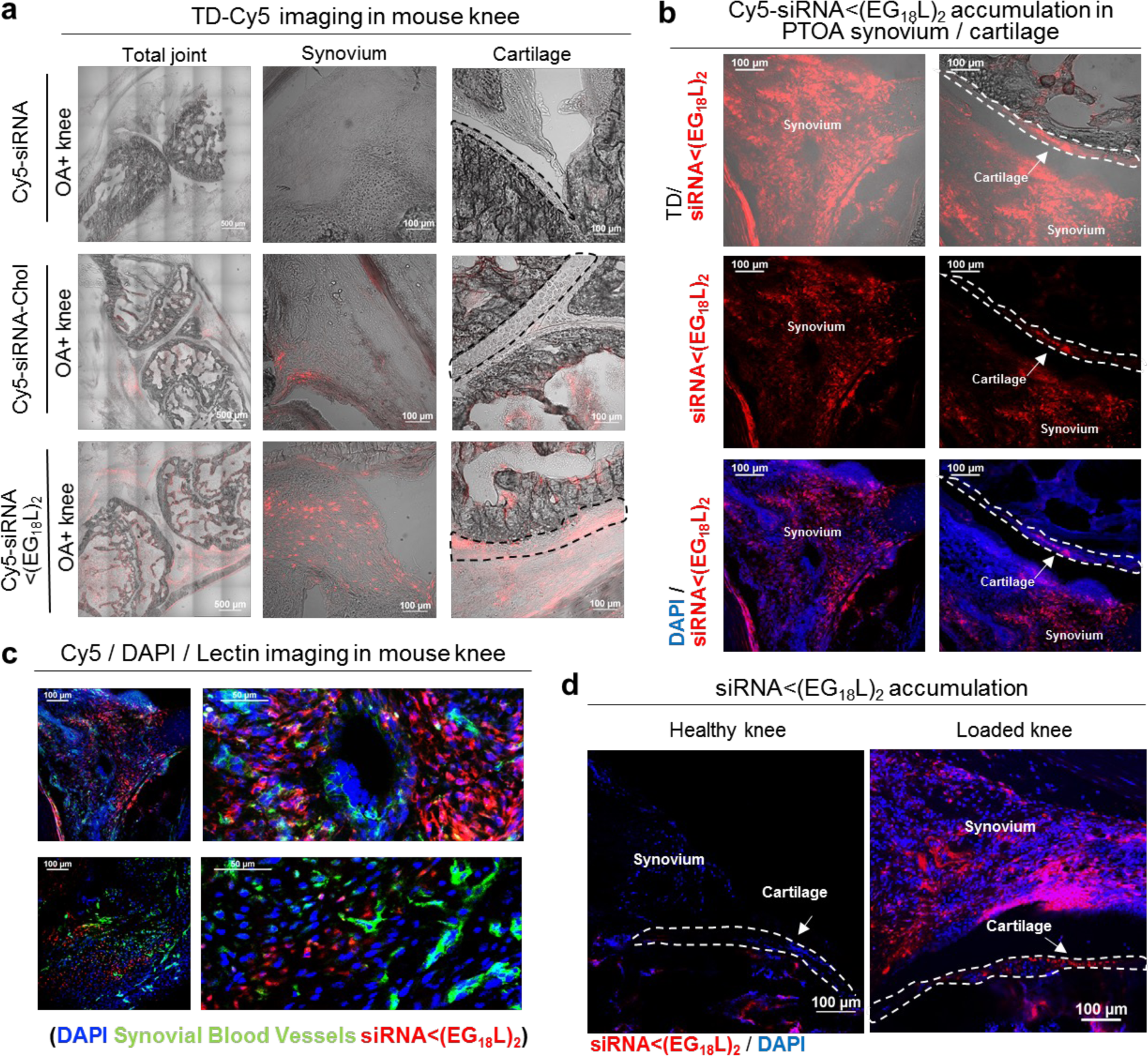
Albumin-hitchhiking siRNA<(EG_18_L)_2_ reaches cartilage and synovial tissues within loaded, arthritic knee joints. **A)** Confocal microscopy of knee joint tissues of mice injected with Cy5-conjugated siRNA molecules – siRNA, siChol, and siRNA<(EG_18_L)_2_. **B)** Additional microscope images of Cy5-siRNA<(EG_18_L)_2_ (red) molecules in synovial and cartilage tissues in loaded, arthritic knee joints. **C)** Confocal microscopy of Cy5-siRNA<(EG_18_L)_2_ (red) in loaded knees stained for blood vessels in the synovium/extra-articular tissues (green). **D)** Microscope images of Cy5-siRNA<(EG_18_L)_2_ (red) molecules in synovial and cartilage tissues in loaded, arthritic knee joints vs. healthy, unloaded knee joints. TD – Transmission Detector. Dashed lines outline articular cartilage.

**Supplementary Figure 6.**
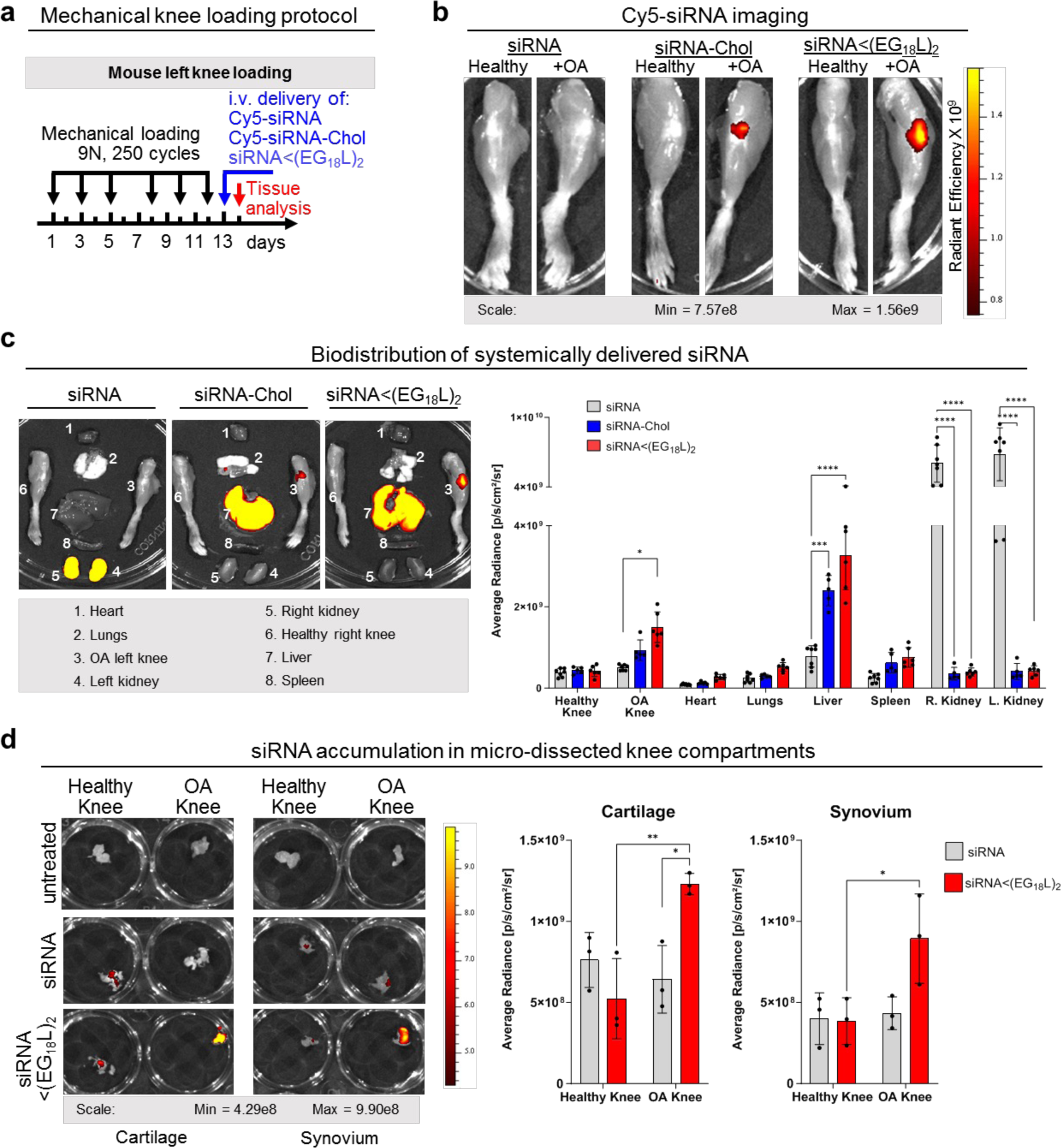
Albumin-hitchhiking siRNA<(EG_18_L)_2_ preferential delivery to loaded, arthritic knee joints. **A)** Unilateral mechanical knee loading protocol/timeline used and injection schedule for systemic delivery of Cy5-conjugated siRNA. **B)** Representative ex-vivo IVIS images. **C)** Organ biodistribution representative images and quantification (N=5-7). **D)** Representative IVIS images of cartilage and synovial tissues dissected out of arthritic and healthy knee joints and quantification of IVIS signal (N=3). Statistics markers: *P < 0.05, **P < 0.01, ***P < 0.001, ****P < 0.0001.

**Supplementary Figure 7.**
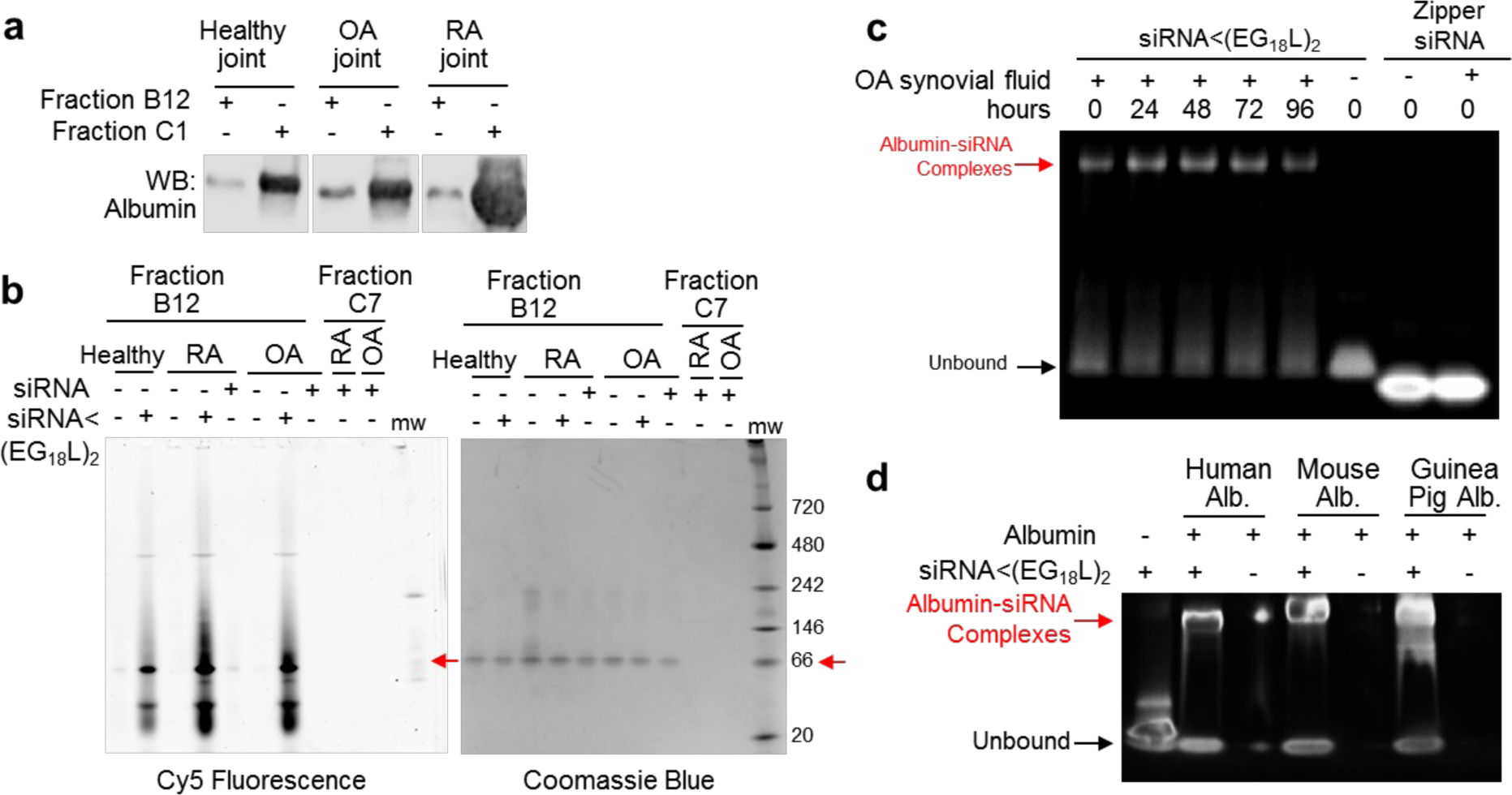
siRNA<(EG_18_L)_2_, but not free siRNA, binds to albumin in human synovial fluid, with increased binding in specimens from OA- and RA-affected joints. A and B are the basis for definition of peaks in the chromatograph in main figure 1. **A)** Western blot for human albumin in B12 and C1 synovial fluid fractions shows increased albumin in OA- and RA-derived specimens. **B)** Fractions B12 and C7 were fractionated by SDS-PAGE, assessing both Cy5 fluorescence (left panel) and total proteins (coomassie blue). Fraction C7 was used as a negative control, non-albumin containing fraction where unbound siRNA elutes. Red arrow indicates molecular weight of human albumin (66 kDa). **C)** Native PAGE of siRNA<(EG_18_L)_2_ sequences after incubation in synovial fluid collected from an OA patient compared to zipper siRNA without a terminal lipid modification. **D)** Gel electrophoresis mobility shift assay demonstrating ability of siRNA<(EG_18_L)_2_ to bind to albumin in a species-independent manner.

**Supplementary Figure 8.**
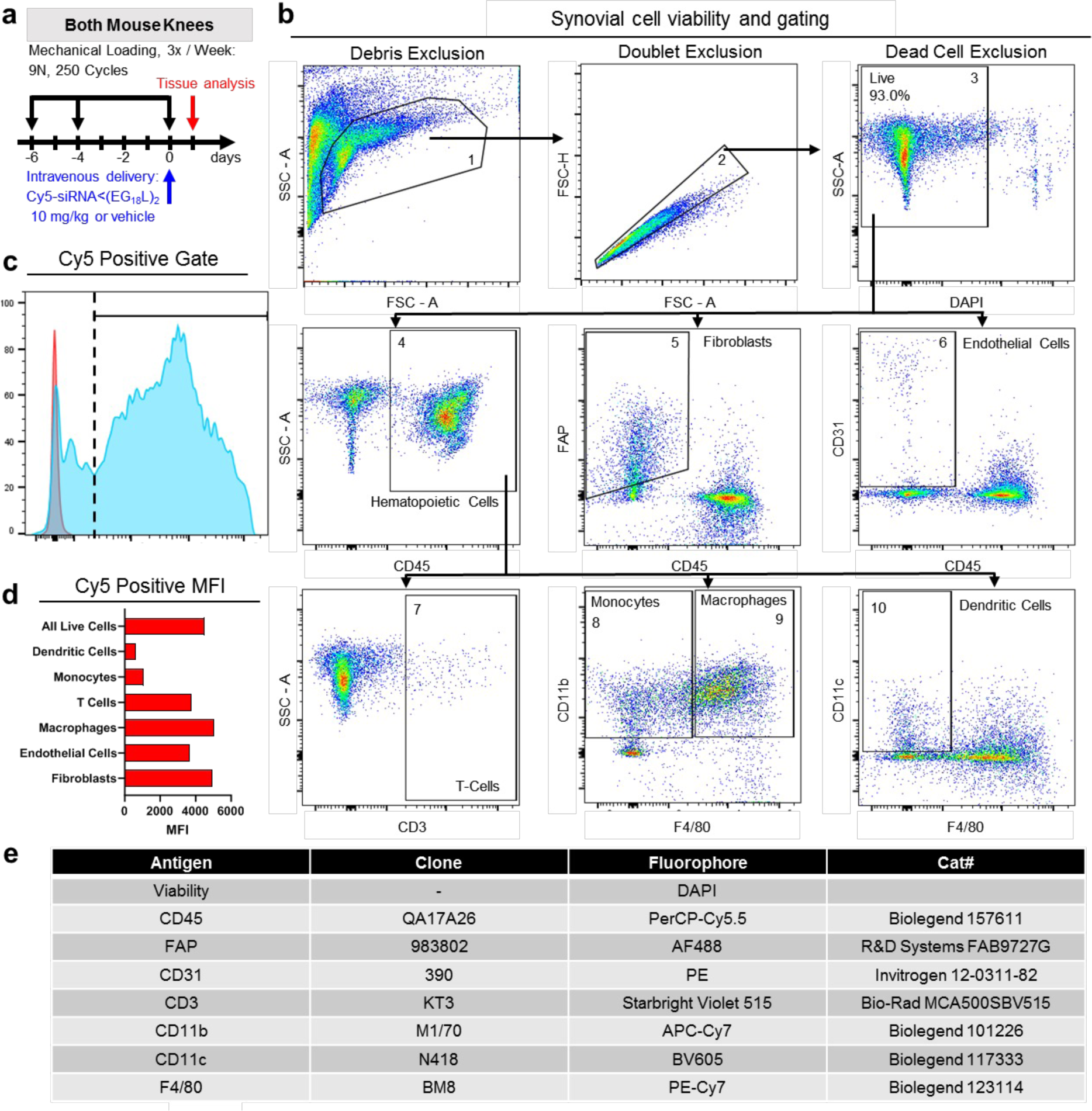
siRNA<(EG_18_L)_2_ exhibits high levels of cellular uptake across several synovial cell types. **A)** Bilateral mechanical knee loading protocol/timeline used and injection schedule for systemic delivery of Cy5-siRNA<(EG_18_L)_2_. **B)** Flow cytometry gating strategies of synovial cells to assess viability and to separate cell populations. Cells were gated on SSC-A and FSC-A for small debris exclusion (1), then single cells gated on FSC-A and FSC-H (2). Dead cell exclusion (3, including percentage of live cells) was performed, and the resulting live single cells were analyzed for cell type. Cells were first gated into hematopoietic cells (4, CD45+), fibroblasts (5, FAP+, CD45-), and endothelial cells (6, CD31+, CD45-). The hematopoietic population (4), was then refined to T-cells (7, CD3+, CD45+), monocytes (8, CD11b+, F4/80-, CD45+), macrophages (9, CD11b+, F4/80+, CD45+), and dendritic cells (10, CD11c+, F4/80-, CD45+). **C)** All single live cells gated on Cy5, indicating siRNA<(EG_18_L)_2_ uptake. **D)** Mean fluorescence intensity (MFI) of Cy5 positive cells **E)** Table of antibody clones and fluorophores used in flow cytometry experiment.

**Supplementary Figure 9.**
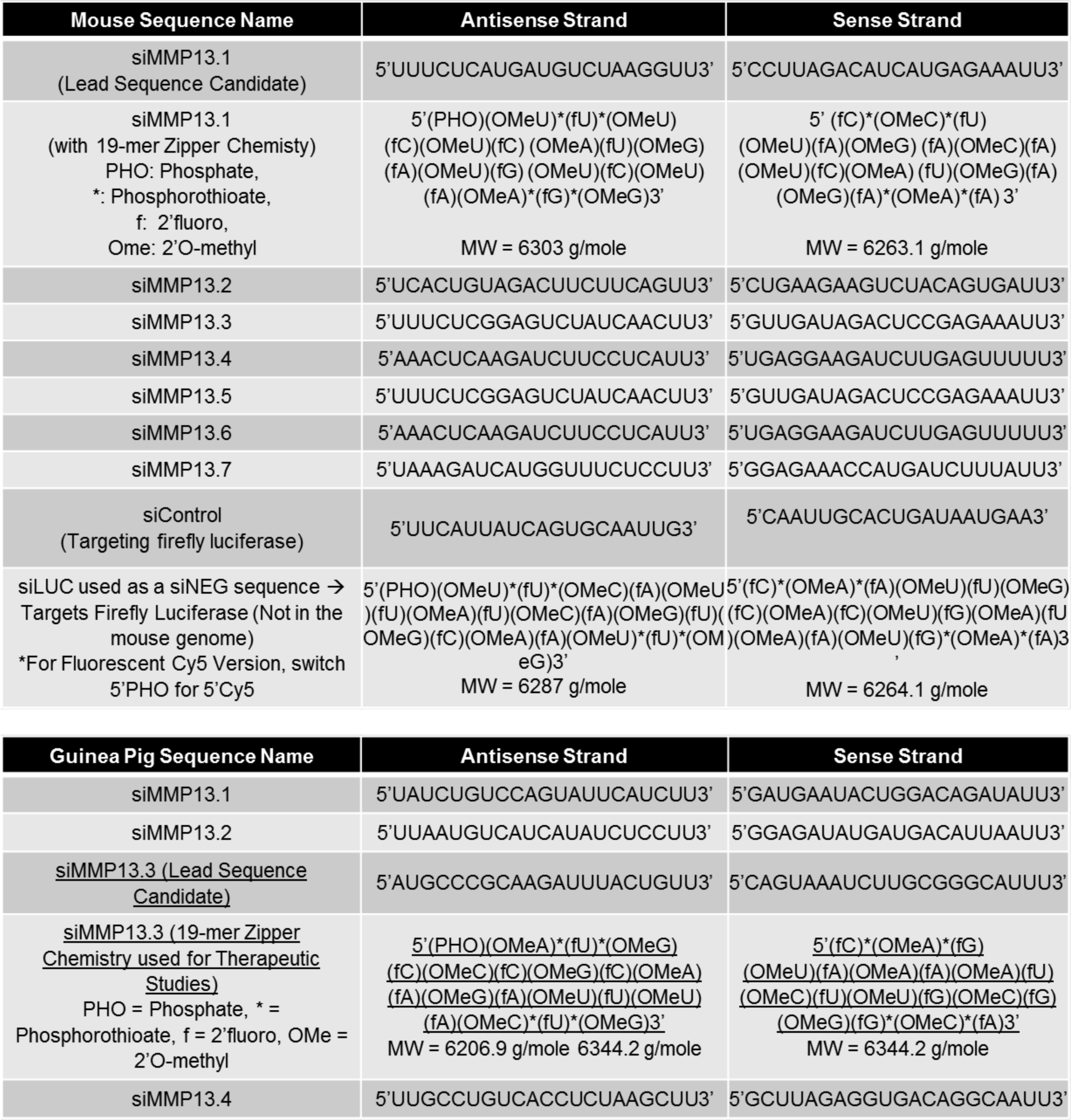
Mouse and guinea pig siRNA sequences and chemical modifications.

**Supplementary Figure 10.**
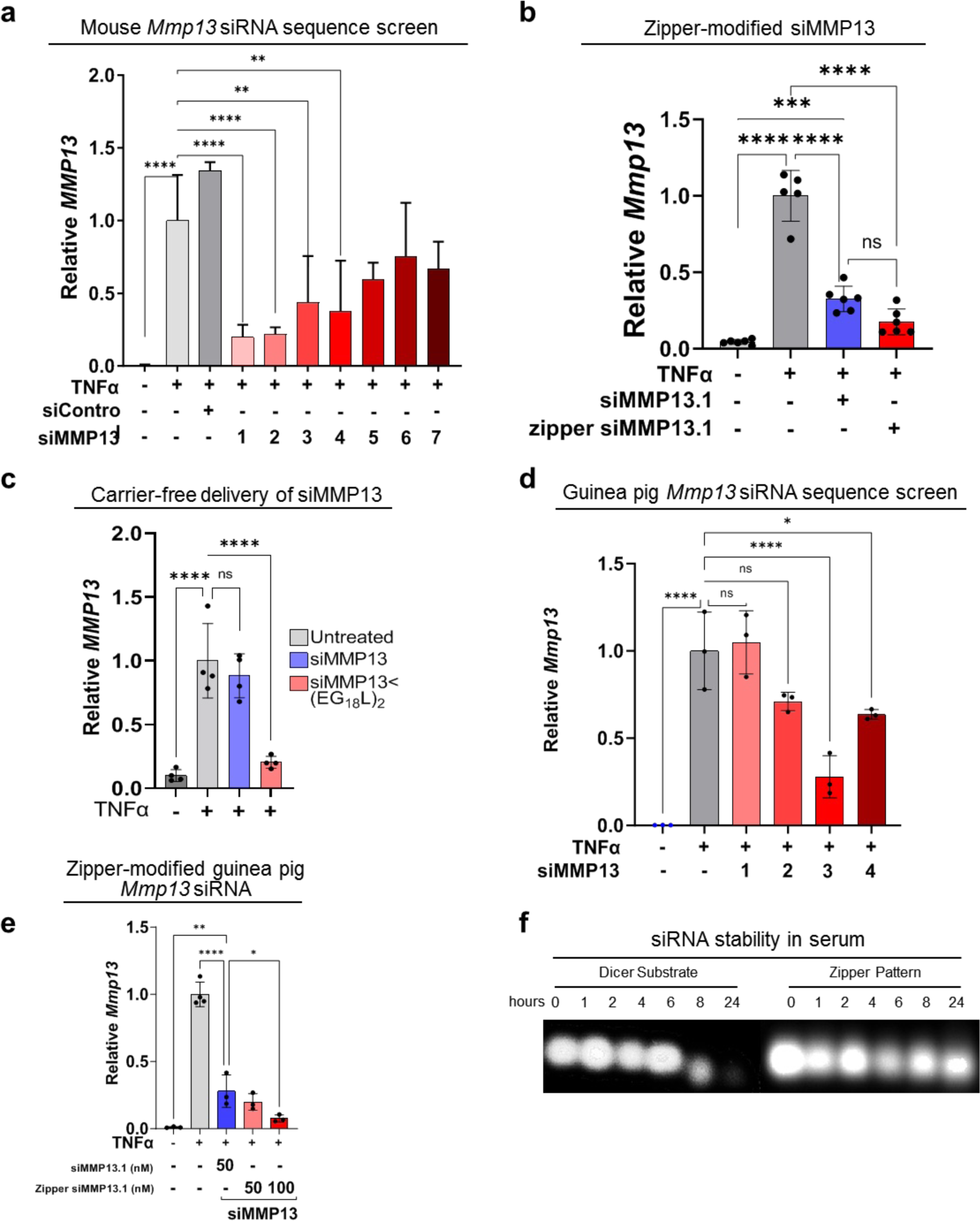
MMP13 siRNA sequence screening across species, carrier free activity, and stability. **A-C)** *Mmp13* mRNA was measured in TNFα-stimulated murine chondrogenic ATDC5 cells by RT-qPCR following transfection of candidate murine MMP13 siRNA sequences (**A**); Lipofectamine-mediated delivery of zipper-modified relative to unmodified siMMP13 sequences (**B**); Carrier-free silencing activity of siMMP13<(EG_18_L)_2_ (**C**). (N=3-4). **D-E)** *Mmp13* mRNA was measured in TNFα-stimulated primary guinea pig chondrocytes by RT-qPCR following lipofectamine-mediated delivery of guinea pig MMP13 siRNA sequences (**D**) or zipper-modified guinea pig siMMP13 (**E**). **F)** Serum stability (60% FBS, fetal bovine serum) at 37°C of zipper modified or Dicer substrate mouse siMMP13. Statistics markers: *P < 0.05, **P < 0.01, ***P < 0.001, ****P < 0.0001.

**Supplementary Figure 11.**
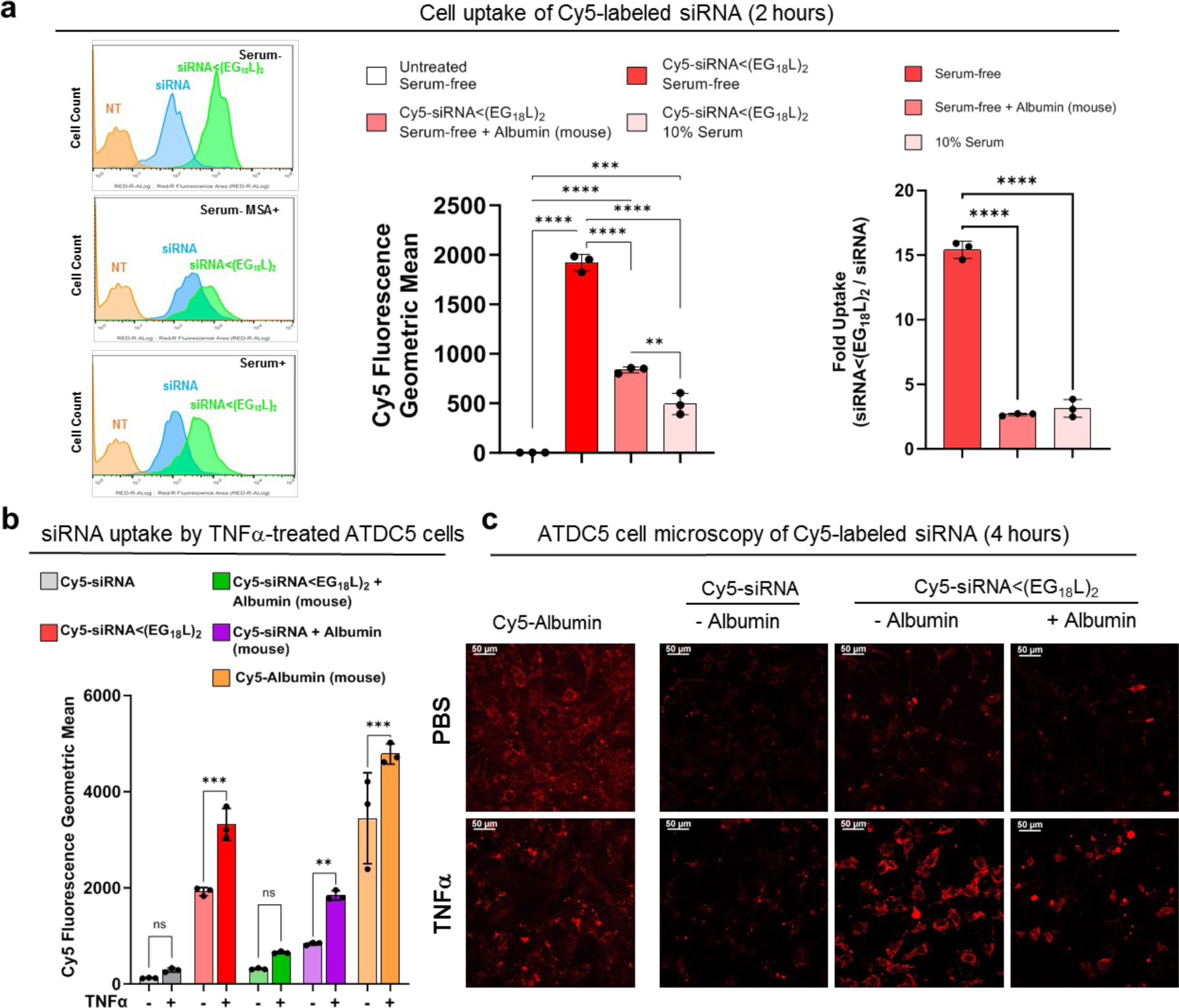
Carrier-Free cell uptake in chondrogenic murine ATDC5 cells. **A)** Flow cytometry histograms of cellular uptake 2 hours after carrier-free treatment of Cy5-conjugated siRNA or siRNA<(EG_18_L)_2_ at a concentration of 100 nM in serum-free, serum-free + supplemented mouse serum albumin (10x, 1 μM), or serum-containing media. Data are quantified as fold change of siRNA<(EG_18_L)_2_/siRNA cell uptake in differing media conditions (N=3). **B)** Quantification of cellular uptake by flow cytometry in ATDC5 cells treated with or without murine TNFα (20 ng/mL) (N=3). **C)** Representative confocal microscopy images of ATDC5 cellular uptake (Cy5 fluorescence) after 4 hours of treatment with or without stimulation with murine TNFα (20 ng/mL). Statistics markers: *P < 0.05, **P < 0.01, ***P < 0.001, ****P < 0.0001.

**Supplementary Figure 12.**
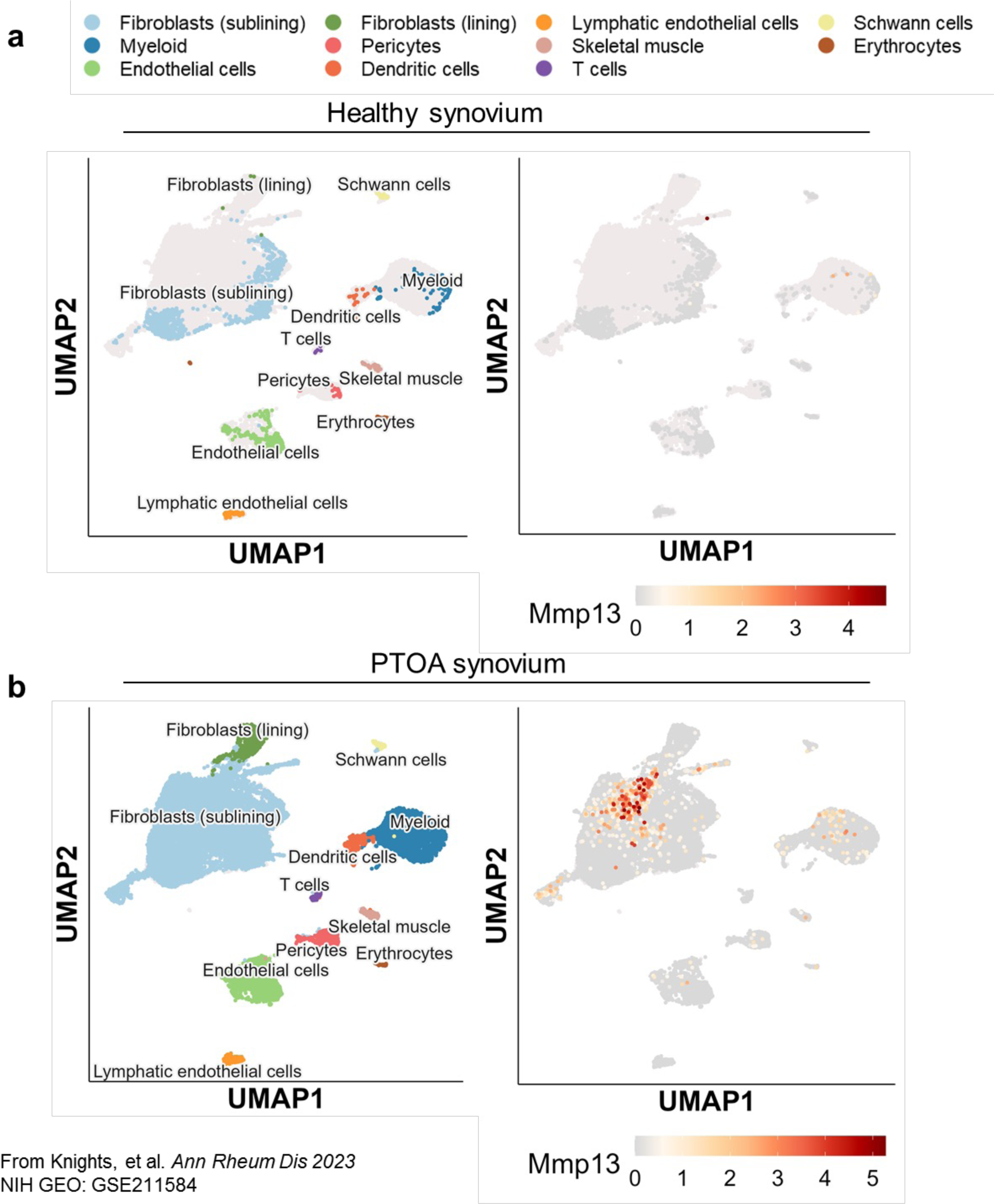
Mmp13 is upregulated in synovial fibroblasts and myeloid cells. **A)** Distribution of cell populations in a healthy mouse synovium (left) with low levels of Mmp13 expression (right). **B)** Distribution of cell populations in a noninvasive ACL rupture model PTOA mouse synovium (left) with increased Mmp13 expression (right)

**Supplementary Figure 13.**
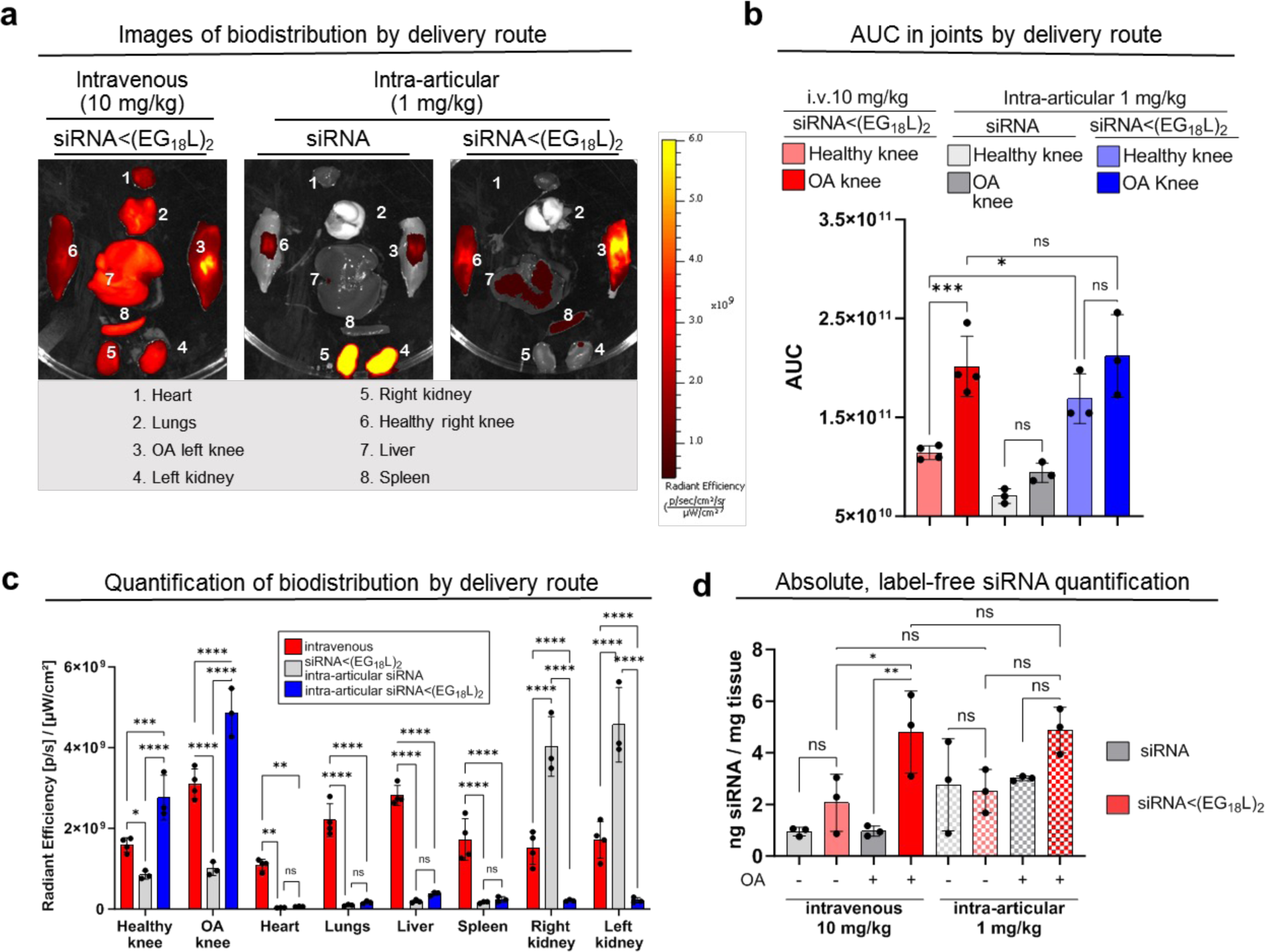
Pharmacokinetic and biodistribution analyses of intra-venous vs. intra-articular routes. **A)** Representative IVIS organ biodistribution images of all 3 groups at the 30-day endpoint. **B)** AUC analysis for i.v. siRNA<(EG_18_L)_2_ (10 mg/kg), intra-articular siRNA (1 mg/kg), and Intra-articular siRNA<(EG_18_L)_2_ (1 mg/kg) throughout the entire 30-day intravital IVIS retention study (N=3-4). **C)** Quantification of organ biodistribution from images in A (N=3-4). **D)** PNA hybridization assay showing ng/mg antisense strand of the siRNA per mg of knee joint tissue at 24 hrs (i.v.) or 48 hrs (i.a.) (N=3). Statistics markers: *P < 0.05, **P < 0.01, ***P < 0.001, ****P < 0.0001.

**Supplementary Figure 14.**
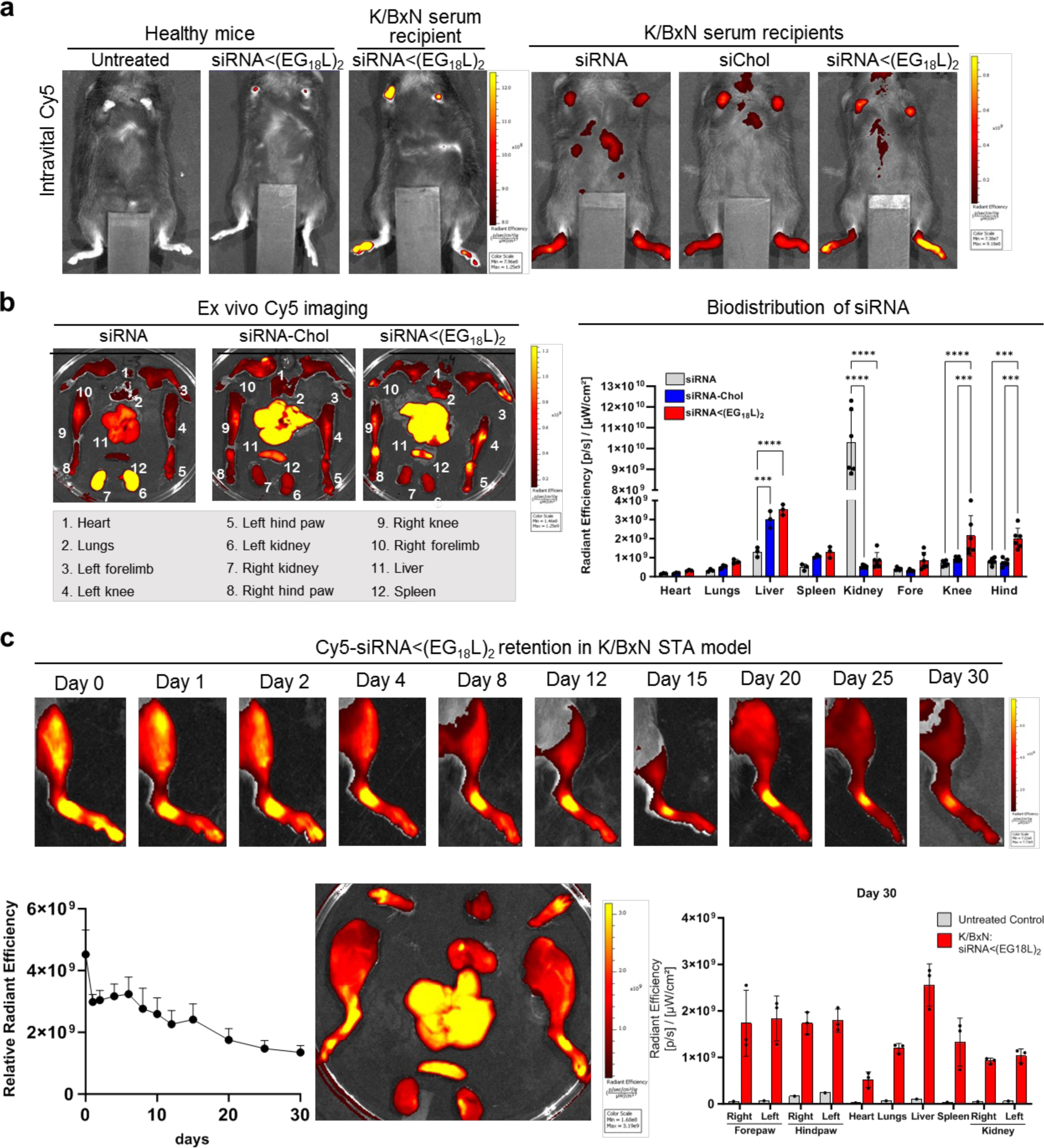
Biodistribution and pharmacokinetics of K/BxN STA model. **A-B)** Cy5-siRNA<(EG_18_L)_2_, Cy5-siRNA-Chol, or Cy5-siRNA was delivered i.v. (1 mg/kg) to healthy and K/BxN STA mice and assessed 24 hrs later. Intravital Cy5 measurements in healthy and K/BxN STA mice in each treatment group (A). Organ Cy5 biodistribution was measured ex vivo (B) (N=3 mice). **C)** Longevity of Cy5-siRNA<(EG_18_L)_2_ retention in PTOA knees following single dose of Cy5-siRNA<(EG_18_L)_2_ (10mg/kg, i.v.) was assessed by longitudinal intravital imaging, and endpoint imaging of knees and organs done ex vivo on day 30 (N=3 mice). Radiant efficiency over time displayed as mean + SEM. Statistics markers: *P < 0.05, **P < 0.01, ***P < 0.001, ****P < 0.0001.

**Supplementary Figure 15.**
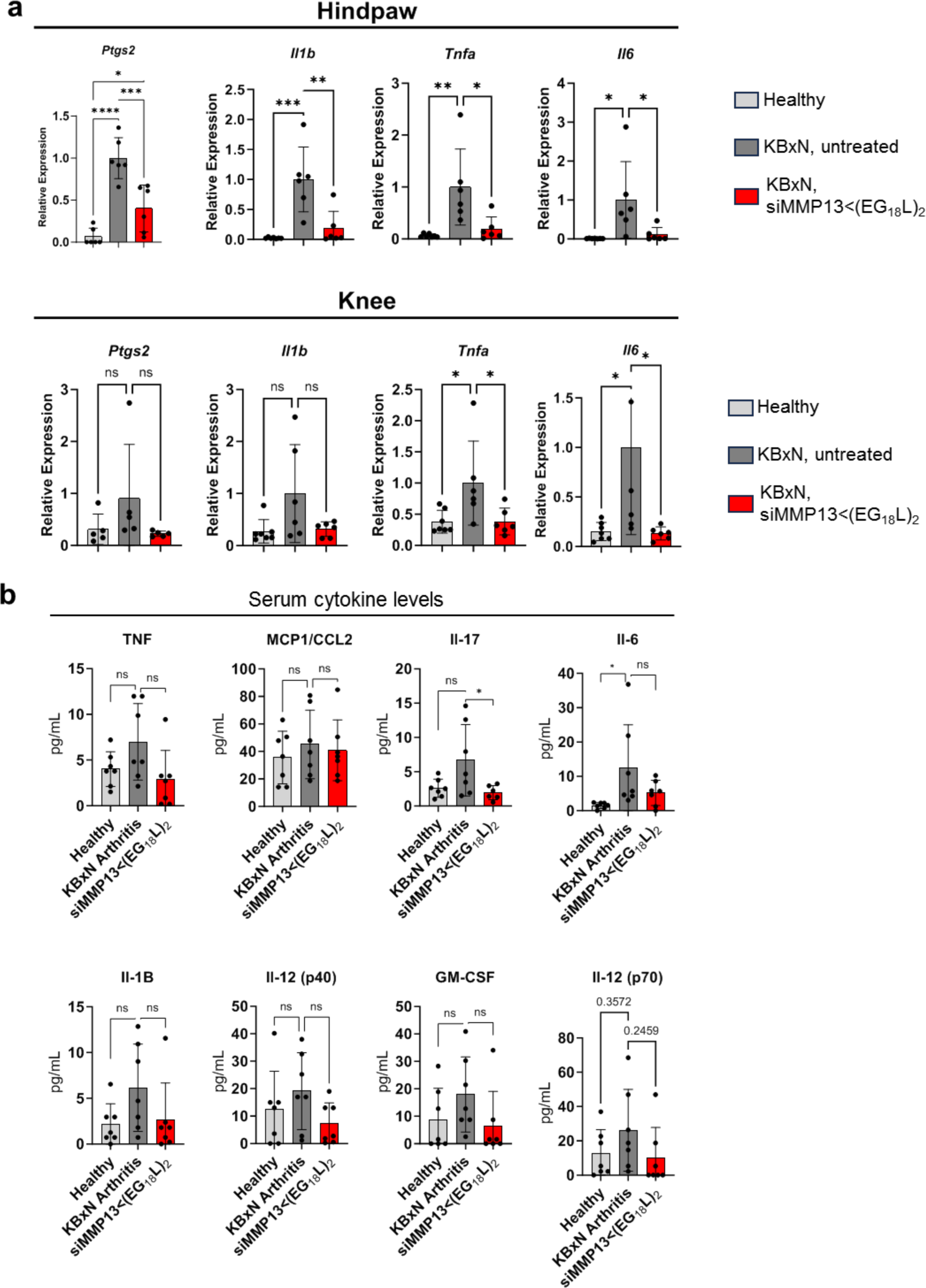
Molecular analysis of K/BxN therapeutic study. **A)** qRT-PCR of genes of importance in rheumatoid arthritis in the K/BxN therapeutic study at the day 10 endpoint (N=5-7). **B)** Multiplex Luminex analyses for analytes in serum in the K/BxN therapeutic study at the day 10 endpoint (N=5-7). Statistics markers: *P < 0.05, **P < 0.01, ***P < 0.001, ****P < 0.0001.

**Supplementary Figure 16.**
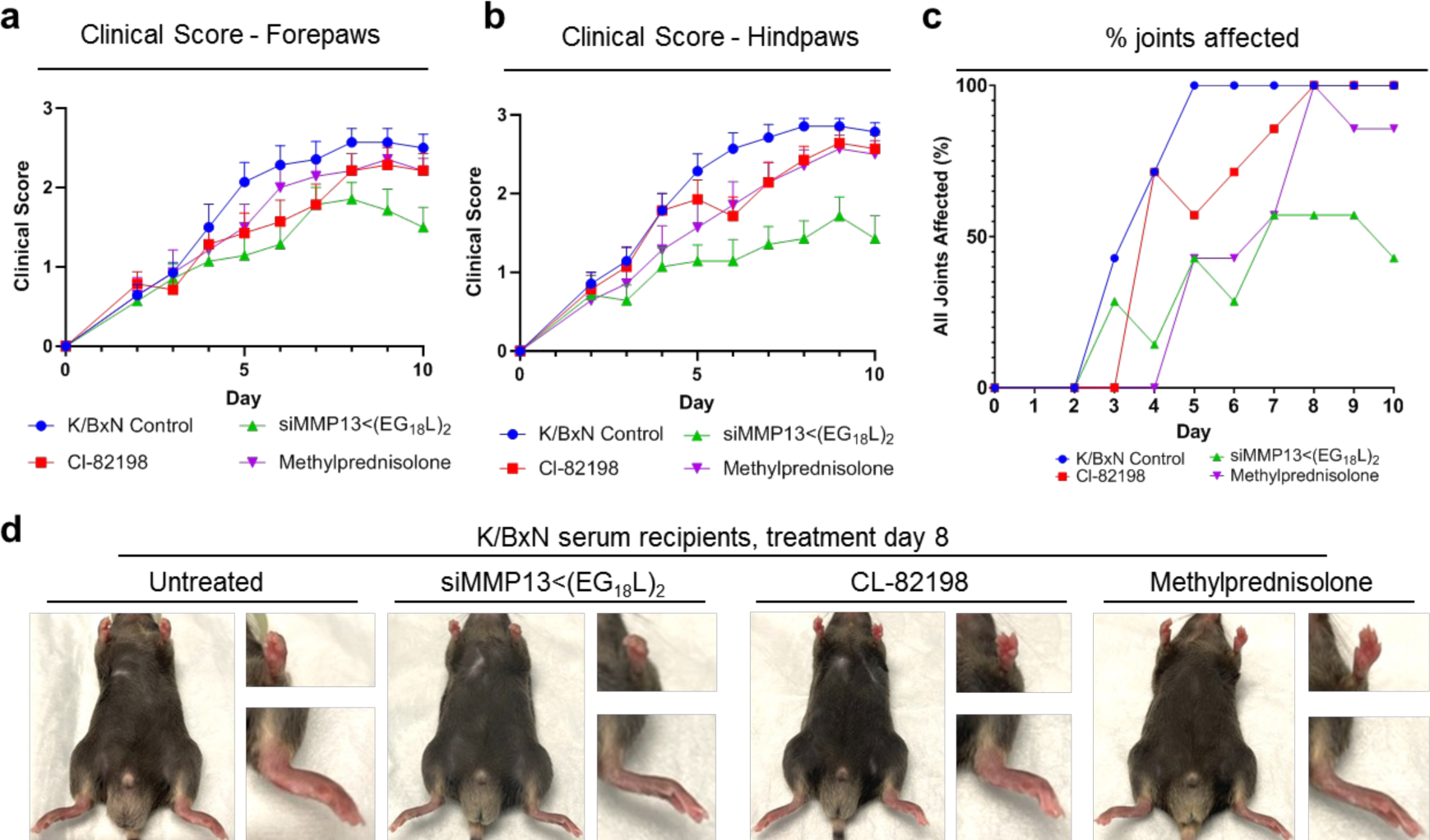
Additional paw clinical score analysis of mice in K/BxN therapeutic study. **A-B)** Clinical score of forepaws (**A**) and hindpaws (**B**) (N=7 mice). Data displayed as mean + SEM. **C)** Percentage of mice exhibiting affliction of all paw joints following K/BxN serum transfer (N=7 mice). **D)** Representative images showing inflammation / swelling in limbs of K/BxN serum recipients on treatment day 8 (10 days after serum transfer). Insets show higher power image of forepaws (upper inset) and hindpaws (lower inset). Images were used for clinical scoring, with a maximal clinical score of 3 per paw, total score of 12 per mouse.

**Supplementary Figure 17.**
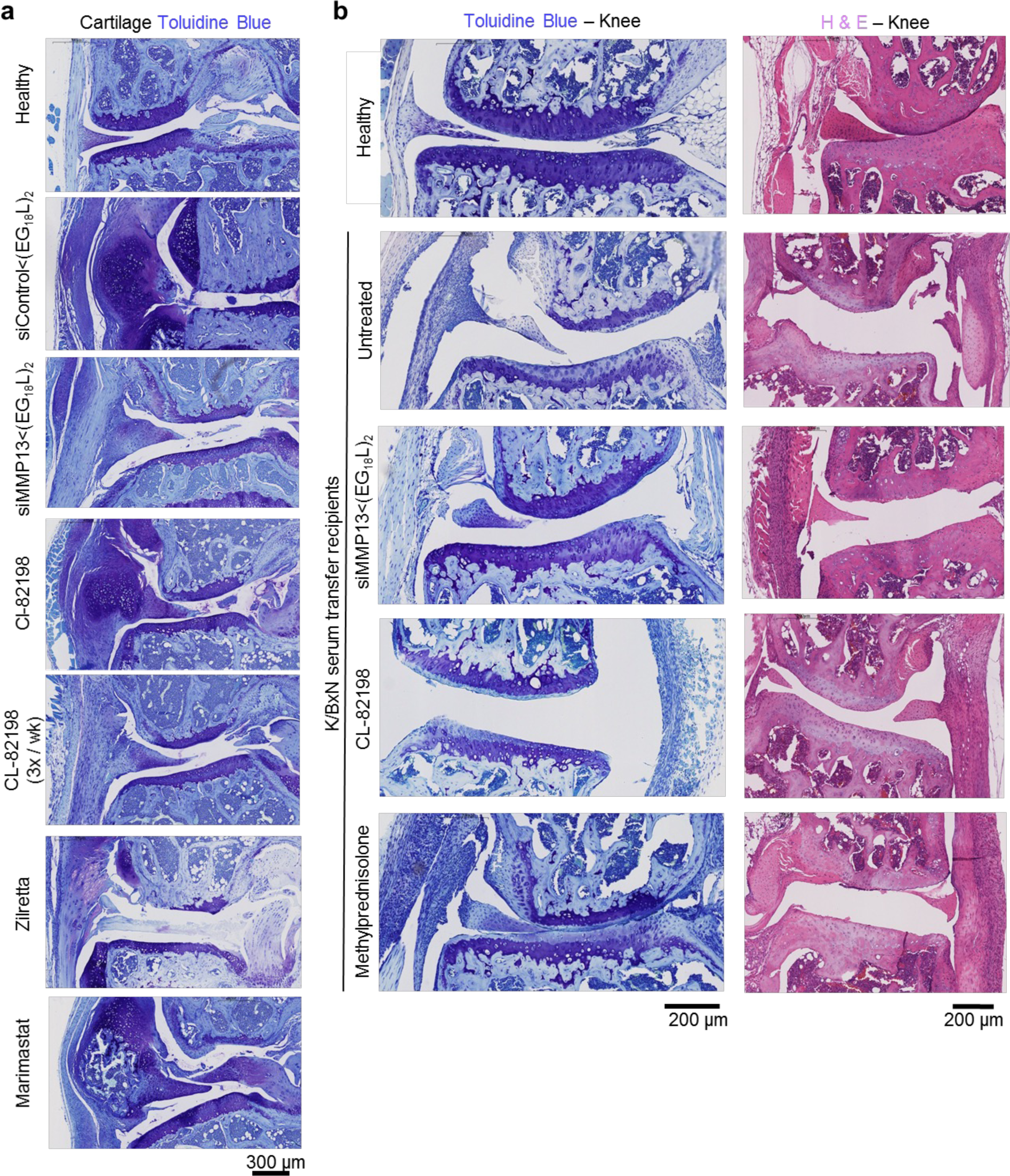
Representative lower magnification knee joint histology for PTOA and K/BxN therapeutic studies. **A)** Lower magnification of medial joint toluidine blue histology for the PTOA therapeutic study. **B)** Toluidine blue and H&E of knee joint tissue for the K/BxN therapeutic study.

**Supplementary Figure 18.**
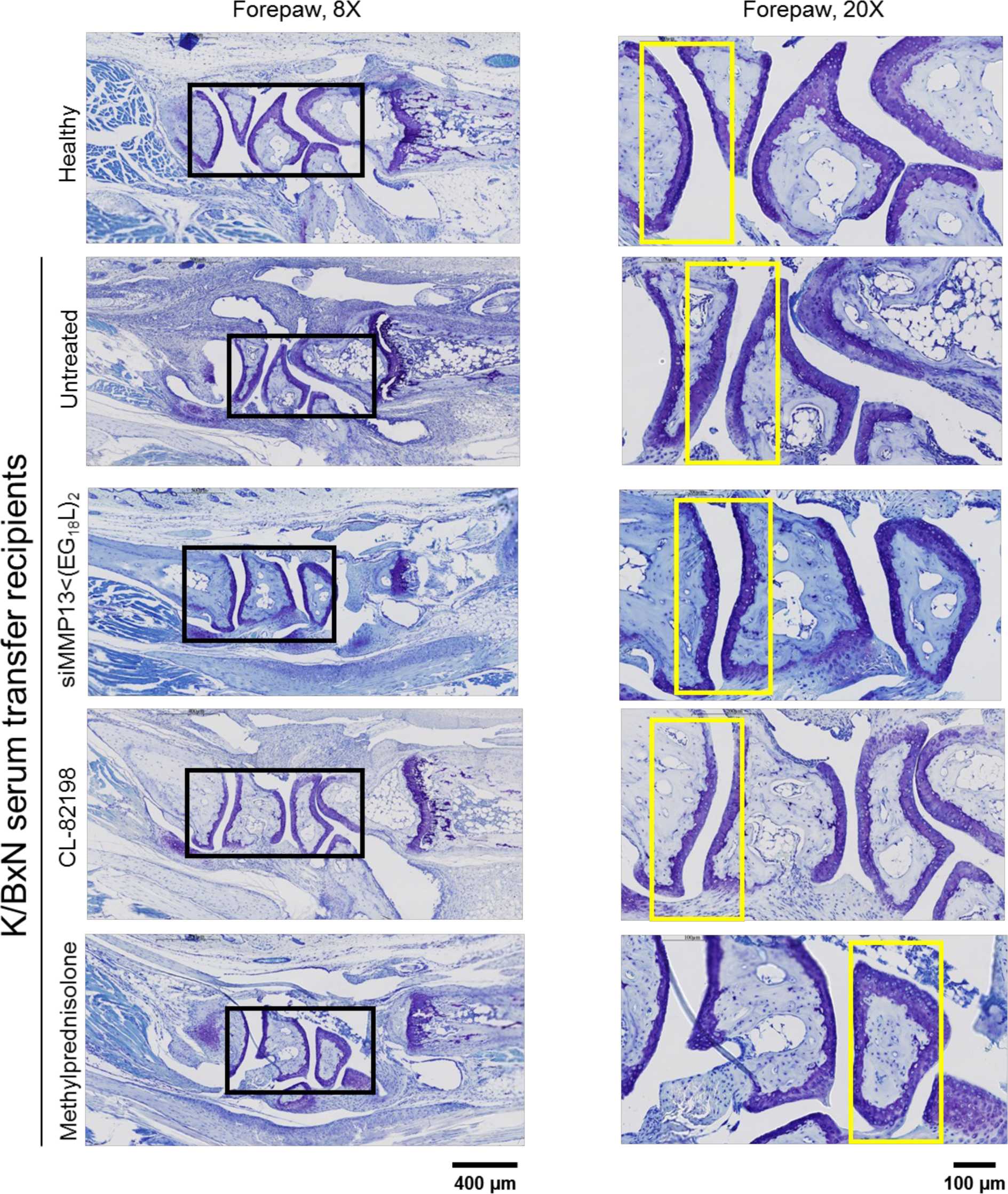
Toluidine Blue of forepaw / wrist cartilage of K/BxN serum recipients. Panels on left show 8X magnification. Black rectangles in left panels are the areas shown at higher magnification (20X) in right panels. Yellow rectangles in right panels indicate the areas of higher magnification shown in extended data figures.

**Supplementary Figure 19.**
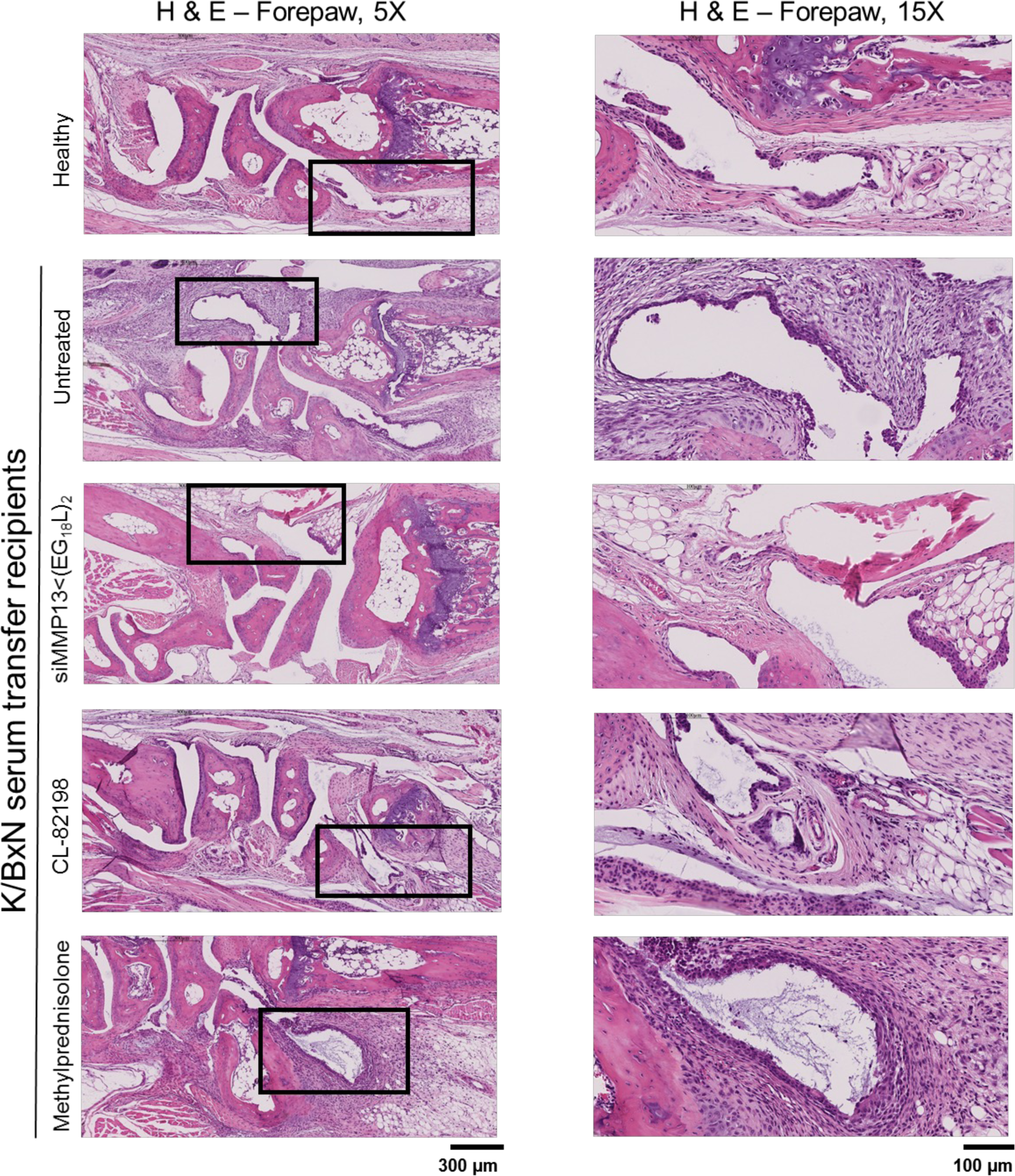
H&E of forepaw/wrist tissue for the K/BxN therapeutic study.

**Supplementary Figure 20.**
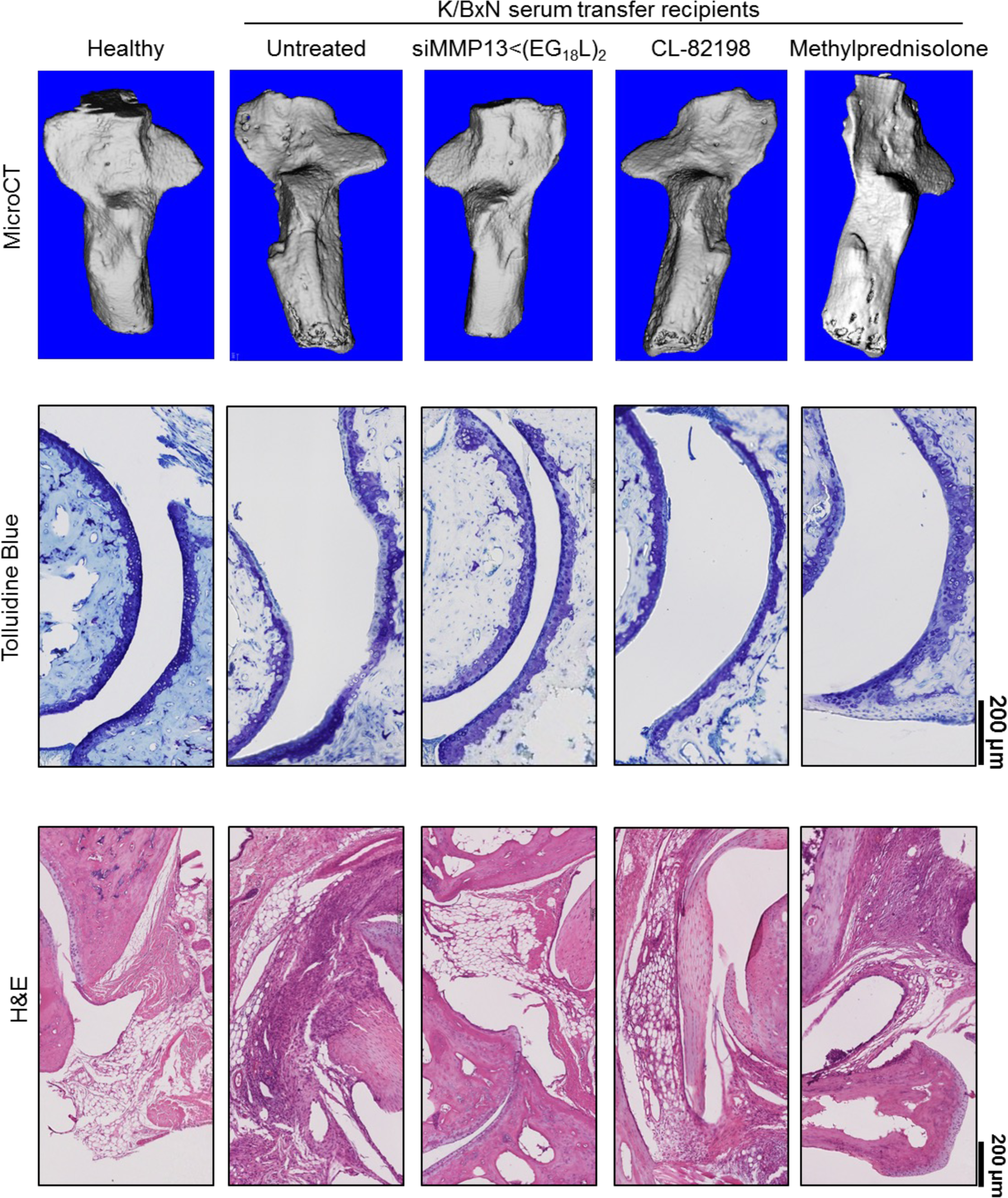
Bone loss and ankle joint assessments in K/BxN therapeutic study. Top: 3D reconstructions of the calcaneus bone (used for microCT parameter quantification). Middle: lower magnification Toluidine Blue of hindpaw/ankle cartilage. Bottom: lower magnification H&E of hindpaw/ankle tissue.

**Supplementary Figure 21.**
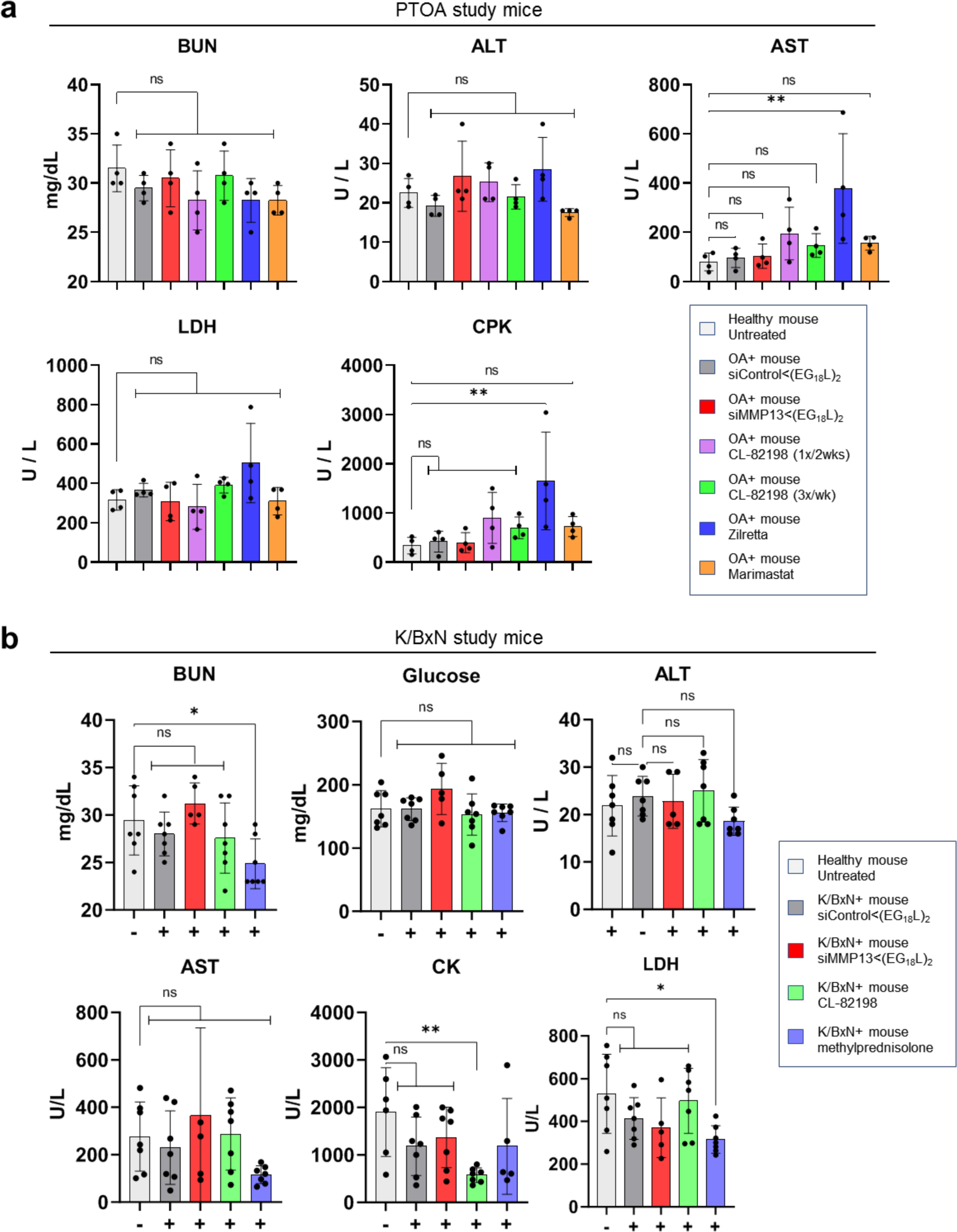
Systemic toxicity analyses in K/BxN and PTOA model therapeutic studies. **A)** Toxicology analyses in serum in the PTOA therapeutic study at the endpoint (N=4 mice). **B)** Toxicology analyses in serum in the K/BxN therapeutic study at the endpoint (N=7 mice). Statistics markers: *P < 0.05, **P < 0.01, ***P < 0.001, ****P < 0.0001.

**Supplementary Figure 22.**
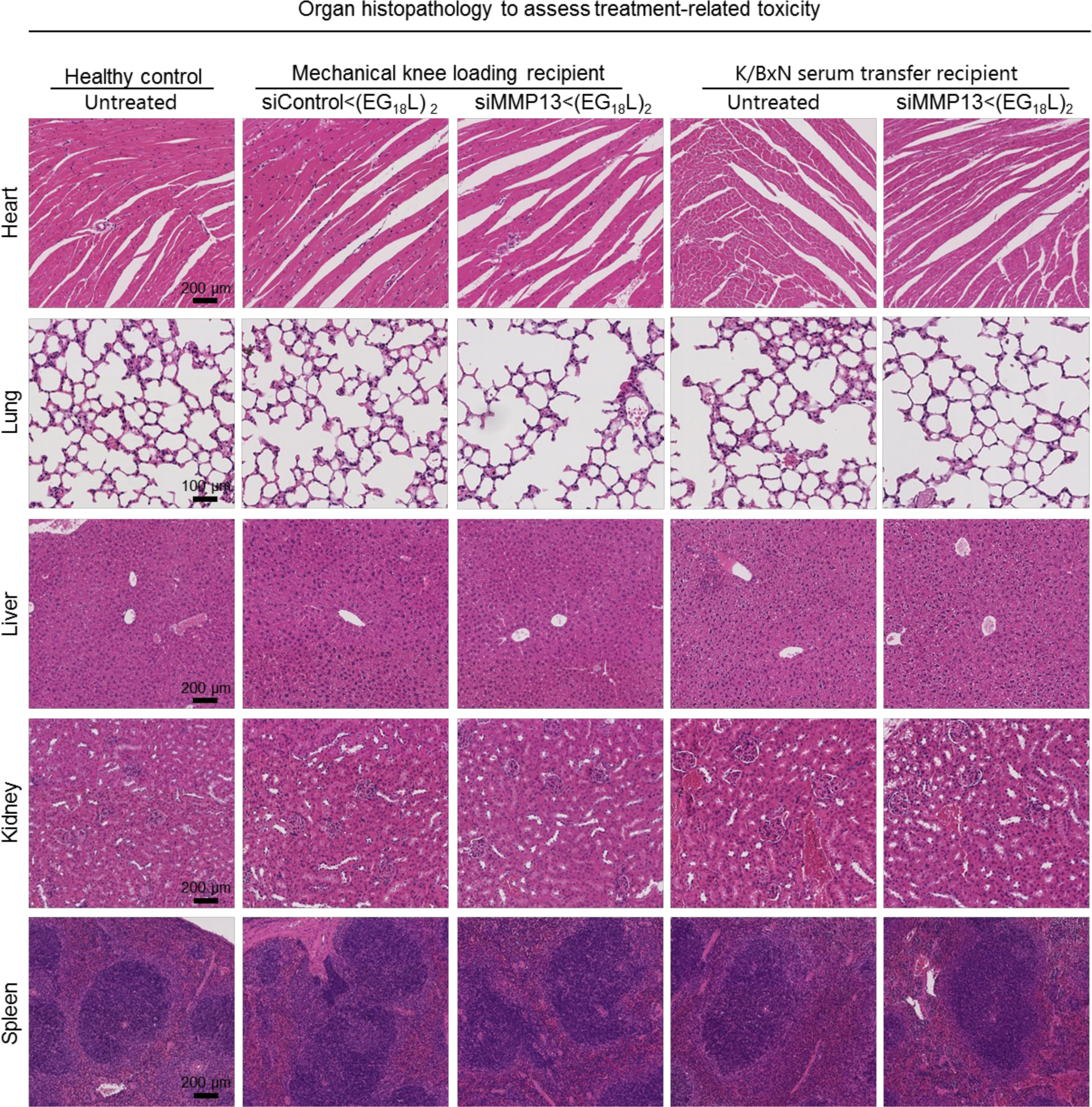
H&E-stained sections of organs (liver, kidney, lung, heart, and spleen) from mice enrolled in the PTOA and K/BxN therapeutic studies. Organs were harvested at the end of the study for analysis, and a healthy (no disease, no treatment) mouse is included as a control.

**Supplementary Figure 23.**
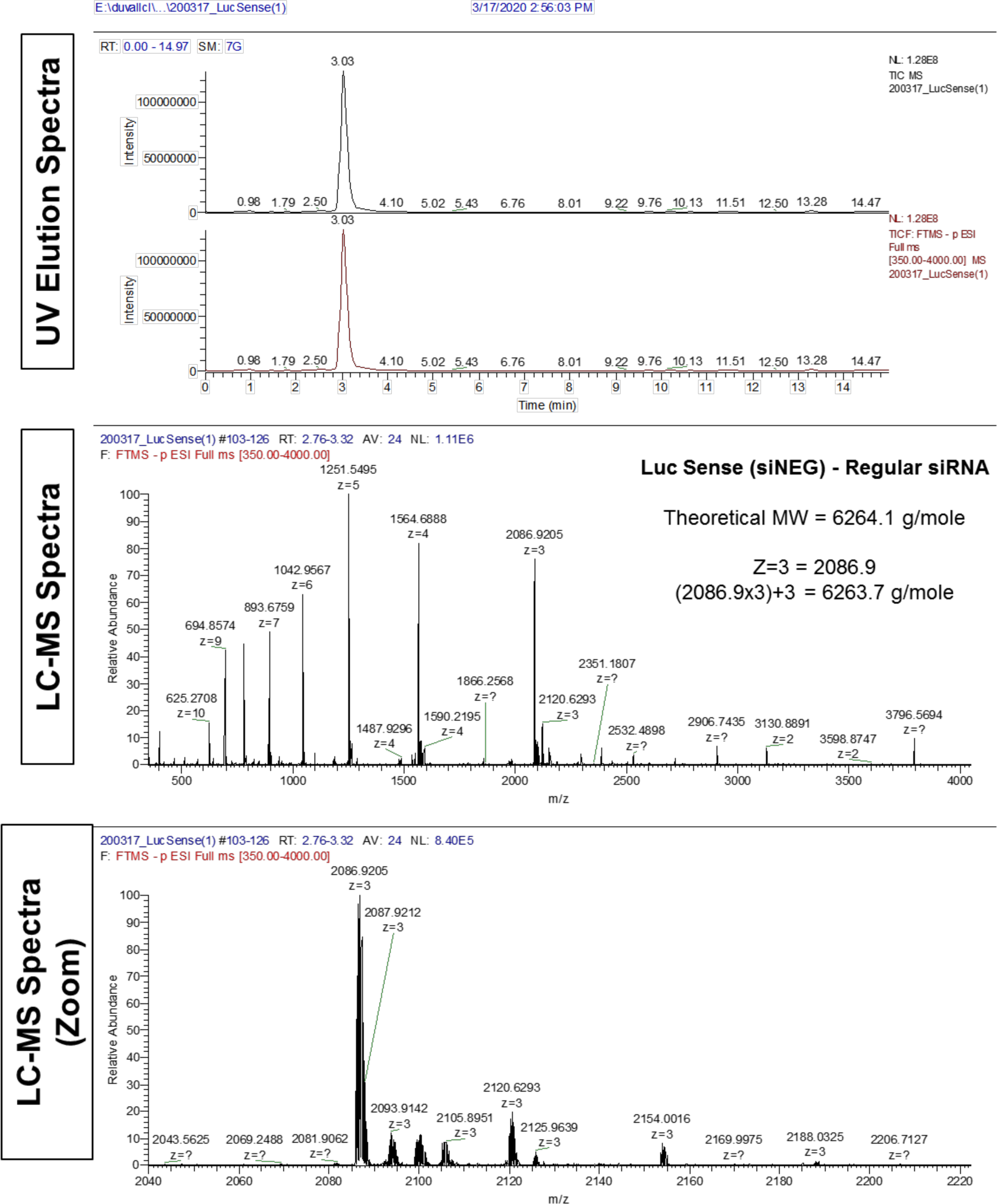
LC-MS characterization of control sense siRNA sequence.

**Supplementary Figure 24.**
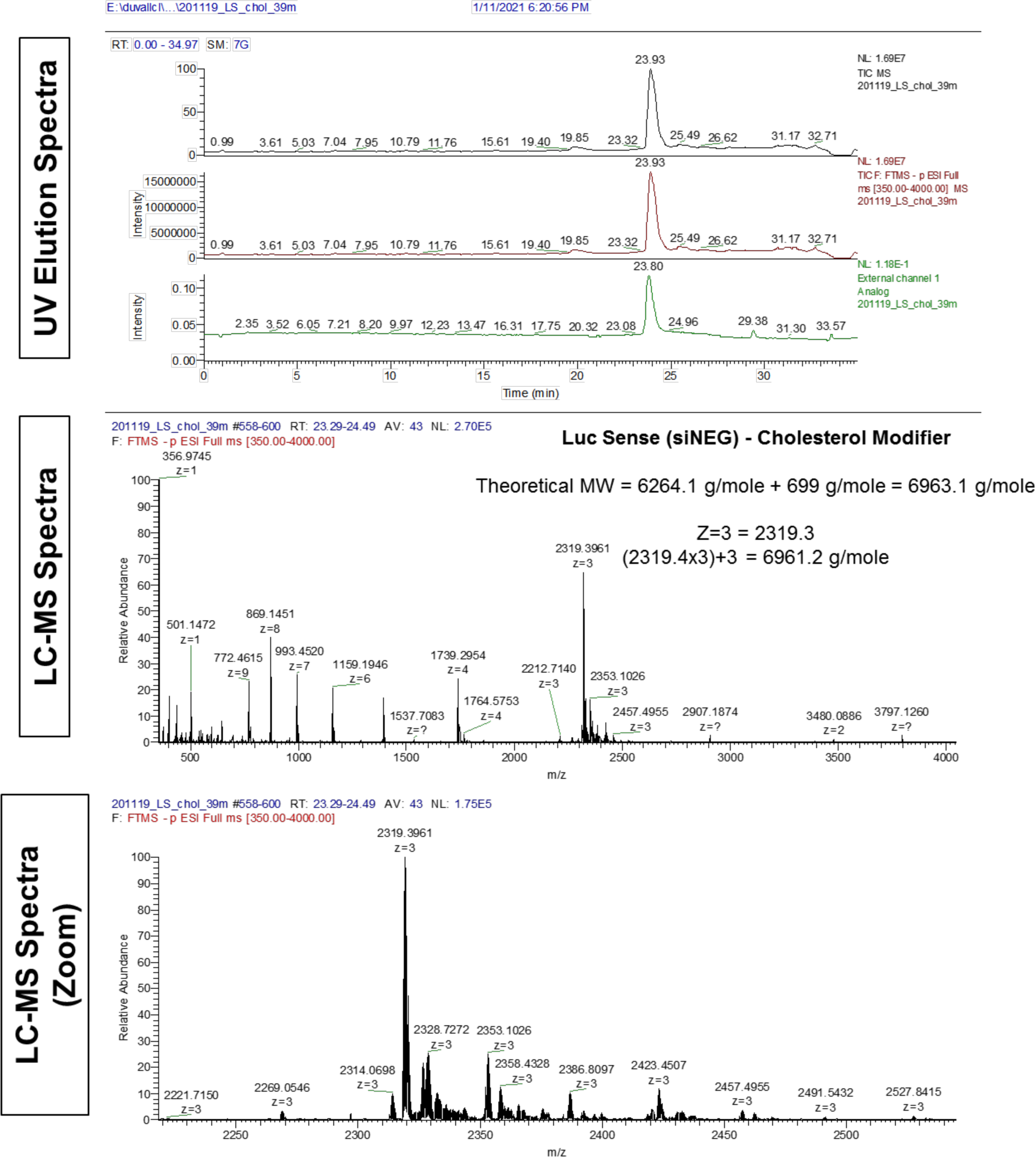
LC-MS characterization of control sense – cholesterol.

**Supplementary Figure 25.**
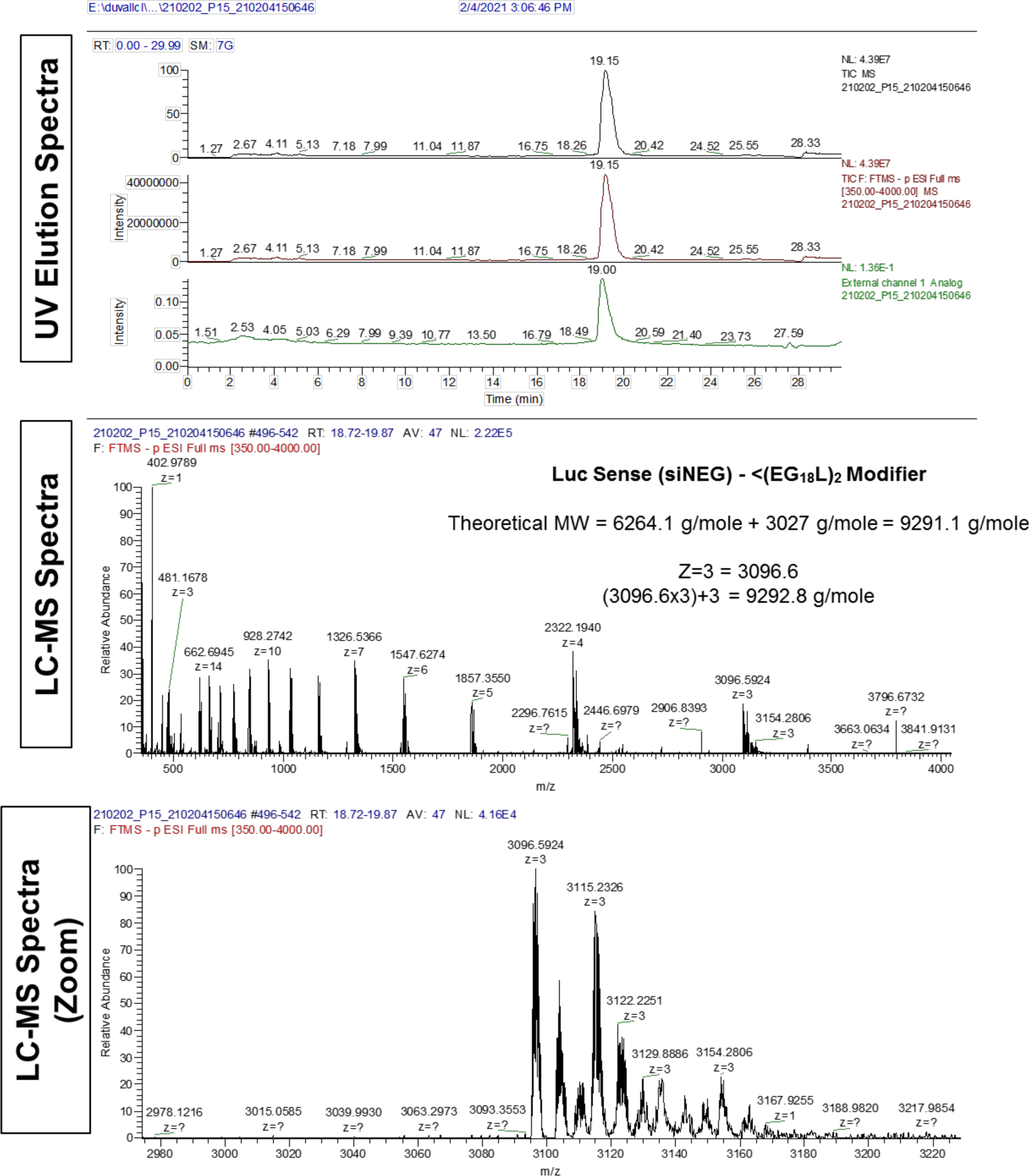
LC-MS characterization of control sense – <(EG_18_L)_2_.

**Supplementary Figure 26.**
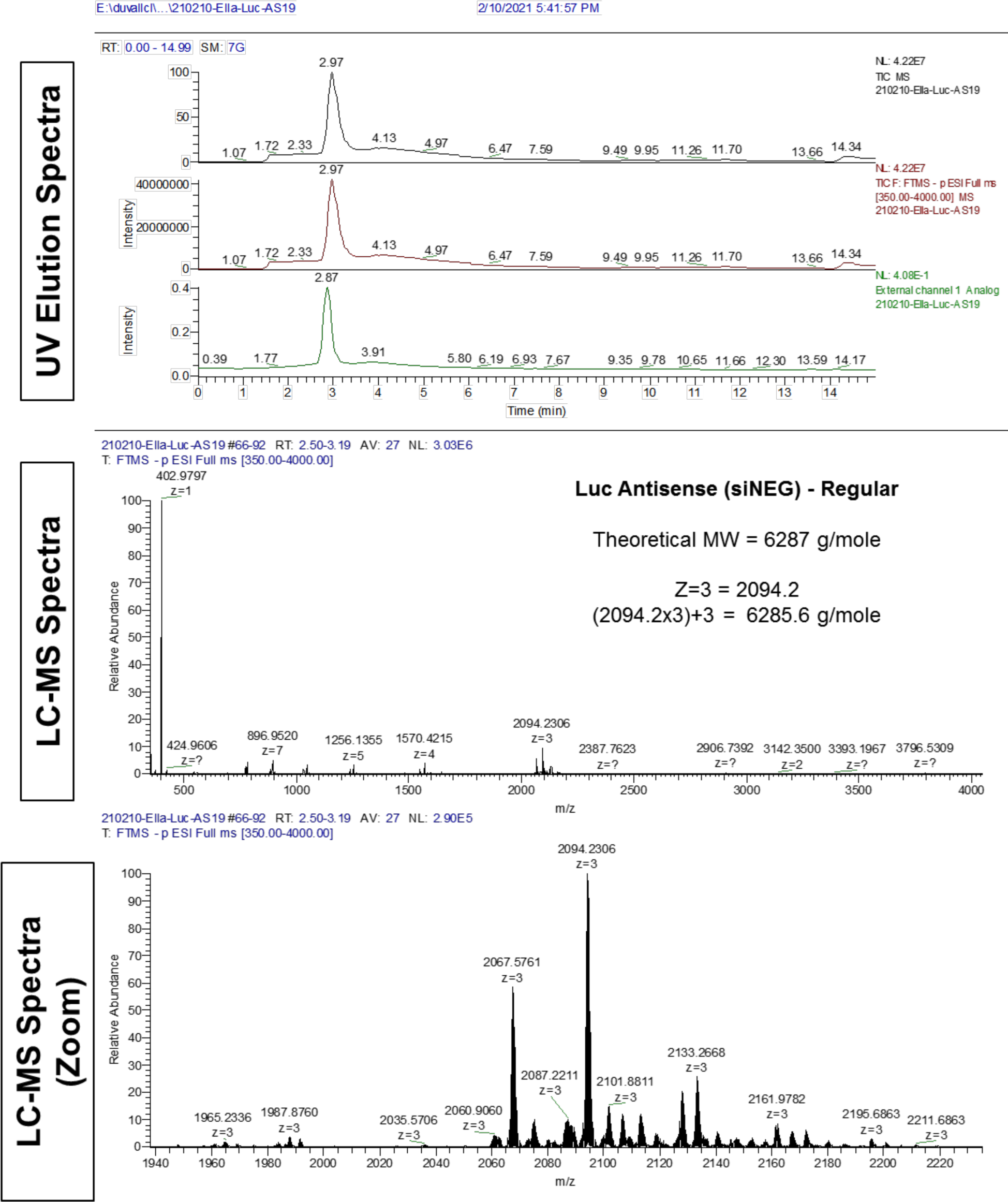
LC-MS characterization of control antisense – siRNA sequence.

**Supplementary Figure 27.**
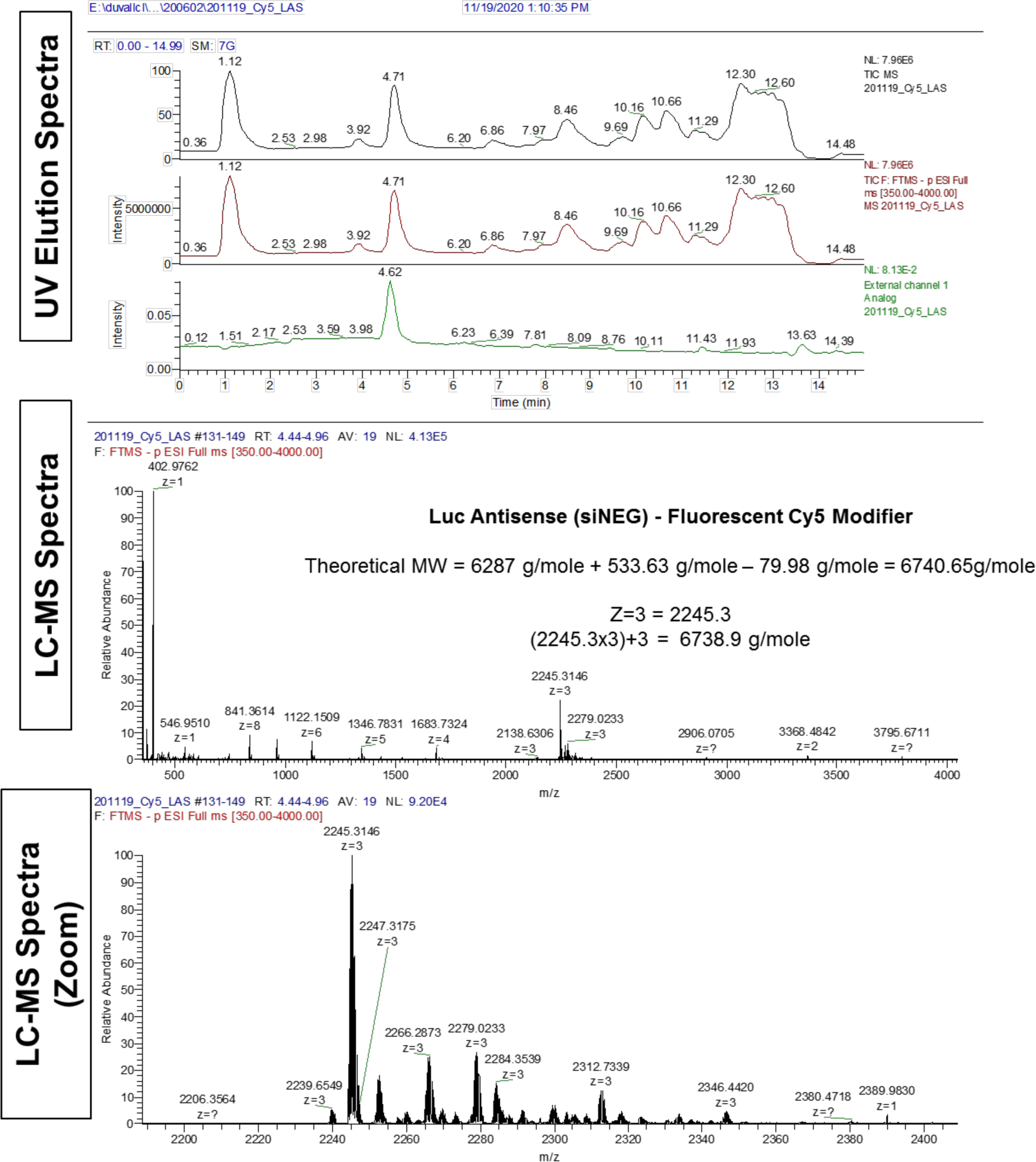
LC-MS characterization of control antisense – Cy5 modifier siRNA sequence.

**Supplementary Figure 28.**
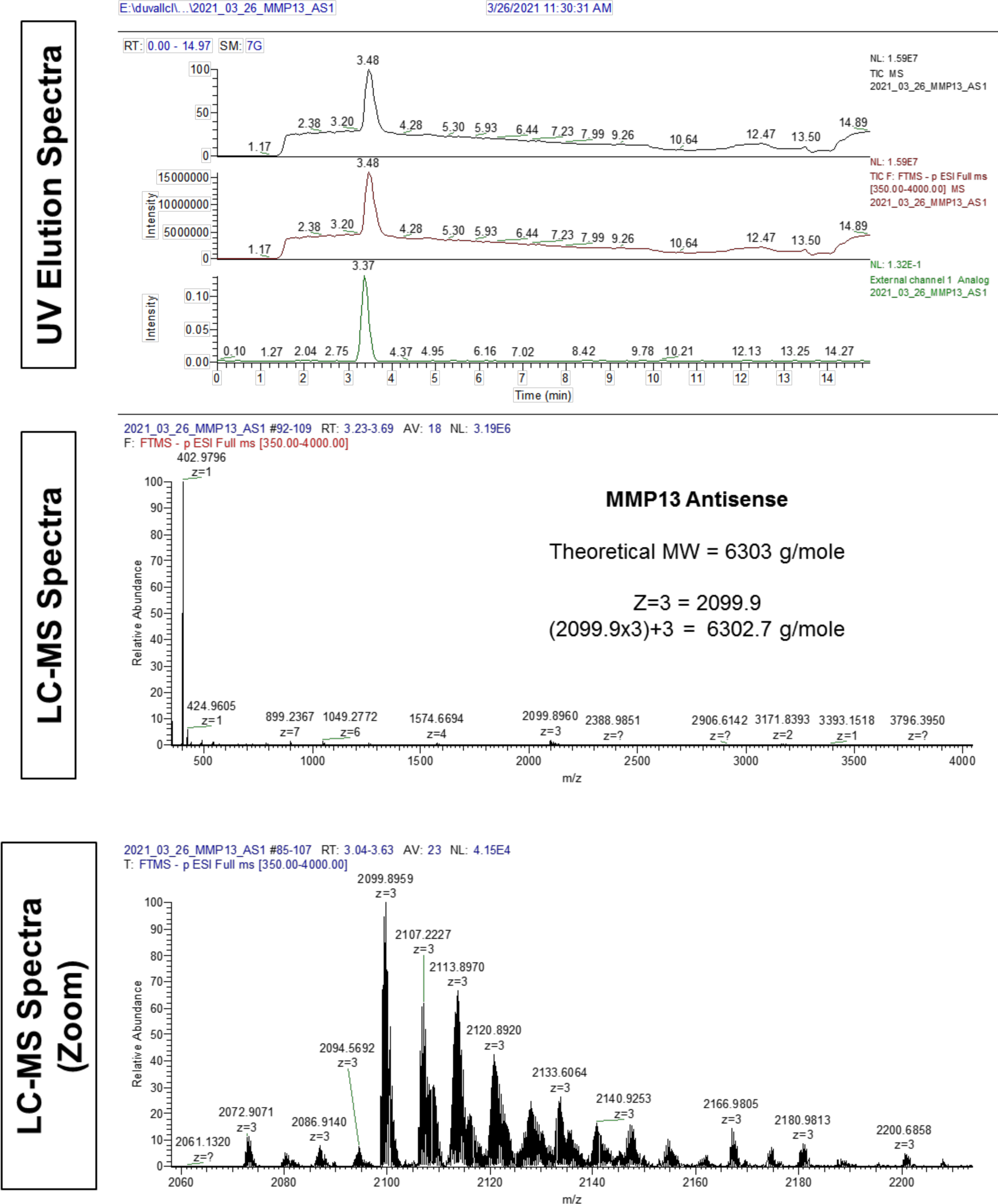
LC-MS characterization of mouse MMP13 antisense – siRNA sequence.

**Supplementary Figure 29.**
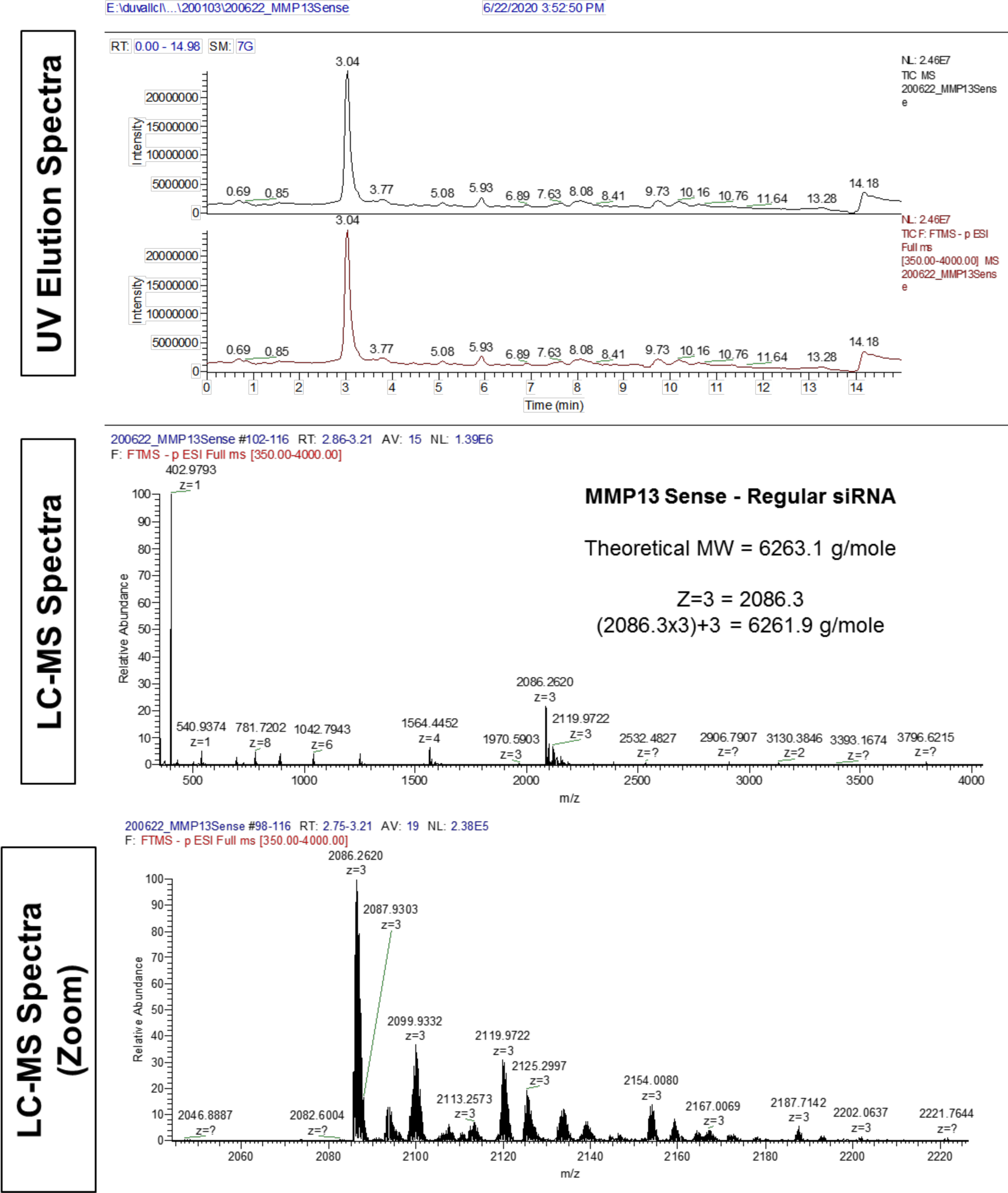
LC-MS characterization of mouse MMP13 sense – siRNA sequence.

**Supplementary Figure 30.**
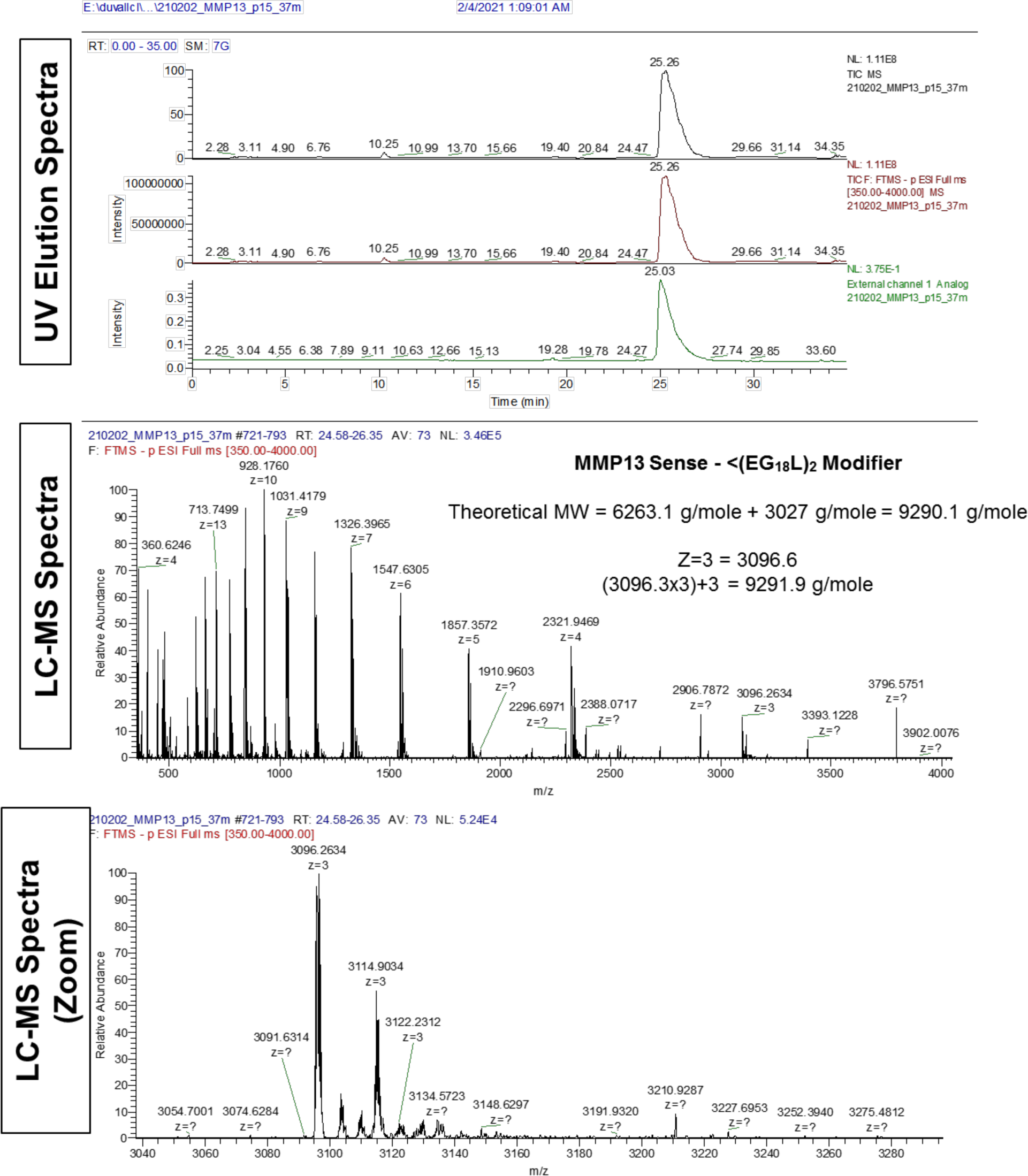
LC-MS characterization of mouse MMP13 sense – <(EG_18_L)_2_ modifier siRNA sequence.

**Supplementary Figure 31.**
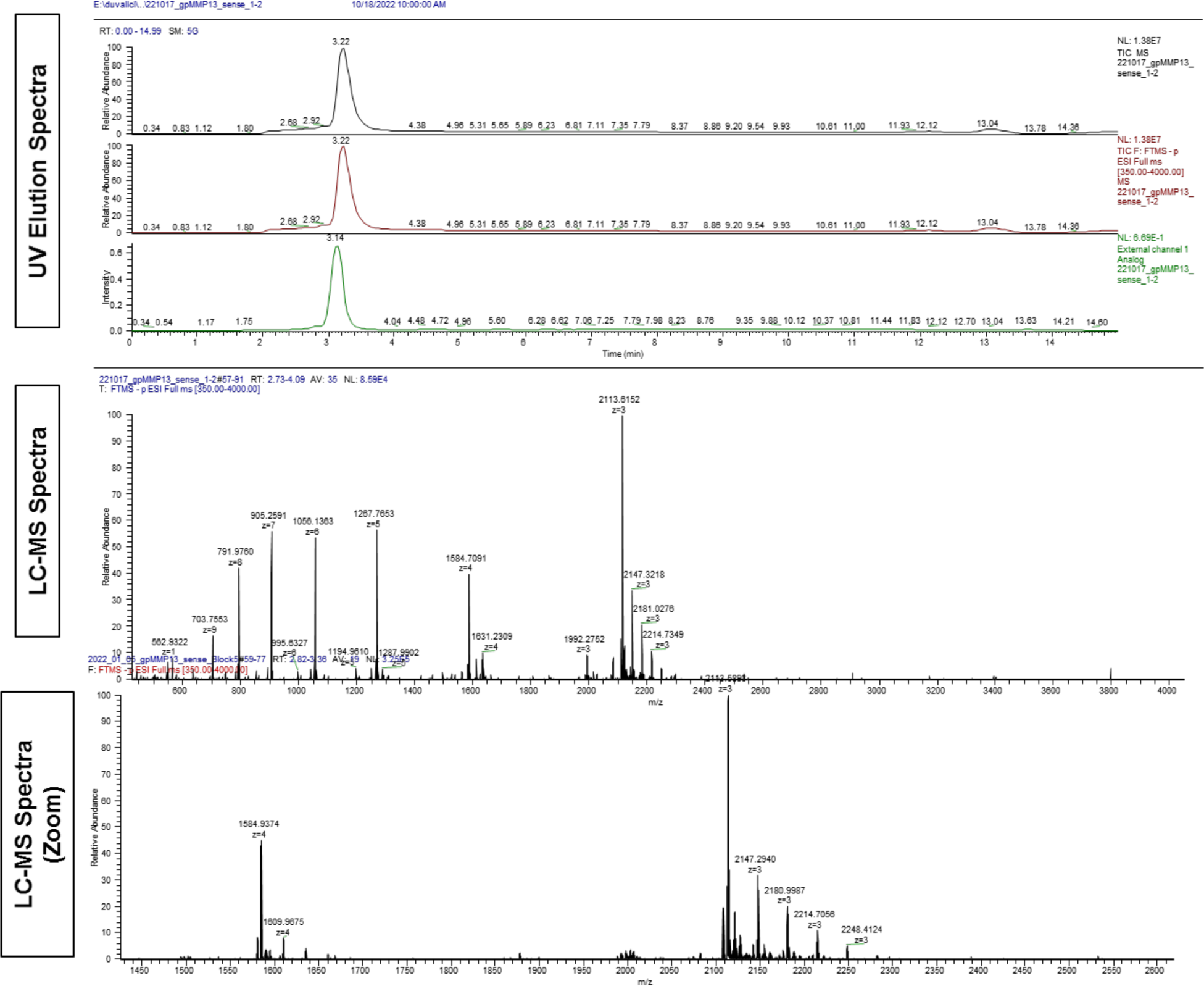
LC-MS characterization of guinea pig MMP13 sense siRNA sequence.

**Supplementary Figure 32.**
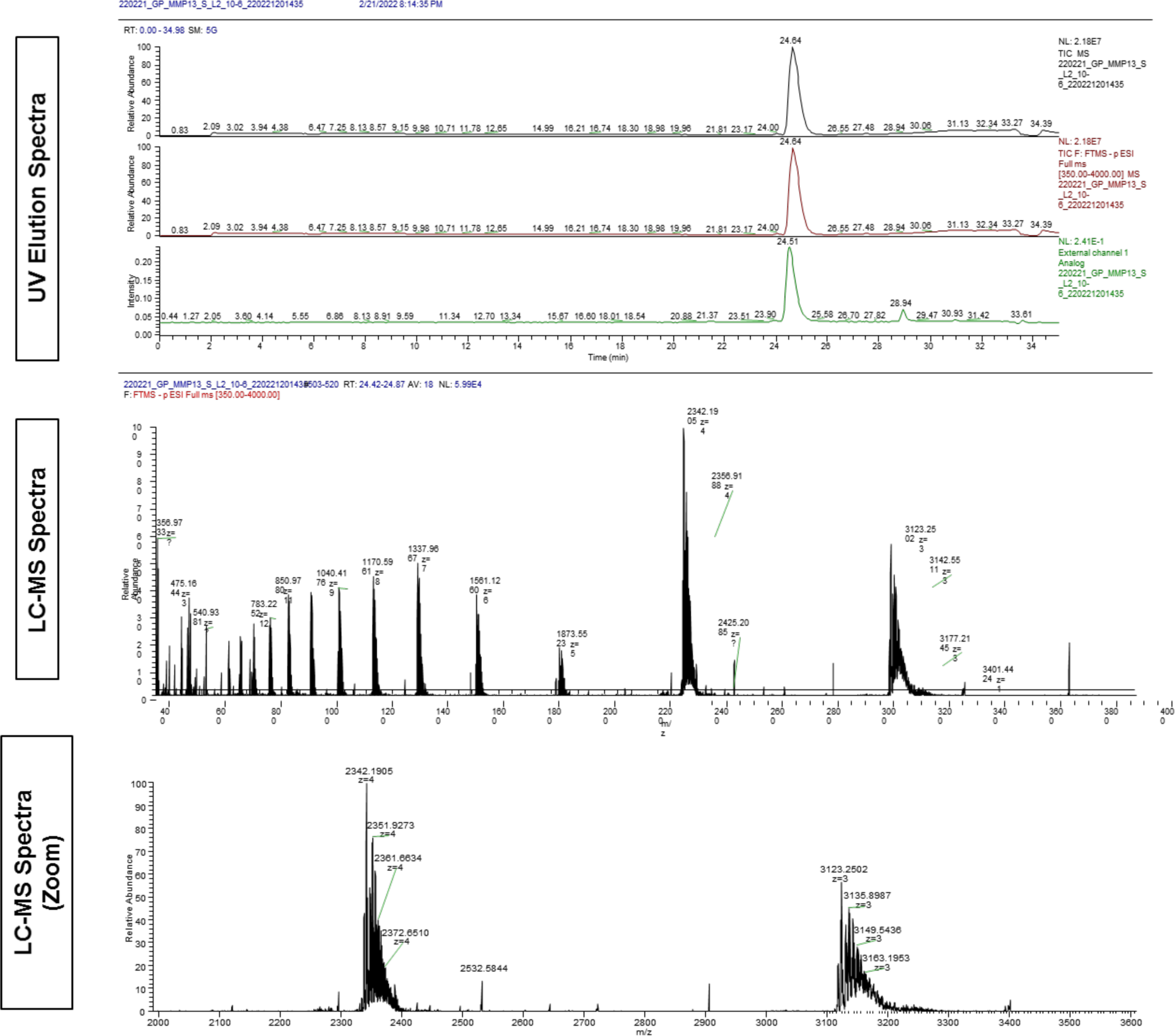
LC-MS characterization of guinea pig MMP13 sense – <(EG_18_L)_2_ modifier siRNA sequence.

**Supplementary Figure 33.**
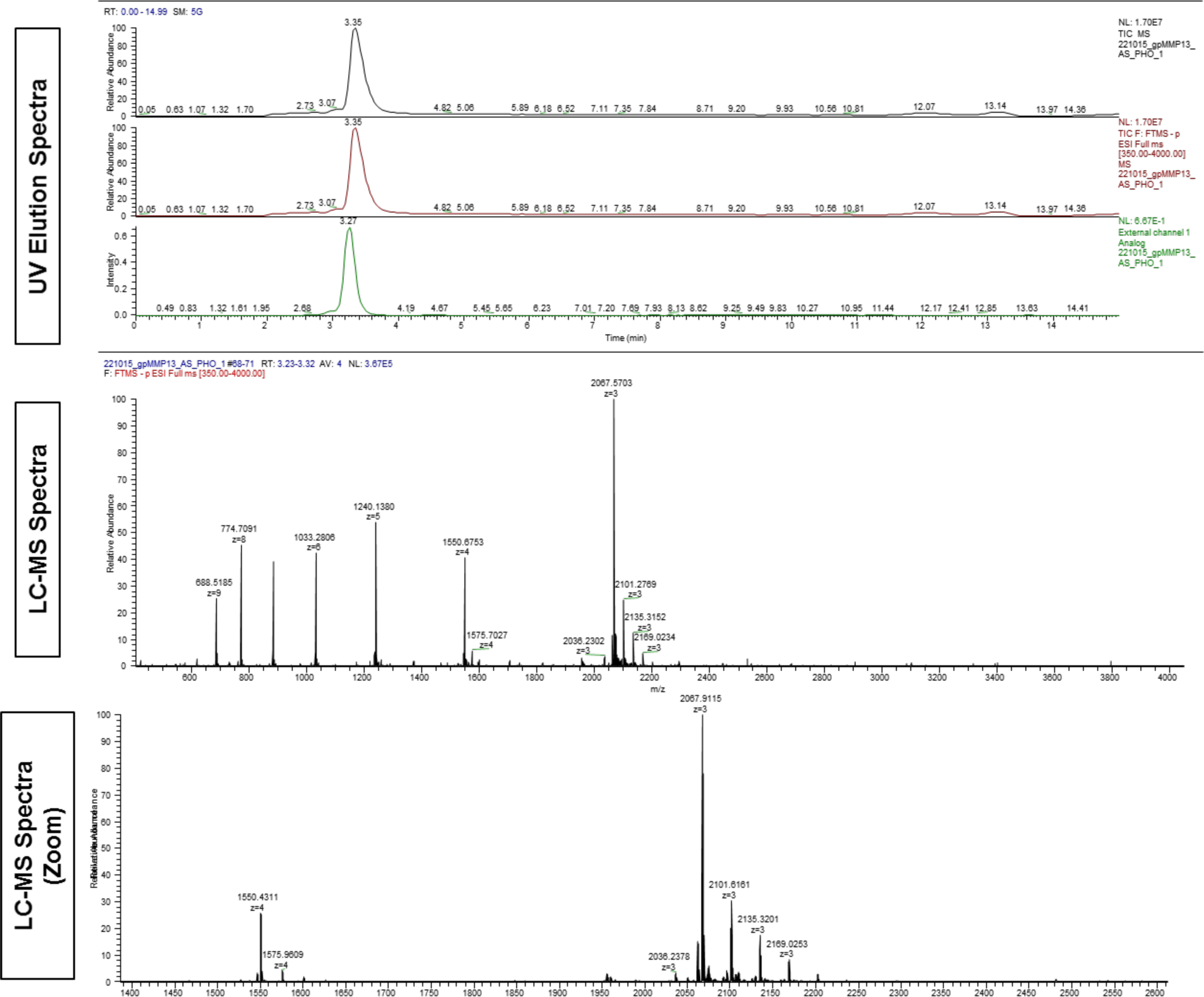
LC-MS Characterization of Guinea Pig MMP13 Antisense – siRNA Sequence.

