## Supplemental Methods for "Albumin-binding RNAi Conjugate for Carrier Free Treatment of Arthritis"

**Methods/Experimental Section**

**Materials.** 2’-O-Me and 2’-F phosphoramidites and universal synthesis columns (MM1-2500-1) were purchased from Bioautomation, symmetrical branching CED phosphoramidite from ChemGenes (CLP-5215), and cyanine 5 phosphoramidite (10-5915), stearyl phosphoramidite (10-1979), hexaethyleneglycol phosphoramidite (10-1918), TEG cholesterol phosphoramidite (10-1976), and desalting columns (60-5010) were from Glen Research. Unless otherwise stated, materials and reagents were purchased from Fisher Scientific (Waltham, MA, USA) or Sigma-Aldrich (St. Louis, MO, USA). CL-82198 was purchased from MedChemExpress (HY-100359). Marimastat was purchased from APExBio (A4049). Clinical formulations of Zilretta and methylprednisolone were purchased through the Vanderbilt University Medical Center pharmacy. C2C ELISA kit and C1,2C antibody was from IBEX Pharmaceuticals.

**siRNA synthesis, modification, annealing, and storage.** siRNA sequences were synthesized using modified (2’-F and 2’-O-Me) phosphoramidites with standard protecting groups on a MerMade 12 Oligonucleotide Synthesizer (Bioautomation). Amidites were dissolved at 0.1M in anhydrous acetonitrile, except for 2'OMe U-CE phosphoramidite which was dissolved in 20% (v:v) anhydrous dimethylformamide and stearyl phosphoramidite which was dissolved in 3:1 (v:v) dichloromethane:acetonitrile. Coupling was performed under standard conditions. Strands were grown on controlled pore glass with a universal terminus (1 µmol scale, 1000Å pore), cleaved and deprotected using 1:1 methylamine:40% ammonium hydroxide. Lipophilic RNAs were purified by reverse-phase (RP) high performance liquid chromatography (HPLC) using a Clarity Oligo-RP column (Phenomenex) under a linear gradient from 85% mobile phase A (50 mM triethylammonium acetate in water) to 100% mobile phase B (methanol) or 95% mobile phase A to 100% mobile phase B (acetonitrile). Oligonucleotide-containing fractions were dried, resuspended in nuclease free water, sterile filtered, and lyophilized. Conjugate molecular weight and purity was confirmed by Liquid Chromatography-Mass Spectrometry (LC-MS, ThermoFisher LTQ Orbitrap XL Linear Ion Trap Mass Spectrometer). Chromatography was performed using a Waters XBridge Oligonucleotide BEH C18 Column under a linear gradient from 85% A (16.3 mM triethylamine – 400 mM hexafluoroisopropanol) to 100% B (methanol) at 45°C. Mass spectrometry combined with liquid chromatography was utilized for product characterization **(Supplementary Figures 21-31)**. Purified oligonucleotide was resuspended in 0.9% sterile saline, annealed to the complementary strand by heating (95°C) and stepwise cooling (15°C / 9 min) to 25°C, and then stored at -80°C.

**MMP13 siRNA delivery to cultured cells.** siRNA sequences used are listed in **Supplementary Figure 9.** Seven candidate siRNA sequences (Dharmacon) targeting murine *Mmp13*, and 4 candidates (Dharmacon) targeting guinea pig *Mmp13*, were transfected (Lipofectamine 2000, Thermofisher) into ATDC5 cells or primary guinea pig chondrocytes (respectively) in OptiMEM for 24 hrs. Stabilized “zipper” siRNA sequences (50nM and 100nM, described below) were transfected into cells using Lipofectamine 2000 in OptiMEM for 24 hrs. For carrier-free delivery of siRNA and siRNA<(EG_18_L)_2_ to cells, siRNA (1000 nM OptiMEM) was added to cells for 24 hrs. In all cases, complete media supplemented with mouse or guinea pig tumor necrosis factor (TNF)-α (20 ng/ml) was added for the final 24 hrs. RNA was harvested, reverse transcribed, and assessed by real-time polymerase chain reaction (PCR) using primers against mouse or guinea pig *Mmp13*. CellTiter-Glo (Promega Corporation) assay was used to assess cytotoxicity.

**siRNA stability analysis.** siRNA (1 nmol) was incubated at 37° for 0-24 h in 60% fetal bovine serum (**FBS**) in PBS, or in patient-derived arthritic joint synovial fluid (83-year-old male patient with untreated, active osteoarthritis). Samples were resolved on 2% agarose gels. Gels were stained with GelRed Nucleic Acid Stain (Biotium) and imaged with UV transillumination.

**siRNA<(EG_18_L)_2_** **stability analysis (long-term).** siRNA<(EG_18_L)_2_ (2 nmol) was incubated at 37° for 0-96 hours in 5 µL patient-derived arthritic joint synovial fluid with rocking. Samples were resolved using 4-20% Native PAGE (Mini-Protean) with 250 ng in each lane. Gels were stained with GelRed Nucleic Acid Stain (Biotium) and imaged with UV transillumination.

**Cell culture.** Immortalized mouse chondrogenic ATDC5 cells (ATCC) were cultured in DMEM/F-12, GlutaMAX medium with 10% FBS and 1% penicillin/streptomycin (P/S). Primary guinea pig knee joint chondrocytes were cultured in DMEM with 10% FBS and 1% P/S. Cells were incubated at 37°C in 5% CO_2_. All cells tested negative for *Mycoplasma* using MycoAlert Mycoplasma Detection Kit (Lonza).

**Quantitation of siRNA uptake in cultured cells.** ATDC5 cells (15,000/well) were seeded overnight in 8-well chamber slides, treated with Cy5-labeled siRNAs pre-complexed with mouse serum albumin (MSA, 10x molar excess) for 4 hrs in Opti-MEM, washed with PBS, fixed and counterstained with DAPI. Cy5 imaging was performed by confocal microscopy on a Nikon Eclipse Ti-0E inverted microscopy base. Alternatively, ATDC5 cells (10,000/well) in 96-well plates were seeded overnight, treated 2 hrs with Cy5-labeled siRNA (100 nM) with or without 1 μM MSA or 10% serum, in OptiMEM. Cells were collected by trypsinization and assessed by flow cytometry on a Guava EasyCyte (Luminex), gating >500 cellular events.

**Size exclusion chromatography (SEC).** 1 μM of Cy5-labeled siRNA molecules were incubated 30 min with 100 mL patient-derived synovial fluid collected from joints of a non-arthritic donor (myocardial infarction death, no history of rheumatic disease), a donor with untreated osteoarthritis, or a donor with untreated rheumatoid arthritis. The siRNA-synovial fluid mixtures were filtered (0.22 µm) and injected into AKTA Pure Chromatography System (Cytiva) for fractionation with three inline Superdex 200 Increase columns (10/300 GL) at 0.3 mL/min using 10mM Tris-HCl, 0.15M NaCl, 0.2% NaN_3_. 1.5 mL fractions were collected (F9-C 96-well plate fraction collector, Cytiva). Cy5 fluorescence was measured in fractions (100 µL) in black walled 96-well plates on a SynergyMx (Biotek). Albumin-associated fractions were determined empirically using known protein standards, examining A280 of eluent from each fraction. Fractions were resolved on 4-20% SDS-PAGE (Mini-Protean), transferred to nitrocellulose (iBlot2, Invitrogen), and stained with Coomassie Blue or probed with anti-human albumin antibody (ab19180, Abcam) and IRDye^®^ 800CW Donkey anti-Goat IgG, (Li-Cor).

**Mouse models of arthritis.** A mechanical overload model of post-traumatic osteoarthritis was used, using an ElectroForce 3100 test frame (TA Instruments). Knee joints of anesthetized, 6 week old C57BL/6 male mice were positioned in flexion at 140° with the tibia approximately vertical and placed directly under the loading point ^1^. Repetitive knee loading was performed with 250 cycles of 9 N of compressive mechanical force as described in previous studies ^1,2^. Loading was repeated three times weekly for up to 5 weeks in either the left knee only, or in both knees, as indicated. For modeling rheumatoid arthritis, serum (200 µl) collected from donor K/BxN transgenic mice was transferred by intraperitoneal delivery to 8-week-old male C57BL/6 mice recipients ^3^.

**Cy5 labeling.** MSA (Sigma) and 40kDa PEG (Sigma) were Cy5-labelled [Cy5® Conjugation Kit, (Abcam)] and purified, confirming Cy5 fluorescence by plate fluorimeter (Tecan).

**Pharmacokinetic and retention studies.** For pharmacokinetics, Cy5-siRNA, Cy5-siChol, Cy5-siRNA<(EG_0_L)_2_, Cy5-siRNA<(EG_6_L)_2_, Cy5-siRNA<(EG_18_L)_2_, Cy5-siRNA<(EG_30_L)_2_ molecules were delivered i.v. (1 mg/kg), subcutaneously (2 mg/kg), or intra-articular (0.25 mg/kg). Cy5-MSA (3mg/kg) and Cy5-PEG (3mg/kg) were delivered i.v. For retention studies, Cy5-siRNA<(EG_18_L)_2_ was delivered intravenously (10 mg/kg), or by intra-articular injection (1 mg/kg). Mouse hindlimbs were depilated prior to intravital Cy5 imaging (IVIS Lumina III, Caliper Life Sciences) at indicated timepoints. Ex vivo Cy5 imaging of organs and denuded limbs collected at necropsy was performed. Where indicated, knee joints were micro-dissected for Cy5 imaging of knee cartilage and synovial tissues. For Cy5 imaging in histological sections, hindlimbs were embedded into OCT freezing compound, serially sectioned at various depths along the joint at 20 µm, captured onto polyvinylidene chloride film coated with synthetic rubber cement (<http://section-lab.jp/>), placed on slides, formalin fixed, and counterstained with DAPI (in some cases). Cy5 fluorescence was imaged using a Nikon Eclipse Ti inverted confocal microscope. Whole joint imaging was performed by stitching 4 x 4 10x images.

**Cellular uptake assessment via flow cytometry.** Cy5-siRNA<(EG_18_L)_2_ (10 mg/kg) or vehicle (.9% NaCl) was delivered i.v. to mice having undergone one week of bilateral loading as described above (**Supplementary Figure 8A**). Mice were sacrificed 24 hours after injection and synovial tissue was isolated from the anterior, medial, and lateral compartments as previously described^4^. The posterior synovium was not collected. Two synovia were digested together in a volume of 1.5 mL digestion media (DMEM with 400 μg/mL collagenase IV, liberase, and DNaseI). Synovia were digested for 40 minutes at 37°C with intermittent vortexing at 0, 15, 30, 35 and 40 minutes, and then pelleted at 500 x g for 5 minutes at 4°C. Each treatment group (vehicle and Cy5-siRNA<(EG_18_L)_2_) was comprised of 5 male C57/B6 mice, which were pooled and divided into the controls and samples. Cold FACS buffer (PBS containing 1% FBS, 2 mM EDTA) was used for all antibody staining and wash steps. Non-specific binding was blocked using Mouse Seroblock FcR (Bio-Rad) for 5 minutes. CD45-PerCp/Cy5.5 (Biolegend), FAP-AF488 (R&D Systems), and CD31-PE (Invitrogen) were used to identify major synovial cell populations. CD3-SBV515 (Bio-Rad), CD11b-APC/Cy7 (Biolegend), F4/80-PE/Cy7 (Biolegend) and CD11c-BV605 (Biolegend) were used to identify specific immune populations. All cell populations were gated off fluorescence-minus-one (FMO) controls **Supplementary Figure 8B**. The Cy5 gate was established using the vehicle mouse sample and applied to each cell population (**Supplementary Figure 8C**) and a %Cy5 positive proportion (**Figure 2F**) and median fluorescence intensity (MFI) (**Supplementary Figure 8D**) was calculated for all synovial cells and each cell type separately. Experiments included single-stained compensation controls to account for fluorescent bleed over. Acquisition was performed using a BD LSRFortessa cytometer with FACSDiva software (BD Biosciences), then data compensation and analysis was performed using FlowJo v10 (TreeStar/BD Biosciences). Antibody details are listed in **Supplementary Figure 8E**.

**Peptide nucleic acid (PNA) hybridization assay.** Quantification of antisense strands in tissue was performed using a PNA hybridization assay as previously described^5,6^. In brief, knee joints were dissected into mixed cartilage/meniscus and synovial tissues (5-15 mg) and lysed in 300 μL QuantiGene™ homogenizing solution (Invitrogen) containing 0.2 mg/mL Proteinase K (Invitrogen). Sodium dodecyl sulfate (SDS) was precipitated from homogenates with 20 μL 3M potassium chloride and centrifuged at 4,000 × *g* for 15 min. To anneal the PNA probe, 100 μL of hybridization buffer (50 mM Tris, 10% ACN, pH 8.8) and 5 pmol of Cy3-labeled probe (PNABio) were added to 150 μL of each homogenate. Samples were then incubated for 15 min at 90°C and then 50 °C before analysis by ion exchange on an iSeries LC equipped with RF-20A fluorescence detector (Shimadzu) over a DNAPac PA100 anion-exchange column (Thermo Fisher Scientific). Mobile phases consisted of buffer A (50% acetonitrile and 50% 25 mM Tris–HCl, pH 8.5; 1 mM ethylenediaminetetraacetate in water) and buffer B (800 mM sodium perchlorate in buffer A) and a gradient was obtained as follows: 10% buffer B for 4 min, 50% buffer B for 1 min and 50% to 100% buffer B within 5 min. The final mass of siRNA was calculated from a standard curve of known quantities of siRNA or EG18 spiked into untreated tissue homogenates.

**Therapeutic treatment of arthritis models.** Zipper-modified siRNA targeting mouse MMP13 (siMMP13) or in non-targeting siRNA (siControl) was delivered to mice at either 5 mg/kg or 10 mg/kg i.v., 1 mg/kg intra-articular, or 20-50 mg/kg subcutaneous. Mice were treated in parallel studies with Marimastat (10 mg/kg, i.p.), Cl-82198 (10mg/kg i.p.), methylprednisolone (10 mg/kg i.p.), or Zilretta (8 mg/kg)**^7^**.

**In situ imaging of MMP activity and collagen fragments.** Mice were injected with mAbCII-680 (50uL of 1 mg/ml, trail vein) or with MMPsense750 fast (100uL of resuspended formulation per instructions, tail vein, NEV10168, PerkinElmer, Waltham, Massachusetts, USA). Hindlimbs were depilated for IVIS imaging of fluorescence intensity.

**Gene expression analysis in animal tissues.** At necropsy, hindlimbs were cleaned of excess muscle and placed in RNAlater solution (ThermoFisher Scientific). Anterior synovial tissue pads, menisci, and tibial and femoral articular surface cartilage specimens were collected by microdissection. Cartilage, meniscal tissue, and synovial tissue were admixed in equal mass ratios and termed ‘whole joint’ and used for mRNA extraction (RNeasy Plus Mini Kit from Qiagen). RNA from whole joint, liver, or kidney was reverse transcribed (iScript cDNA Synthesis Kit, Bio-Rad) and used for real-time PCR with the following: murine Taqman probes: **[***Actb* (Mm02619580_g1); *Gapdh* (Mm99999915_g1); Mmp13 (Mm00439491_m1); *Il1b* (Mm00434228_m1)**;** *Il6* (Mm00446190_m1); *Cox2* (Mm03294838_g1); *Tnf* (Mm00443258_m1); *Sparc*: ([Mm00486332_m1](https://www.thermofisher.com/taqman-gene-expression/product/Mm00486332_m1?CID=&ICID=&subtype=)); *Cav1* ([Mm00483057_m1](https://www.thermofisher.com/taqman-gene-expression/product/Mm00483057_m1?CID=&ICID=&subtype=)); *Fcrn* (Mm00438887_m1); *Ngf* (Mm00443039_m1); *Cdkn1b* (Mm00494449_m1)]. Guinea Pig Taqman Probes: [*Actb* (Cp03755210_g1); *Gapdh* (Cp03755743_g1)**;** *Mmp13* (APRWJ74)]. RNA was processed for nanoString analysis using the mouse nCounter Inflammation Panel by the Vanderbilt VANTAGE shared resource, hybridizing for 20 hrs per manufacturer directions (Nanostring Technologies, Inc.). Data analysis was performed using nSolver software for comparison, unsupervised analysis, and gene cluster analysis between groups.

**Mining of single-cell RNA-sequencing data for MMP13 expression in synovium.** Single-cell RNA-sequencing (scRNAseq) data from Knights et al, Ann Rheum Dis 2023 (NIH GEO Accession: GSE211584)^4^ was employed to identify *Mmp13-*expressing cells in murine PTOA synovium. Data import, quality control, dimensionality reduction, and clustering was performed as described in the original publication to yield a cellular atlas of healthy and PTOA synovium, comprised of lining fibroblasts, sublining fibroblasts, myeloid cells, dendritic cells, T cells, pericytes, endothelial cells, lymphatic endothelial cells, Schwann cells, skeletal muscle, and red blood cells. Healthy synovium represented cells from the “Sham” condition (no injury loading, analgesia and anesthesia only) and PTOA synovium represented merged cells from the “ACLR 7d” and “ACLR 28d” conditions, which underwent noninvasive anterior cruciate ligament rupture via mechanical loading. All conditions were made up of pooled male and female mice, with ~20,500 total cells in the analysis. Gene feature plots of normalized *Mmp13* transcript expression were derived to demonstrate the cellular sources and PTOA-associated induction of *Mmp13* in murine synovium.

**Evans blue delivery and extraction.** Evans Blue (200 μL of 2% wt/v in sterile saline) was delivered by intravenous injection. After 24 hrs, mice were perfused with PBS. Hindlimbs collected at necropsy were air dried overnight. Joint tissues were micro-dissected. Evans Blue was extracted from dissected tissues in formamide at 55 ºC for 24 hrs. Evans Blue extracted from tissue was measured as absorbance at 610 nm.

**Immunofluorescence Staining.** 20 µm cryosections were fixed 10 min with 4% PFA, blocked in 5% donkey serum, and probed with the following: anti-guinea pig MMP13 (1:100, ARP56350_P050, Aviva Systems Biology); goat anti‐rabbit Alexa Fluor® 488 (1:500, ab150077, Abcam); Lycopersicon Esculentum (Tomato) Lectin (LEL, TL) DyLight™ 488 (1:100, DL-1174-1). Slides were counterstained with DAPI, mounted with ProLong Gold Antifade, and imaged on a Nikon Eclipse Ti inverted confocal microscope. Imaging settings remained constant across all treatment groups.

**Staining of paraffin embedded histological sections.** Joints were formalin fixed, decalcified (20% tetrasodium EDTA), and paraffin embedded. Formalin fixed paraffin embedded coronal (knee) or sagittal (paw) sections (5µm) were stained with hematoxylin and eosin, toluidine blue, or used for antigen retrieval in Epitope Retrieval 1 (Bond Rx) and immunohistochemistry. The following antibodies were used: anti-MMP13 (1:750, Abcam Ab39012), C1,2C (Col 2 3/4Cshort) Polyclonal Rabbit Antibody (1:500, IBEX Pharmaceuticals 50-1035). The Bond Refine Polymer detection system was used for visualization. Slides were imaged on the Leica SCN400 Slide Scanner (Leica Biosystems Inc.).

| **Severity Score** | **OARSI**  **Scale** | **Degenerative Joint Disease (DJD)**  **Scale** | |
| --- | --- | --- | --- |
| 0 | Normal | **Within normal**  **Limits** | mild DJD as a feature of expected age-related change (e.g., attenuation of articular cartilage and proteoglycan loss) |
| 1 | Loss of SO staining, articular cartilage thinning; no defects | **Moderate** | - articular cartilage degeneration - beginning of secondary pathology - synovitis - joint capsule fibrosis - meniscal metaplasia |
| 2 | 1 + fibrillation or pyknotic articular chondrocytes | **Marked** | - metaplasia and/or fragmentation of one meniscus - osteophyte formation - synovitis or hyperplasia |
| 3 | 2 + loss of articular cartilage <50% (e.g., erosion, flap, or callus) | **Severe** | - metaplasia and/or fragmentation of both menisci - eburnation (total loss) of at least 1 articular surface - osteophyte formation - advanced synovitis, hyperplasia |
| 4 | 3 + fragmentation and fissuring in 1+ articular plateau | **Extreme** | - most advanced from severe category - epiphyseal osteolysis - collapse of joint space |
| 5 | 4 + fragmentation & fissuring of >75% cartilage surface | **N/A** | |
| 6 | Total loss of normal articular cartilage (end-stage) | **N/A** | |

**Supplementary Table 1. (A) Description of OARSI scale for scoring Safranin-O/Fast Green stained slides of femoral/tibial cartilage plateaus (left); (B) Criteria for scoring of H&E-stained slides to assess overall joint by the DJD scale (right).**

**Clinical and histological scoring of arthritis.** Knee joint hyperalgesia was quantified using a hand-held algesiometer (Bioseb SMALGO: SMall animal ALGOmeter). For measurements, the pressure applicator tip was positioned on the medial aspect of the knee joint and forced increased until eliciting a nociceptive response. Each limb was assessed in triplicate. The clinical score of arthritis was assessed as swelling / erythema on a scale of 0 to 3 (0, none; 1, slight; 2, moderate and in multiple digits; 3, pronounced and in entire paw)**^8^**. Ankle thickness was measured with calipers. Serum specimens were assessed for collagen degradation fragments using C2C enzyme-linked immunosorbent assay kit (IBEX Pharmaceuticals). Stained coronal knee joint sections were scored from at least 2 mid-frontal coronal sections per joint^9^ using the Osteoarthritis Research Society International (OARSI) and Degenerative Joint Disease (DJD) scales by a board-certified veterinary pathologist. OARSI scores (0-6 semiquantitative scale; **Supplementary Table 1**) were assigned from assessment of the medial and lateral plateaus of the tibia and femur ^10^. DJD severity score (0-4 semiquantitative scale) was determined from H&E-stained sections using a semi-quantitative index (**Supplementary Table 1**) based on cartilage erosion, subchondral osteosclerosis, synovial/meniscal metaplasia, subchondral osteosclerosis, inflammation, osteophytes, and meniscal ectopic mineral deposits^11^. For the K/BxN STA study, hindpaws and forepaws were sectioned sagitally with knee joints being sectioned coronally; they were also stained with Hematoxylin and Eosin and Toluidine Blue for scoring purposes. Toluidine Blue staining was verified by visualization of mouse skin mast cells and mouse ear cartilage. For the K/BxN serum transfer arthritis model, forepaw, hindpaw, and knee joint sections were also stained with Hematoxylin and Eosin and Toluidine Blue and were scored using 3 scales adapted as previously described^8,12^. Scoring was performed by a histopathologist blinded to the treatment.

| **Inflammation score (H&E)** | |
| --- | --- |
| 0 | Absent |
| 1 | Slight/mild inflammation (diffusely located single cells and/or small perivascular infiltrates of neutrophils with lymphocytes and plasma cells) |
| 2 | Moderate inflammation (moderate, multifocal infiltrates of neutrophils with lymphocytes and plasma cells) |
| 3 | Marked inflammation (diffuse, confluent infiltrates of neutrophils with lymphocytes and plasma cells) |
| **Bone erosion score (H&E)** | |
| 0 | None/normal |
| 1 | Minimal/sparse (small areas of resorption, hard to identify) |
| 2 | Mild (mild but multifocal resorption of cortical and/or trabecular bone) |
| 3 | Moderate (resorption of cortical and/or trabecular bone; not full thickness) |
| 4 | Marked (full thickness defects in the cortical bone with marked trabecular bone loss) |
| **Cartilage destruction score (Toluidine Blue)** | |
| 0 | None/normal |
| 1 | 0-25% staining loss |
| 2 | 25-50% staining loss |
| 3 | 50-75% staining loss |
| 4 | 75-100% staining loss |

**MicroCT.** Joints were fixed with formalin and submerged in 100% ethanol during micro-CT imaging with the ScanCo μCT-50 (Scanco Medical) with reconstructions completed in the ScanCo software (Scanco USA). Images were collected with 20 μm thick slices, isotropic 12 μm voxel at 114 mA / 70 kVp, and 200 msec integration time. Initial contouring encompassed all mineralized joint components. Secondary contouring segmented out mineralization in the soft tissues surrounding cortical bone. 3D renderings (sigma: 1.5; support: 3; threshold: 388) were generated using Scanco software and are shown at a consistent density threshold (42.0% of maximum bone density, or 420 per mille). Imaging, contouring, and sample measurements were acquired by a treatment-blinded user. Osteophyte length was measured on each sample in femur and tibia. For K/BxN serum recipients, contouring was performed over the calcaneus. Hydroxyapatite calibration phantoms were used to calibrate bone density values (g/cm^3^).

**Guinea pig anterior cruciate ligament (ACLT) model**. 3-month-old male Dunkin Hartley guinea pigs (Charles River Laboratories) underwent left knee ACLT surgery using procedures adapted from Jimenez et al. **^13^**. Surgery was performed under direct visualization in anesthetized animals. A medial longitudinal parapatellar incision over the anterior knee exposed the patellar tendon. The patella was everted, knee placed in flexion, and ACL incised. After confirming anterior joint laxity, the site was closed with the joint capsule continuously sutured. Ketoprofen was given immediately before and every 24 hours post-surgery for 3 days.

**Blood chemistry.** Whole blood was collected in EDTA-coated tubes or spun down at 1,000 x g for 10 min at 4°C to isolate serum. Samples were then submitted to the Vanderbilt Translational Pathology Shared Resource for chemistry analyses.

**Animal ethics statement**. All animal experiments described herein were carried out according to protocols approved by Vanderbilt University’s Institutional Animal Care and Use Committee (IACUC), and all studies followed the National Institutes of Health’s guidelines for the care and use of laboratory animals.

**Statistical methods**. Data are displayed as mean plus/minus standard deviation (unless stated otherwise). Statistical tests employed either one-way or two-way ANOVAs with multiple comparisons test or two-tailed student’s T-test between only two groups with α = 0.05 unless otherwise indicated. All statistical analysis were performed as described in figure captions with GraphPad Prism software, except for nanoString data analysis with nSolver V3.0 (Nanostring Technologies).
